# Altered mRNA transport and local translation in iNeurons with RNA binding protein knockdown

**DOI:** 10.1101/2024.09.26.615153

**Authors:** Rachael Dargan, Alla Mikheenko, Nicholas L. Johnson, Benjamin Packer, Ziyi Li, Emma J. Craig, Stephanie L. Sarbanes, Colleen Bereda, Puja R. Mehta, Matthew Keuss, Mike A. Nalls, Yue A. Qi, Cory A. Weller, Pietro Fratta, Veronica H. Ryan

## Abstract

Neurons rely on mRNA transport and local translation to facilitate rapid protein synthesis in processes far from the cell body. These processes allow precise spatial and temporal control of translation and are mediated by RNA binding proteins (RBPs), including those known to be associated with neurodegenerative diseases. Here, we use proteomics, transcriptomics, and microscopy to investigate the impact of RBP knockdown on mRNA transport and local translation in iPSC-derived neurons. We find thousands of transcripts enriched in neurites and that many of these transcripts are locally translated, possibly due to the shorter length of transcripts in neurites. Loss of frontotemporal dementia/amyotrophic lateral sclerosis (FTD/ALS)-associated RBPs TDP- 43 and hnRNPA1 lead to distinct alterations in the neuritic proteome and transcriptome. TDP-43 knockdown (KD) leads to increased neuritic mRNA and translation. In contrast, hnRNPA1 leads to increased neuritic mRNA, but not translation, and more moderate effects on local mRNA profiles, possibly due to compensation by hnRNPA3. These results highlight the crucial role of FTD/ALS-associated RBPs in mRNA transport and local translation in neurons and the importance of these processes in neuron health and disease.

## Introduction

Due to their long axons and need to respond to stimuli in a fraction of a second, neurons rely on transport of mRNAs to axons and dendrites to rapidly and efficiently synthesize new proteins at distal sites. This strict spatial and temporal control of translation is regulated in part by RNA-binding proteins (RBPs). RBPs bind to mRNAs early in their life cycle and participate in splicing, stabilization, transport, and translation of those transcripts (1,2). To date, most studies have focused on the localization of specific transcripts of interest (3–5), rather than looking at transcript localization in an unbiased, whole transcriptome manner. A recent study identified the local transcriptome and translatome in mouse embryonic stem cell-derived neurons and found that neurite targeted mRNAs encode approximately half of the neurite localized proteome (6). However, similar studies have not been performed in human neurons.

Mutations in RBPs have been identified in patients with familial forms of frontotemporal dementia/amyotrophic lateral sclerosis (FTD/ALS) (7). Even without mutations, approximately 40- 50% of FTD cases and >90% of ALS cases have inclusions of one RBP, TDP-43 (8). Occasionally, these inclusions can sequester other RBPs, potentially leading to decreased levels of many RBPs in nuclei (9). Loss of nuclear RBPs can induce splicing changes and mutant RBPs may have altered mRNA binding properties (10). Loss of TDP-43 is well known to induce splicing changes leading to inclusion of “cryptic exons” that are normally intronic and spliced out and can lead to either decreased protein levels or aberrant translation of the cryptic exon, further stressing the cell (10). Further, although some FTD/ALS-associated RBP mutations have been shown to alter mRNA transport of specific transcripts (11–13), the role of loss-of-function of FTD/ALS-associated RBPs on mRNA transport and local translation has not been evaluated. Here, we use proteomics, transcriptomics, and microscopy to obtain a comprehensive picture of the effect of loss of TDP-43 and hnRNPA1 on mRNA transport and local translation in induced pluripotent stem cell (iPSC)-derived neurons.

## Methods

### iPSC Maintenance and Differentiation

All policies of the NIH Intramural research program were followed for the procurement and use of human iPSCs. iPSCs were previously engineered to express mouse neurogenin 2 (NGN2) under a doxycycline-inducible promoter (14). iPSCs were maintained and differentiated as previously described (15). Briefly, iPSCs were maintained on human embryonic stem cell-qualified Matrigel (Corning 354277) coated plates in Essential 8 (E8, Gibco A1517001) in a 37°C incubator with 5% CO2. Media was replaced every 1-2 days. Cells were passaged with 5-10 minute accutase (Gibco A1110501) treatment at 37°C. Accutase was removed and cells were plated in E8 supplemented with 50 nM Chroman 1 (MedChemExpress HY-15392) to promote survival of single cells. Chroman 1 was removed when cells grew into colonies of 5-10 cells. For neuronal differentiation, cells were digested in Accutase at 37°C for 5-10 minutes and resuspended in neuronal induction media (Knockout DMEM F12 (Gibco 12660012), 1x N2 supplement (Gibco 17502048), 1x non-essential amino acids (ThermoFisher 11140050), 1X Glutamax (ThermoFisher 35050-061), 2 µg/mL Doxycycline (SigmaAldrich D9891), 50 nM Chroman 1). On days 1 and 2, fresh induction media was added to cells. Day 3, cells were split with accutase and plated on a Poly-L-Ornithine (PLO) (Sigma-Aldrich P3655) dissolved in borate buffered saline (SigmaAldrich 08059) coated plate in neuronal maturation media (BrainPhys (STEMCELL Technologies 05790), 1x B27 supplement (Gibco 17504044), 10 ng/mL BDNF (Peprotech 450-02), 1 µg/mL laminin (R&D Systems 3446-005-01), 10 ng/mL NT3 (Peprotech 450-03), and 2 µg/mL Doxycycline). To plate neurons in Boyden chambers, 6-well cell culture inserts with 1 µm pore size transparent PET membranes (Falcon 353102) were coated with PLO overnight. After washing with water and fully drying chambers and holding 6 well plates, the bottoms of the chambers were coated with 10 µg/mL laminin. After plating neurons in the final plate, half media changes were performed every other day. All neuron cultures were differentiated to day 17 post dox induction.

### Sphere Differentiation

To differentiate iPSCs as neurospheres, iPSCs were plated in Corning 384 well round bottom ultra-low attachment spheroid microplates (Corning 4516) after coating with anti-adherence rinsing solution (StemCell Technologies 07010) and washing with PBS twice. iPSCs were split with Accutase as normal and 10,000 cells per well plated in neuronal induction media. The plate was centrifuged at 150 xg for 2 minutes to aggregate cells then incubated at 37°C. On day 1, preheated neuronal induction medium was added. On day 3, recipient plates were coated with PLO and Laminin (15 µg/mL). A wide-orifice pipette was used to gently aspirate the neurosphere and place it in the recipient plate in neuronal maturation media.

### CRISPRi knockdown

iPSCs were previously engineered to express dCas9 tagged to a KRAB transcriptional repressor domain under the CAG promoter (16). sgRNAs targeting TDP-43 or hnRNPA1, as well as a non-targeting guide were packaged into lentivirus for transduction of iPSCs as follows: PLO-coated plates were seeded with Lenti-X HEK cells in high glucose + GlutaMAX DMEM (Life Technologies 10566024) supplemented with 10% FBS (Sigma TMS-013- B). The next day, arrayed transfection mixes were prepared as follows for 2.5 million cells on a 6 well plate: 1.2 µg sgRNA plasmid and 2.4 µg packaging mix (1.6 µg psPAX2, 0.6 µg pMD2G, and 0.2 µg pAdVantage) were combined with 5 µL P3000 reagent (ThermoFisher Scientific L3000015) in 150 µL Opti-MEM (Gibco 31985070); 3.75 µL Lipofectamine 3000 Transfection Reagent (ThermoFisher L3000075) was diluted into 150 µL Opti-MEM and incubated at room temperature for 5 minutes. DNA mixture was added to the diluted Lipofectamine mixture and flicked to mix before incubating at room temperature for 30-45 minutes. Following incubation, mixture was added to cells in 1.5 mL DMEM+FBS dropwise with rocking to mix. The following day, media was changed to 3 mL fresh DMEM+FBS supplemented with 6 µL ViralBoost (Alstem VB100; diluted 1:500 in media). Two days later, media was transferred to a conical tube, centrifuged for 5 minutes at 1100 xg, and supernatant was filtered with a 0.2 µm filter into a new tube to remove cell debris. Lenti-X Concentrator (Takara Bio 631232) was added at 1:3, mixed, and incubated at 4°C for 2- 4 days. After incubation, concentrated virus was pelleted by centrifugation at 1500 xg at 4°C for 45 minutes. Virus pellet was resuspended in cold PBS at 100 µL per mL of media. Virus was aliquoted and stored at −80°C until use. iPSCs were transduced with 100 µL virus per 100,000 cells at 6 well scale or 1 mL per 10 million cells at 15 cm plate scale in E8 Chroman 1. Cells were checked for co-expressed fluorophore the following day and media was changed to fresh E8 Chroman 1. The following day, cells were treated with antibiotic (puromycin or blasticidin, depending on the resistance gene carried in the sgRNA plasmid) for 2 days. Subsequently, differentiation was initiated as usual. sgRNA sequences are as follows: non targeting control: GTCCACCCTTATCTAGGCTA, *TARDBP*: GGGAAGTCAGCCGTGAGACC, *HNRNPA1*: GAGAGAGGGCGAAGGTAGGC. For hnRNPA1-hnRNPA3 compensation immunofluorescence, a dual guide plasmid was used (17). NT guide 1: GTCCCAGGCTCTCCACTATG, NT guide 2: GTACAGCTGGAAGGAAAGCC, *HNRNPA1* guide 1: GAGAGAGGGCGAAGGTAGGC, *HNRNPA1* guide 2: GCGCGCAACAAAGGCTTGGG, *HNRNPA3* guide 1: GTAATACCTCCTCCCCCCGG, HNRNPA3 guide 2: GCGGGCGAGGGGCCCCAAGG.

### Western blot

For sgRNA validation, cells were scraped in PBS, centrifuged at 1500 xg for 5 min, and washed in PBS prior to cell lysis in RIPA buffer (Rockland MB-030-0050). For boyden chamber western blot, the bottom and top of chambers were scraped into PBS and pelleted at 500 xg for 5 min. Cells or cell parts were lysed in RIPA buffer. Lysates were rocked at 4°C for 1 hour and centrifuged at 4°C at top speed for 10 minutes. Clarified lysate was transferred to a new tube and protein was quantified using DCA (BioRad 5000113, 5000114, 5000115). 10 µg protein was mixed with loading dye (BioRad 1610747) with beta-mercaptoethanol (BioRad 1610710) and incubated at 95°C for 5 minutes. Proteins were resolved on a 4–15% Mini-PROTEAN TGX Precast Protein Gel (BioRad 4561086) and then transferred to nitrocellulose membranes (BioRad). Membrane was blocked in a 5% milk solution in TBST for 1 hour, washed in TBST, and incubated with primary antibody overnight at 4°C. The following primary antibodies were used at their respective dilutions: Anti-TDP-43 (Proteintech 10782-2-AP, 1:1000), Anti-HNRNPA1 (Proteintech 11176-1-AP, 1:1000), Anti-RPL24 (Proteintech 17082-1-AP, 1:1000), Anti-Actin (Abcam ab8226, 1:1000), Anti-TUBB3 (Biolegend 801201, 1:1000), Anti-NeuN (Cell Signaling D4G4O, 1:2000). Blots were then washed in TBST and then incubated in a secondary antibody (Rb680 (LiCOR 926-32213) or Ms800 (LiCOR 926-68072) 1:10,000 as appropriate) at room temperature for one hour. Blots were then washed in TBST then PBS before imaging on Odyssey Clx (LiCOR). Densitometry was performed in ImageStudio Lite and graphed in Prism.

### Neurite versus whole cell proteomics

Boyden chambers were washed with cold PBS and neurites were scraped from the underside of the chamber. The top of the chamber was subsequently scraped as the whole cell fraction. Fractionated cells were pelleted, PBS removed, and cells flash frozen before proceeding to SP3/benzonase protocol as described (18). Briefly, samples were resuspended in SP3 buffer (50 mM Tris-HCI [pH = 8.0], 50 mM NaCl, 1% SDS, 1% Triton X-100, 1% NP-40, 1% tween 20, 1% glycerol, 1% sodium deoxycholate (w/v), 5 mM EDTA [pH = 8.0], 5mM dithiothreitol (DTT), 5 KU benzonase, and 1× complete protease inhibitor (Roche 05892970001)), then shaken at 1200 rpm, 65°C for 30 min. Cell lysates were alkylated using 500 mM iodoacetamide (IAA) in the dark and then reduced by 500 mM DTT. Alkylated proteins were captured by hydrophilic magnetic beads using a KingFisher APEX robot, followed by a trypsin/LysC digestion in 50 mM ammonium bicarbonate overnight at 37°C and 1200 rpm. Beads were removed using KingFisher APEX and digested peptides were lyophilized and reconstituted to 0.5 mg/mL with MS loading buffer (2% acetonitrile, 0.1% formic acid). The peptides were analyzed on an Orbitrap Eclipse mass spectrometer, coupled with FAIMS interface using data-independent acquisition (DIA) approach with scan range 400-1,000 m/z, and 24 isolation windows (8 m/z per window). The RAW data files were database searched against UniProtKB Human Database containing 20,435 reviewed entries using Spectronaut 19 with label free or SILAC pipeline. Carbamidomethyl on cysteine was used as fixed modification, and acetylation on protein N-terminus and oxidation on methionine were used as variable modifications. The false discovery rate of protein and peptide level were set at 1%, and cross-run medium normalization was enabled.

### RNA sequencing

Neurons grown in Boyden chambers were fractionated after washing in cold PBS by scraping the underside of chambers to collect neurites and then the top of the chamber to collect the whole cell fraction. RNA was collected using the Qiagen RNeasy Miniprep kit (Qiagen 74104). RNA was quantified with Qubit (ThermoFisher Q32852) and integrity evaluated by BioAnalyzer using the Pico kit (Agilent 5067-1513). We added 10 ng of RNA per sample into the Takara Total RNA-seq Kit v3 - Pico Input Mammalian (634485) and followed manufacturer instructions. Libraries were quantified by Qubit (ThermoFisher Q32850) and size confirmed with BioAnalyzer (Agilent 5067-4626). Libraries were normalized to 3 nM, pooled, and iSeq performed to check normalization. If necessary, re-normalization was performed, and libraries then sequenced (2 x 100 bp, depth about 50 M per sample) with Illumina NovaSeqS1 200.

### UPF1 KD and RNA sequencing

To knock down UPF1, sgRNAs targeting either a non-targeting control guide (GTCCACCCTTATCTAGGCTA) or UPF1 (GGCCAGACGCAGACGCCCCC) were delivered to iPSC by lentiviral transduction as above. iNeurons were differentiated as above. On day 14 post doxycycline induction, iNeurons were collected and RNA was extracted with a RNeasy mini kit (Qiagen 74104) following manufacturer’s instructions and including the on-column DNA digestion step. RNA concentrations were measured using Nanodrop and 500 ng of RNA was used for reverse transcription. Samples undergoing RNA sequencing were further assessed for RNA quality on a TapeStation 4200 (Agilent) and bands were quantified with TapeStation Systems Software v3.2 (Agilent). RNA integrity number (RIN) for iNeurons was above 8.3. Sequencing libraries were prepared with polyA enrichment using a TruSeq Stranded mRNA Prep kit (Illumina 20020594) and sequenced on an Illumina HiSeq 2500 for NovaSeq 6000 machine at UCL Genomics at 2 x 150 bp, depth > 40 M per sample.

### QuanCAT

Neurons were starved of methionine, lysine and arginine amino acids in SILAC medium (Athena Enzyme Systems 0423) for 45 minutes at 37°C. 4 mM AHA was added to light and heavy media along with 0.5 mM Lysine and 0.3 mM Arginine with either the heavy or light isotope of both respectively. Neurons were incubated in the respective media (either light or heavy) for 2 hours at 37°C before methionine free media supplemented with 4 mM methionine was added for 15 min. Following incubation, cells were placed on ice. Bead-based enrichment of AHA-containing proteins and subsequent on bead digestion were performed per manufacturer’s instructions (ThermoFisher C10276) with a few modifications. Briefly, urea lysis buffer supplemented with protease inhibitors (Roche 05892970001) was added to each well lysis and shaken at 65°C at 1200 rpm for 30 minutes. The lysates were then centrifuged for 5 minutes and returned to ice while a 2X solution of 18 megaOhm water, component D (ThermoFisher C10416) and Copper (II) sulfate (ThermoFisher C10416) were prepared. DCA was performed and 10% of lysate was collected as the input fraction and processed by SP3 as above. The remaining 90% of lysate was normalized, added to catalyst and beads solution and rotated end over end at room temperature for 18 hours. The following day, the reaction was centrifuged, and the supernatant was aspirated and collected for SP3 processing (flow through fraction). Water was added to the resin bead mixture and centrifuged again. SDS wash buffer and DTT was added to the beads and was vortexed briefly to resuspend the resin. The resin was heated at 70°C on a heating block for 15 min and then cooled to room temperature before being centrifuged for 5 minutes at 1000 xg, aspirating the waste. Iodoacetamine (IAA) (ThermoFisher A39271) was added to each reaction, the reaction was then vortexed and then incubated in the dark for 30 minutes. A 1 mL pipette was used to resuspend the resin solution and transferred to a gravity column. The resin tube was rinsed with water and the rinse was added to the column, followed by SDS wash buffer and 8 M urea/100 mM Tris pH 8, each repeated 10 times. The final 10 washes were with 20% acetonitrile. The bottom of the column was capped, and a digestion buffer composed of 100 mM Tris, 2 mM CaCl2 and 10% acetonitrile was added to the resin before it was resuspended and transferred to a clean tube. The column was rinsed with a digestion buffer and added to the transferred resin.

The resin was pelleted by centrifugation for 5 minutes at 1000 xg and the supernatant was aspirated to waste leaving some digestion buffer with the resin. Mass spectrometry-grade trypsin (Promega V5071) solubilized to 0.1 µg/µL with 50 mM ammonium bicarbonate was added to the resin slurry, vortexed and incubated for 37°C for 16 hours overnight. The resin was pelleted by centrifugation at 1000 xg and the supernatant was transferred to a clean tube. Water was added to the digest to dilute the acetonitrile to 2%. The digest was then acidified with 10% TFA (ThermoFisher 85183) to a pH lower than 4 and then dried in a vacuum concentrator.

### pSILAC

Neurons were starved of essential amino acids in a similar manner as above and incubated in heavy amino acid media with 0.5 mM lysine and 0.3 mM arginine for 2 hours at 37°C. Neurons were then incubated in methionine free media supplemented with 4 mM for 15 min before they were washed with cold PBS and scraped (top and bottom) into PBS on ice. Cell parts were centrifuged, and supernatant was removed. Pelleted cell parts were resuspended in SP3 buffer supplemented with 1 M DTT, benzonase, and a protease inhibitor tablet (Roche 05892970001) as above and shaken at 1200 rpm for 30 min. The lysate was centrifuged for 5 minutes at 10,000 xg before the supernatant was removed to a new tube and quantified via DCA. Samples were normalized to 40 µg protein. DTT was added to each tube and shaken at room temperature at 1200 rpm for 30 minutes. After adding 500 mM IAA (ThermoFisher A39271) to the reduced lysate, the samples were then shaken in the dark at 25°C for 30 min at 1200 rpm. 500 mM DTT was added to each sample and shook at room temperature at 1200 rpm in the light for 30 minutes. Following this, the preparation proceeded as above with magnetic bead capture and remaining steps.

### Microscopy

#### Boyden chamber axon/dendrite quantification

Cells expressing a cytosolic mScarlet and a GFP-tagged End Binding protein 1 (EB1) were plated in Boyden chambers at approximately 1% of the total cell population and left to differentiate as normal. At day 17, the chamber was removed from the 6 well plate and placed on a coverslip in a temperature and CO2-controlled incubator on a Nikon Ti2 spinning disk confocal using a 60x oil objective with numerical aperture 1.4. Thirty second videos were taken of the underside of the microporous membrane, capturing the RFP channel once (100 ms exposure time, laser power 50%, excitation 561 nm, emission 620 nm) and the GFP channel every 500 ms (100 ms exposure time, laser power 50%, excitation 488 nm, emission 535 nm), spinning disk speed 4000 rpm for each channel. EB1 comets were quantified by agnostically generating kymographs along sections of processes clearly belonging to a single cell (avoiding thicker processes that may have been 2 parallel processes and between branch points that may have been either true branches or separate processes crossing). Kymographs were then manually scored as either containing unidirectional or bidirectional EB1 comets. Eight kymographs were generated in each of seven videos in one chamber and eight videos in a second chamber using the Fiji KymographBuilder. Data was graphed in GraphPad Prism. Z-stacks were also collected to visualize neurite and cell body distribution on either side of the membrane and two maximum intensity projections generated to visualize the underside and top of the membrane. *Immunofluorescence*: Cells were fixed with 4% PFA (paraformaldehyde Electron Microscopy Sciences 15710) and then washed with PBS. Permeabilization was performed with 0.1% Triton-X 100, and permeabilized cells were blocked in 2% bovine serum albumin (BSA, Jackson ImmunoResearch 001000173). Samples were incubated with primary antibodies, rocking overnight at 4°C. The following primary antibodies were used at the respective concentrations and dilutions: Anti-TDP-43 (Proteintech 10782-2-AP, 1:1000), Anti-HNRNPA1 (Proteintech 11176-1-AP, 1:500), Anti-HNRNPA3 (Proteintech 25142-1-AP, 1:1000). Cells were washed with PBS and samples were incubated with goat anti-rabbit secondary antibody (Biotium 20015) at a 1:1000 dilution at room temperature for one hour. Samples were washed and counterstained with DRAQ5 (Thermo 62252) at a 1:1000 dilution. Neurons were then imaged at 20x numerical aperture 0.75 with a Nikon Ti2 spinning disc confocal for KD validation. BFP exposure time 200 ms, binning 2×2, 50% laser power, excitation 405 nm emission 460 nm. GFP exposure time 100 ms, binning 2×2, 25-50% laser power, excitation 488 nm, emission 535 nm. Far red exposure time 100 ms binning 2×2 50% laser power, excitation 640 nm, emission 670 nm. Spinning disk speed was 4000 rpm for all channels. Fluorescent intensity quantification was performed in CellProfiler (19) by identifying neurons containing sgRNA and calculating the mean integrated intensity of the immunofluorescence signal. Data was graphed in GraphPad Prism. Neurons were imaged at 10x, numerical aperture 0.45 with a Nikon Ti2 spinning disc confocal (disk spinning at 4000 rpm) for hnRNPA1-hnRNPA3 staining. BFP exposure time 300 ms, excitation 405 nm laser power 100%, emission 405 nm. GFP exposure time 100 ms, excitation 488 nm laser power 100%, emission 525 nm. Far red exposure time 100 ms, excitation 640 nm laser power 100%, emission 705 nm. Spinning disk speed was 4000 rpm for all channels. Data was analyzed using a custom General Analysis 3 pipeline in Nikon Elements and graphed in GraphPad Prism.

#### HCR FISH

HCR FISH was performed following manufacturer’s recommendations (Molecular Instruments). Briefly, cells were fixed in 4% PFA and permeabilized overnight at −20°C in 70% ethanol. Cells were washed with SSC (2x, Invitrogen AM9770) and pre-hybridized in probe hybridization buffer at 37°C. Probes against ATXN2, UBQLN1, and RPS27A were added to their respective wells and incubated overnight at 37°C. Following washing with probe wash buffer and SSCT, the samples were pre-amplified in amplification buffer at room temperature. HCR Amplifiers (B1-h1-488 and B1-h2-488) were prepared by heating to 95°C and then cooling down to room temperature in the dark. A hairpin solution consisting of snap cooled hairpins and amplification buffer was added to samples to incubate overnight in the dark at room temperature. Samples were washed with SSCT and were stored at 4°C in PBS protected from light before imaging. Immunofluorescence proceeded as normal prior to imaging with anti-Tau (Invitrogen HT7, 1:500) and nuclear counterstaining with Hoechst (1:10,000, Thermo 62249). Cells were imaged at 60x water immersion objective numerical aperture 1.27 on a Nikon Ti2 spinning disc confocal. DAPI exposure time 100 ms, laser power 100%, excitation 405 nm, emission 460 nm; GFP exposure time 100 ms, laser power 50%, excitation 488 nm, emission 535 nm; far red exposure time 100 ms, laser power 100%, excitation 640 nm, emission 670 nm; spinning disk speed 4000 rpm for all channels. FISH puncta were quantified in Nikon Elements using a custom General Analysis 3 pipeline consisting of masking for axons (Tau signal) or cell bodies (Hoechst signal grown by 10 nm radially) prior to counting puncta. Resulting puncta quantities were normalized to nuclei count per image (number of Hoechst-positive objects) and graphed in GraphPad Prism.

#### Neurite outgrowth

iPSCs were transduced with virus to express cytoplasmic mScarlet and nuclear mNeonGreen (16) (I2 and H53), then transduced with sgRNA as normal. iPSCs were differentiated either as spheres or standard differentiation. Spheres contained 100% red-green labeled neurons. On day 3 post dox induction, spheres were replated as above. Fluorescent standard differentiations were replated as a 1% mixture of labeled neurons in 99% unlabeled, wild type neurons. Neurons were imaged at 20x numerical aperture 0.75 on a Nikon Ti2 spinning disc confocal (disk speed 4000 rpm) on days 3, 4, 5, 6, 7, 8 for spheres and days 3, 4, 5, 6, 7, 8, 10, 12, 14, 16, and 18 for 2D differentiation. For sphere differentiation, RFP exposure time 100 ms, 100% laser power, excitation 561 nm, emission 620 nm; GFP exposure time 100 ms, 100% laser power, excitation 488 nm, emission 535 nm; BFP exposure time 100 ms, 100% laser power, excitation 405 nm, emission 460 nm. Spinning disk speed 4000 rpm for all channels. Overlapping images were stitched to obtain an image of the entire well. For 2D differentiation, RFP exposure time 100 ms, 100% laser power, excitation 561 nm, emission 620 nm; GFP exposure time 100 ms, 100% laser power, excitation 488 nm, emission 535 nm; BFP exposure time 100 ms, 100% laser power, excitation 405 nm, emission 460 nm. Spinning disk speed 4000 rpm for all channels. Overlapping images were stitched to obtain an image of the entire well. To obtain sphere outgrowth radius, a custom General Analysis 3 pipeline in Nikon Elements was used to obtain the EqDiameter of the neurites (red channel) and the cell bodies (green channel). For standard differentiation, a custom General Analysis 3 pipeline in Nikon Elements was used to threshold, skeletonize, and calculate the line length, count branch points and endings of red neurites, and count the number of cells (green nuclei). Resulting data was graphed in Prism.

#### PolyA FISH

polyA FISH was performed as established (20). Briefly, polyA probes (TTTTTTTTTTTTTTTTTTTTTTTTACACTCGGACCTCGTCGACATGCATT, IDT) were hybridized with the fluorescently labeled FLAP (/5Cy3/AATGCATGTCGACGAGGTCCGAGTGTAA/3Cy3Sp/, IDT) by heating to 85°C then incrementally decreasing temperature to 25°C. Neurons were fixed with 4% paraformaldehyde (Electron Microscopy Sciences 15710) for 10 minutes, then washed three times with PBS. 70% ethanol was added for one to two hours at 4°C. Ethanol was removed and cells were washed twice with PBS and cells were incubated in a pre-hybridization buffer (2x SSC (Invitrogen AM9770), 10% deionized formamide (Millipore S4117)) for 30 minutes at room temperature. Hybridized probes were added in 1X SSC, 0.34 µg/µL *E. coli* tRNA (Sigma R1753), 15% deionized formamide, 2 mM VRC (Sigma R3380), 10% dextran sulfate (MP Biomedicals 160111) and anti- NEFL antibody (Cell Signaling 2837, 1:1000). Cells were incubated at 37°C overnight. Samples were washed twice with 2x SCC and then anti-rabbit secondary antibody (Biotium 20015) at a 1:1000 dilution was added to the cells for 30 minutes at 37°C. Cells were washed twice with SCC and stained with DRAQ5 in 2X SSC for 5 minutes, followed by a wash in PBS. Neurons were imaged with a 60x water immersion objective, 1.27 numerical aperture with a Nikon Ti2 spinning disc confocal, spinning disk at 4000 rpm for all colors. BFP was imaged with 300 ms exposure, excitation at 405 nm with laser power 70%, emission 455 nm. GFP was imaged with 100 ms exposure, excitation at 488 nm with laser power 100%, emission 525 nm. Far red was imaged with 200-300 ms exposure, excitation at 640 nm with laser power 100%, emission 705 nm. Images were analyzed using a custom General Analysis 3 Pipeline in Nikon Elements software to determine FISH intensity in neurites at least 10 nm away from a cell body and then per well data were median normalized and outliers removed in GraphPad Prism for three independent differentiations (replicates). Averages for each condition for each replicate were used for a one-way ANOVA, with Šídák’s multiple comparisons test used to calculate p values for TDP-43 or hnRNPA1 KD compared to the NT condition.

#### FUNCAT

FUNCAT was performed as previously established (21). Cells were starved of methionine (SILAC DMEM/F12 (AthenaES 0423) supplemented with B27, BDNF, laminin, NT3 (concentrations and catalog numbers as above), arginine (0.5 mM, Sigma 11039), leucine (0.8 mM, Sigma 61819), and lysine (1 mM Sigma 62929)) for 45 minutes. Cells were then incubated in either 40 mM azidohomoalanine (AHA) (SigmaAldrich, 900892) supplemented methionine-free medium, 40 mM AHA supplemented methionine-free medium plus 1 mM cycloheximide (Sigma C4859) or anisomycin (Sigma Aldrich, A5862), or 40 mM methionine (Met) (Sigma 64319) supplemented methionine-free medium for two hours. After incubation, cells treated with 40 mM Met supplemented methionine-free medium for 15 minutes. Plates were put on ice and washed twice with cold PBS-MC (1 mM MgCl2, 0.1 mM CaCl2, in PBS), followed by a twenty-minute fixation period in PBS-MC with 4% paraformaldehyde (PFA, Electron Microscopy 15710) and 4% sucrose (Sigma 84100). Following fixation, cells were washed three times with PBS pH 7.4, then incubated in B-block (10% horse serum (Thermo Fisher Scientific, 16050130), 5% sucrose (Sigma Aldrich, 84100-1KG), 2% bovine serum albumin (BSA; Jackson Laboratories, 152734), 0.1% TritonX-100 (Sigma Aldrich, X100-100ML). Following three PBS pH 7.8 washes, FUNCAT reaction mixture was added to the cells, containing 200 µM Tris [(1-benzyl-1H-1,2,3-triazol-4- yl)methyl]amine (TBTA; Sigma Aldrich, 678937-50MG), 500 µM Tris(2-carboxyethyl)phosphine hydrochloride (TCEP; Sigma Aldrich, 75259-1G), 0.4 µM AlexaFluor 488 Alkyne (Thermo Fisher Scientific, A10267) and 200 µM copper (II) sulfate (Sigma Aldrich, 451657-10G). Cells were incubated overnight with rocking at room temperature. The following day, cells were washed in wash buffer (0.5 mM EDTA and 1% Tween-20 in PBS 1x pH 7.8) three times. 1000x DRAQ5 (Thermo 62252) was added for 5 minutes, and cells were washed three times in PBS pH 7.4. Following the final wash, plates were then images on the Nikon Ti2 spinning disc confocal with a 60x water immersion objective, 1.27 numerical aperture. BFP was imaged with 300 ms exposure, excitation at 405 nm with laser power 70%, emission 455 nm. GFP was imaged with 100 ms exposure, excitation at 488 nm with laser power 100%, emission 525 nm. RFP was imaged with 200 ms exposure, excitation at 561 nm with laser power 100%, emission 605 nm. Far red was imaged with 100 ms exposure, excitation at 640 nm with laser power 100%, emission 705 nm. Spinning disk was at 4000 rpm for all colors. Images were analyzed using a custom General Analysis 3 Pipeline in Nikon Elements software to determine FUNCAT intensity in neurites at least 10 nm away from a cell body and then per well data were median normalized and outliers removed in GraphPad Prism for three independent differentiations (replicates). Averages for each condition for each replicate were used for a one-way ANOVA, with Šídák’s multiple comparisons test used to calculate p values for TDP-43 or hnRNPA1 KD compared to the NT condition.

### Data analysis

#### Bulk RNA seq analysis

Demultiplexed 100-bp paired-end reads were aligned to the GRCh38.p12 human reference using STAR two-pass version 2.7.10b (22). Reads were trimmed using Cutadapt version 4.4Bams were indexed with samtools version 1.16.1 (23). Reads were then counted in genes with the featureCounts program of the subread package v2.0.3 (24), using the GENCODE release 28 annotations. PCA was performed using counts normalized with the DESeq2 v1.38.3 variance stabilizing transform (25). Differential expression analysis was conducted using raw counts provided to DESeq2. Most comparisons presented here pertain to the targeting factor, i.e., whether the sample used targeting or non-targeting guides. All contrasts examining transcriptional differences between neurites and the rest of the cell were performed using ’pair’ as a blocking factor in a model of ’∼pair + zone’, where ’pair’ indicates each neurite/whole-cell sample pair. The MA plots display the shrunken log2 fold change calculated by DESeq2 using the ’ashr’ method. Wald tests of significance used a log2(fold change) cutoff of 0.5. Gene ontology analysis used an overrepresentation analysis using clusterProfiler v4.6.2 (26,27) and terms from KEGG and MSigDB’s biological process (BP), cellular component (CC), and molecular function (MF) ontologies (28–31).

#### Splicing analysis

Aligned BAMs were used as input to MAJIQ (v2.4) (32) for differential splicing analysis of whole cell samples using the GRCh38 reference genome. A threshold of 10% Δ*PSI* was used for calling the probability of significant change between KD and NT samples. The results of the MAJIQ Modulizer were parsed to obtain *PSI* and probability of change for each junction. Cryptic splicing was defined as junctions with *PSI* < 5% in control samples and Δ*PSI* > 10%. IsoformSwitchAnalyzeR (33) was used to analyze alternative splicing and isoform switches. Reads were pseudo aligned using Salmon (v1.4) (34), and counts were used as input to IsoformSwitchAnalyzeR. QAPA (35) was used to infer and quantify alternative polyadenylation sites. gProfiler (36) was used for GO analysis of NMD-sensitive transcripts; figure displays only non-redundant terms. Transite (37) was used to identify the overrepresentation of potential RBP targets in 3’ UTR of differentially expressed transcripts. The analysis was performed on transcripts significantly enriched (|logFoldChange| > 5) in neurites or whole cell fraction. The information about binding dynamics and preferred subcellular localization of the most overrepresented RBP in each fraction (RBM8A and SRSF7, respectively) was obtained at https://chronology.rna.snu.ac.kr/ (38).

#### Proteomics analysis

Mass spectrometry data for soma and neurite fractions were processed with ProtPipe (39) to first convert .raw files to mzML format (https://github.com/ProteoWizard/pwiz) and conduct DIA using DIA-NN (40). For the WT experiment, no samples were excluded (Figure S1C-E). Three samples (TDP43-Soma-NT_6, TDP43-Soma-KD_6, HNRNPA1-Neurite-NT_2) were excluded due to their low numbers of protein groups (Supplementary Figure S4I), intensity values that did not respond to median-normalization (Supplementary Figure S4K), or unexpected pairwise sample correlation patterns (Supplementary Figure S4J). The remaining samples were used for principal component analysis (PCA), differential abundance analysis, and enrichment analysis. Differential abundance analysis was conducted by t-tests contrasting pairs of interest, correcting p-values with the Benjamini-Hochberg method. KEGG and GO analysis was conducted using the R package ClusterProfiler v4.10.1 (27) using sets of enriched (or depleted) genes with an FDR-adjusted p-value ≤ 0.05.

#### pSILAC and QuanCAT analysis

Protein quantification in label-free samples was conducted using Spectronaut v18 with default settings. For database search, the library-free module (direct-DIA) was utilized, using the Swiss-Prot human proteome comprising 20,435 reviewed proteins as the reference database. Enzyme specificity (trypsin) with up to two missed cleavages was applied. Fixed modification included carbamidomethylation of cysteine, while variable modifications encompassed acetylation of protein N-termini and oxidized methionine. Peptide and protein false discovery rates were set to 1%.

For pSILAC DIA-MS datasets, database searching employed the default settings for Pulsar search with specific modifications: the "Labeling Applied" option was enabled, and SILAC labels ("Arg10" and "Lys8") were specified in the second channel.

Proteins with low intensity (Supplementary Figure S3A,B,F,G) were excluded from further analysis. The remaining samples and proteins were subjected to differential abundance and enrichment analyses. Statistical analysis was performed using R software (version 4.0.3) with additional packages. Differential expression tests were carried out using the Limma (version 3.58.1), while KEGG and GO enrichment analyses were conducted using ClusterProfiler v4.10.1. Plots were generated using the R package “ggplot2.”

## Results

### Boyden chambers neurite fractionation from iPSC-derived neurons

We first set out to determine if Boyden chambers adequately isolate neurites of iNeurons. Boyden chambers have previously been used to obtain pure neuritic fractions from other neuron types (41), but to our knowledge, they have not previously been used on iPSC-derived neurons. We differentiated iPSCs into iNeurons (14) and plated pre-differentiated neurons on the top of the microporous membrane (Figure 1A). Visualization of the neurons in the chamber reveals separation of the whole cell (top) from a pure neuritic fraction (bottom) (Figure 1B), as was previously described (41). As not every neuronal process is an axon, microtubule growth directionality dynamics were assessed to determine the proportion of axons on the underside of the chamber. GFP-tagged end binding protein 1 (EB1) shows primarily unidirectional growth, demonstrating that most processes on the bottom of the membrane are axons, which have unipolar microtubules (42) (Supplementary Figure S1A-B). However, since ∼20% of the processes on the underside of the Boyden membrane are not axons, we will refer to this fraction as the “neuritic” fraction going forward. Western blot further validates separation of cell bodies from neurites, as the nuclear protein NeuN was not detected in the neuritic fraction (Figure 1C). To identify the proteins enriched in each fraction, we performed proteomics on the neurite and whole cell fractions. As expected, nuclear proteins were not identified in the neurite fraction (Figure 1D). Principal component analysis shows clear separation of neurite and whole cell samples (Supplementary Figure S1F). Pathway analysis on proteins enriched in the neuritic fraction shows terms related to axon health and function, including axon guidance, actin regulation, and various signaling pathways (Supplementary Figure S1G). Conversely, terms enriched in the whole cell fraction shows pathways related to nuclear functions, like the spliceosome, DNA repair, and nucleocytoplasmic transport (Supplementary Figure S1G). Taken together, Boyden chambers efficiently isolate neurites of iPSC-derived neurons and enable collection of sufficient neuritic material for omics.

**Figure 1:**
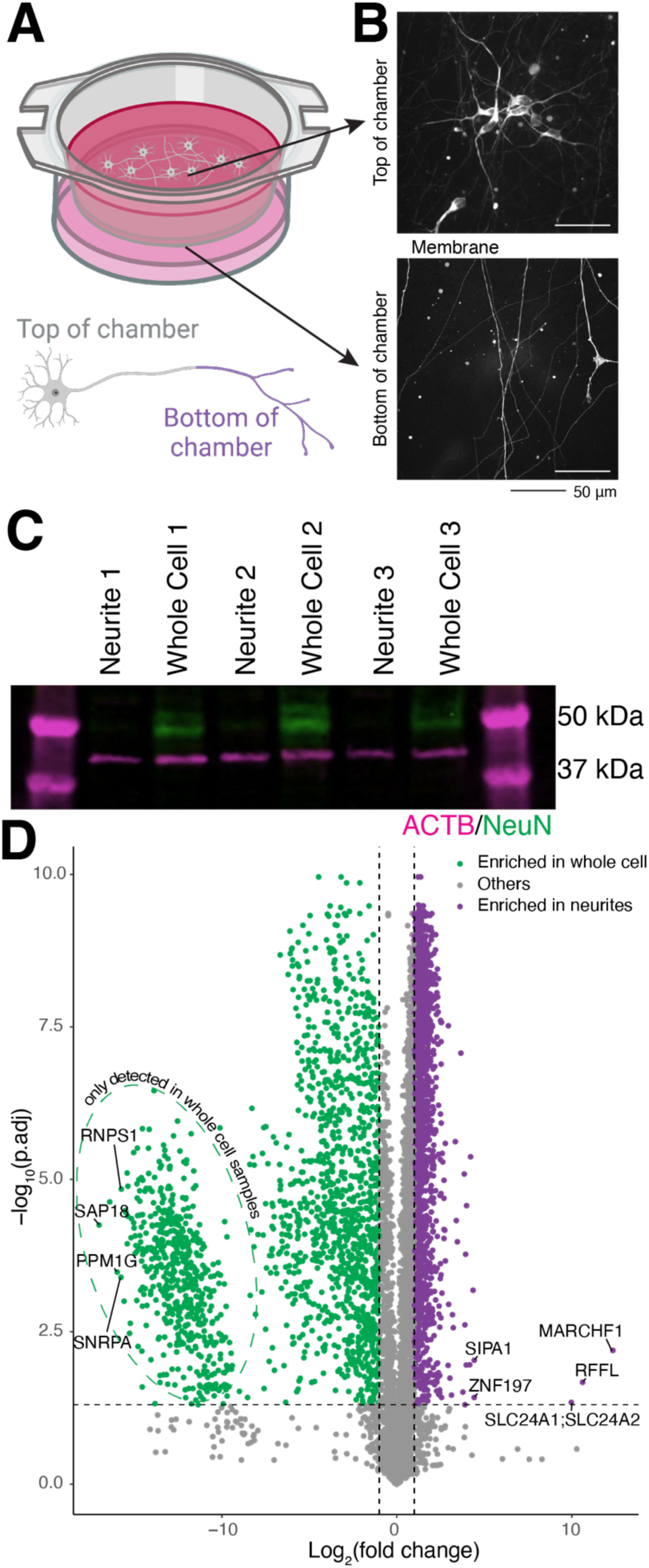
Boyden chambers enable collection of a pure neuritic fraction. A. Schematic of Boyden chambers. Chambers enable separation of neurites (bottom) from whole cell (top) via a thin microporous membrane. B. Images of neurons expressing cytosolic mScarlet in approximately 1% of cells cultured in Boyden chamber. Images were captured as a z-stack and maximum intensity projected. Scale bar 50 µm. C. Western blot validation of neuron fractionation using anti-ACTB (magenta) as a whole cell marker and anti-NeuN (green) as a nuclear marker. n=3, fractionated into 3 neurite samples and 3 whole cell samples. Molecular weight markers are indicated on the right. D. Volcano plot of WT proteomics reveal more whole cell enriched proteins (log2(fold change) < 0, left) than those enriched in neurites. Dashed horizontal line represents adjusted p- value = 0.05. Vertical dashed lines identify proteins with log2(fold change) of 1 or −1. n=6 neurite samples, 5 whole cell samples.

### Thousands of transcripts are identified in neurites

To identify the mRNAs transported into neurites in iPSC-derived neurons, we collected neuritic and whole cell fractions of wildtype (WT) neurons 17 days after doxycycline NGN2 induction and performed transcriptomics. We discovered thousands of RNAs enriched in neurites compared to the whole cell fraction, including some unexpected transcripts, like ribosomal proteins and RBPs (Figure 2A). Principal component analysis (PCA) reveals that 97% of the variance in the samples comes from its cellular fraction, vastly outweighing the sample-to-sample variance (1%) (Supplementary Figure S2A). Quality control metrics show similar expression efficiencies and number of genes identified between neurite and whole cell fractions (Supplementary Figure S2B- C). Percent alignment to rRNA was higher in neurites, but snoRNAs were depleted from neurite fractions (Supplementary Figure S2B,D). KEGG pathway analysis of the transcripts enriched in the neuritic fraction reveals ribosomal protein transcripts among the most overrepresented terms (Figure 2B), while GO Biological process shows terms related to transport, metabolism, and cytoskeleton (Supplementary Figure S2E). Interestingly, terms associated with neurodegenerative diseases, like ALS, Parkinson’s disease, Huntington’s disease, and generic pathways of neurodegeneration are also highly enriched when examining transcripts identified in neurites (Figure 2B). We used FISH to validate the neuritic localization of a few transcripts, including ATXN2 (an ALS-associated gene (43)), the small ribosomal subunit protein RPS27A, and UBQLN1 (a protein degradation chaperone involved in multiple neurodegenerative diseases (44,45)). Indeed, for these mRNAs, we observe more FISH puncta in neurites than in nuclei (Figure 2C-D), thus validating the neuritic enrichment of these transcripts.

**Figure 2:**
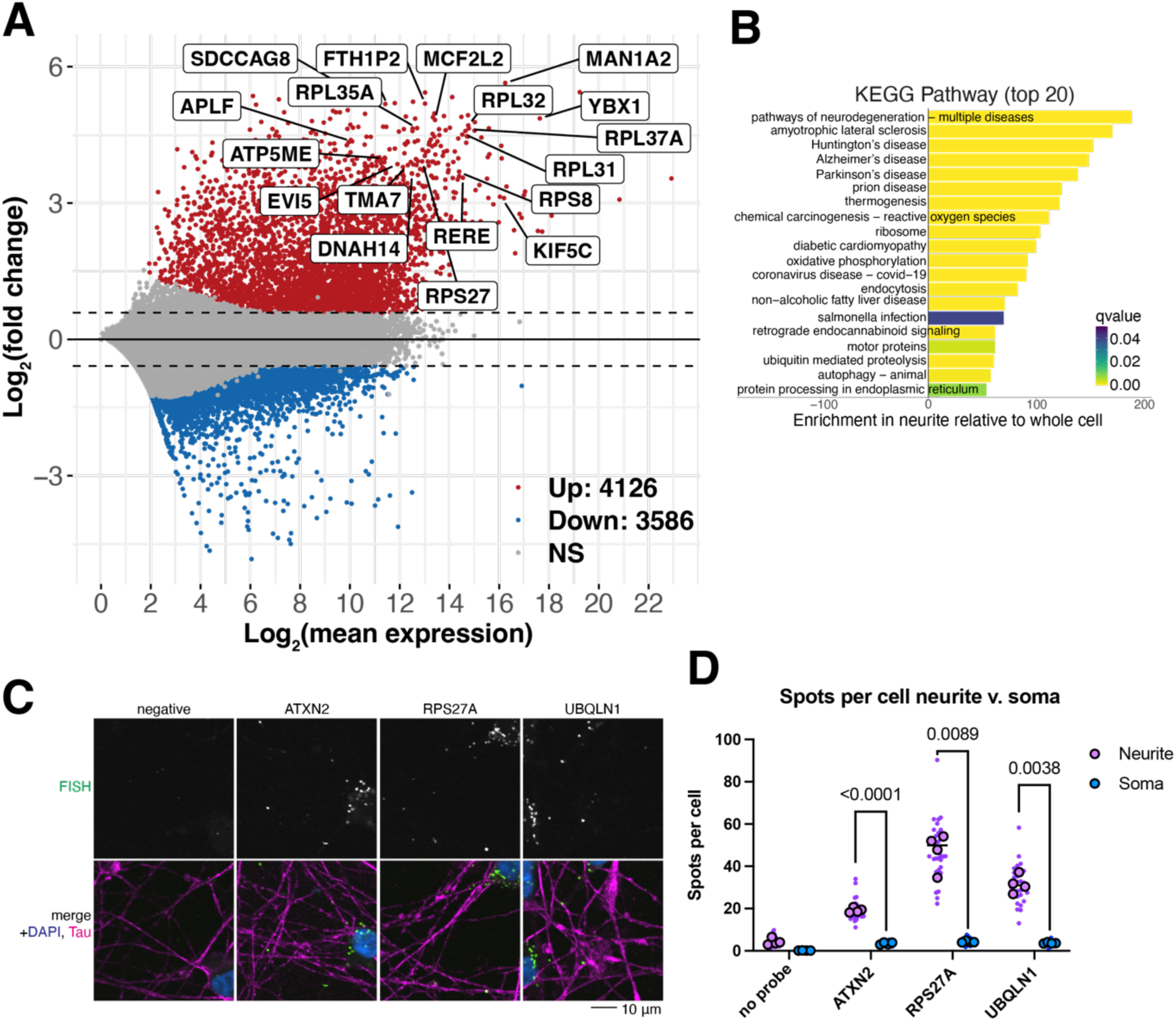
Thousands of transcripts are enriched in neurites. A. MA plot of transcripts identified in WT neuron fractions. Points above the horizontal dashed line are genes that are enriched in neurites while those below are enriched in the whole cell fraction. Y-axis indicates log2(fold change) of a transcript’s relative abundance between neurites and whole cell fractions. An FDR threshold of 0.05 and a fold change threshold of log2(1.5) were applied. Red, blue, and gray represent an increase, decrease, or no change respectively of a gene’s relative abundance. Top 20 genes with largest fold change are indicated by name. n=4 samples each neurite and whole cell fractions. B. KEGG pathway analysis of transcripts localized to neurites reveals enrichment in terms related to neurodegeneration, including ALS, Parkinson’s disease, and Huntington’s disease. C. Representative images of HCR FISH of three highly neuritically enriched transcripts, ATXN2, RPS27A, and UBQLN1. These genes were picked as they were highly differentially localized and are highly expressed in iNeurons. FISH signal in green, anti-Tau staining in magenta, and nuclear stain (Hoechst) in blue. Scale bar 20 µm. D. Quantification of FISH in C, illustrating increased transcripts found in neurites compared to soma, as expected from transcriptomics in A. Small purple/blue points represent the value per image, n=8 per well. Large dots represent the average of each image per well, n=4. Spots identified in cell bodies (grown from Hoechst signal) were divided by the number of nuclei per image. Spots identified in neurites (Tau staining) were divided by the number of nuclei in that image to pseudo-normalize to cell number. P-values from two-way ANOVA with corrections by Šídák’s multiple comparisons test.

### Neurites have shorter isoforms that more often use proximal 3’ UTRs

To determine whether transcripts transported to neurites have any common features that might contribute to their transport, we used IsoformSwitchAnalyzeR, a tool to evaluate transcriptome-wide changes in certain types of alternative splicing (33), to evaluate the changes to isoforms in our WT neurons in neurites. When comparing the neurite to whole cell fractions, we found that genes expressed in both compartments more often have nonsense mediated decay (NMD)- sensitive transcripts in the neuritic fraction than the whole cell fraction (Figure 3A). To investigate this further, we performed a Gene Ontology (GO) analysis on the genes upregulated in neurites with NMD-sensitive transcripts. The NMD-sensitive transcripts are enriched for cytoplasmic, transport, and metabolism terms (Figure 3B). Further, we analyzed genes that switch from transcripts that predicted as NMD-sensitive in the neuritic fraction to NMD-insensitive in the whole cell fraction. We examined the expression of these isoforms in the UPF1 knockdown dataset (manuscript in preparation) and confirmed that the isoforms upregulated in the neuritic fraction (NMD-sensitive) also significantly increase upon UPF1 knockdown, while the isoforms of these genes which are upregulated in the whole cell fraction (NMD-insensitive) do not change (Figure 3C). We also find that on average, neuritic transcripts are shorter or have shorter 3’ untranslated regions (UTRs) (Figure 3A). In contrast, others have found increased 3’UTR length in neuronal processes (46) or that certain categories of transcripts are increased in axons, particularly categories like axon growth and metabolism-associated transcripts (47). Of note, transcripts with shorter UTRs are translated more often (48). The bias we observe towards shorter transcripts may be related to the young developmental stage of our iPSC-derived neurons. We also observe an increased use of the most proximal UTR, the UTR closest to the end of the coding sequence, in the neurites compared to the whole cell (Figure 3D-E). This preference suggests a potential mechanism by which the cell sorts transcripts for transport to distal sites.

**Figure 3:**
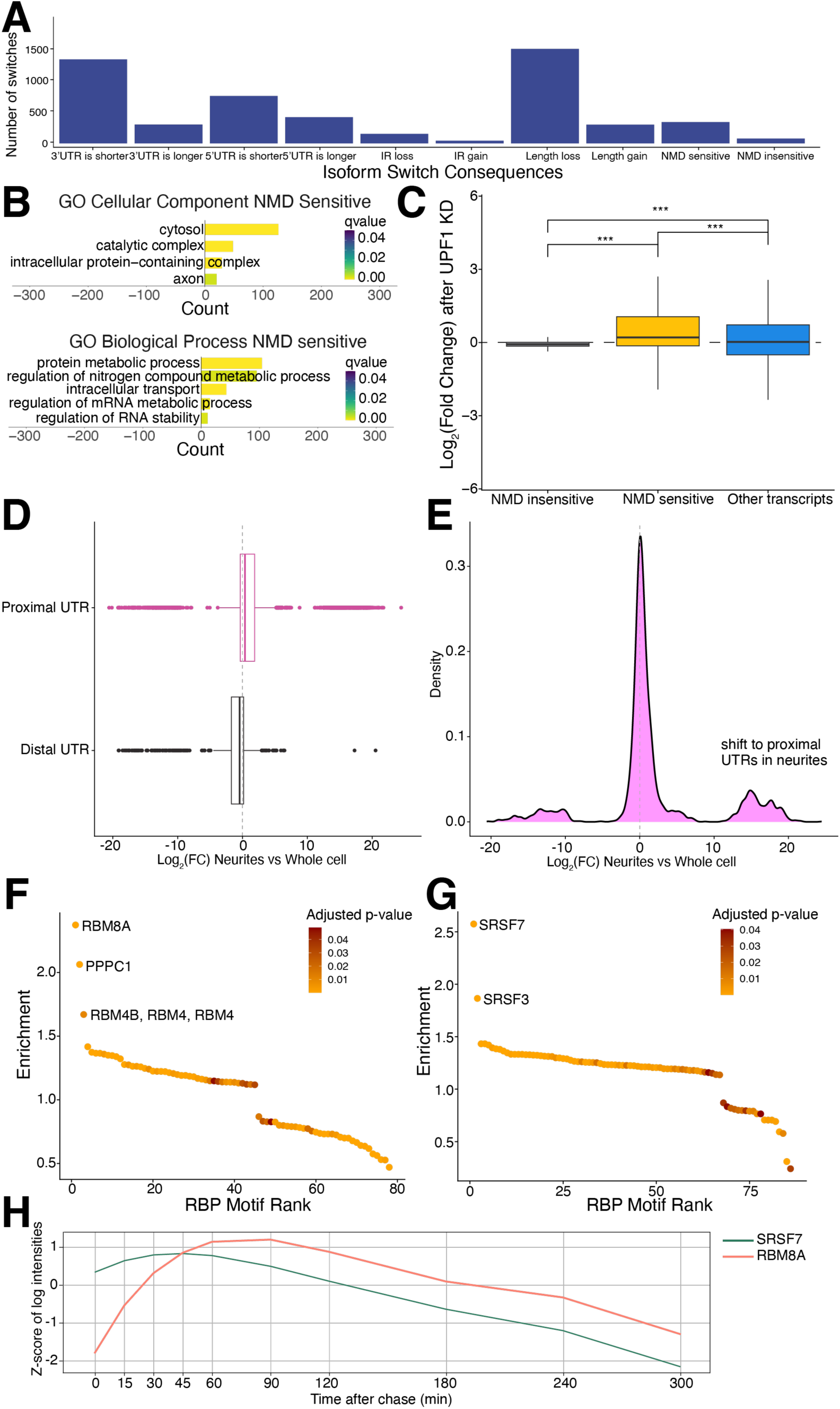
Neuritic transcripts are shorter and more likely to use the most proximal UTR. A. Isoform switch consequences in WT neurites compared to whole cell fraction. Neurites are more likely to have shorter isoforms. IR=intron retention, NMD=nonsense mediated decay. B. GO Cellular Component of NMD sensitive genes from IsoformSwitchAnalzyzeR shows cytoplasmic related terms, while GO Biological Process shows protein and RNA metabolism related terms. C. Expression of genes that switch from NMD-sensitive in neurites to NMD-insensitive in whole cell fractions in UPF1 KD neurons shows that the neuritic NMD-sensitive isoforms have increased expression after UPF1 KD while whole cell NMD-insensitive isoforms do not change, demonstrating they are indeed sensitive to NMD. D. Plot of 3’ UTR usage. Neurites are more likely to use the most proximal UTR (pink) than a more distal UTR (black). E. Density plot of UTR usage. Neurites use more proximal UTRs rather than more distal UTRs. F. Transite analysis of transcripts enriched in neuritic fraction (LFC > 5) identifies RBM8A as the highest scoring RBP enriched in the 3’ UTR of those transcripts. Motif rank plot shows distribution of RBP binding motif enrichment score (y-axis) ordered by RBP motif enrichment rank (x-axis). Color indicates significance of enrichment. Only a subset of significantly enriched RBP binding motifs after multiple testing correction is shown (adjusted p-value ≤ 0.05). G. Transite analysis of transcripts enriched in the whole cell fraction (LFC < −5) identifies SRSF7 as the highest scoring RBP enriched in the 3’ UTR of those transcripts. Motif rank plot shows distribution of RBP binding motif enrichment score (y-axis) ordered by RBP motif enrichment rank (x-axis). Color indicates significance of enrichment. Only a subset of significantly enriched RBP binding motifs after multiple testing correction is shown (adjusted p-value ≤ 0.05). H. RNA binding dynamics of SRSF7 and RBM8A from https://chronology.rna.snu.ac.kr show that SRSF7 binds to mRNAs early in the lifespan of the transcript (peak at 42.8 minutes) while RBM8A binds later (peak at 77.3 minutes).

Additionally, we analyzed transcripts enriched in the neuritic and whole cell fractions to identify potential RBPs regulating subcellular localization or post-transcriptional processes using Transite (37). This tool infers RBPs driving global gene expression changes based on the RBP binding motifs enriched in the 3’ UTRs of differentially expressed genes (37). RBM8A was the highest scoring RBP that emerged from the analysis of neurite-specific transcripts (Figure 3F). RBM8A regulates mRNA splicing, stability, and translation, and was shown to be localized in both the soma and dendrites of neurons (49). SRSF7 had the highest enrichment score in the analysis of whole cell-specific transcripts (Figure 3G); it is an essential RBP required for pre-mRNA splicing and is localized predominantly to the nucleus. To understand the RNA-binding dynamics of the distinct RBPs driving differential expression of our neurite versus whole cell transcripts, we analyzed RBM8A and SRSF7 using an interactive web application (https://chronology.rna.snu.ac.kr) (38) (Figure 3H). According to this classification, SRSF7 is an early binder to mRNA (a peak binding time of 42.8 minutes) from the temporal cluster II associated with pre-mRNA splicing (38). RBM8A has a later peak binding time to mRNA of 77.3 minutes and is included in the temporal cluster V lined with nonsense-mediated decay, mRNA transport, and translation (38). These observations support RBM8A as an important regulator of mRNA processing in neurites.

### Newly synthesized proteins enriched for neuritically localized transcripts

To determine if the neuritically localized transcripts are translated locally, we used two complementary mass spectrometry approaches using the incorporation of heavy amino acids. The first, QuanCAT (50), uses a methionine analog with an azide group, L-Azidohomoalanine (AHA), to enrich for newly synthesized proteins via click chemistry (51) onto an alkyne agarose bead (Figure 4A). Neurons are pulsed with AHA and heavy arginine and lysine for 2 hours prior to whole cell lysis (Figure 4A). We used this short pulse to reduce the effect of protein degradation on heavy amino acid incorporation, ensuring all proteins that incorporate heavy amino acids were newly synthesized. As a control, we exposed cells to AHA without the presence of heavy amino acids (light samples). Interestingly, even though we performed this experiment on whole cells due to the amount of material needed for bead enrichment, many of the proteins we identified as newly synthesized are translated from transcripts transported to neurites (Figure 2A, 4B). We also collected an input fraction prior to on-bead enrichment and a flow through fraction to determine how many heavy proteins are missed by the beads (Figure 4A). Comparing the intensity of the heavy and light mass spec channels, there were few proteins found in the flow through and the QuanCAT samples were enriched for heavy labeled peptides (Supplementary Figure S3A). However, the number of protein groups identified is drastically reduced in the QuanCAT samples compared to the input or flow through, demonstrating enrichment of the small number of proteins that are labeled with heavy amino acids (Supplementary Figure S3B). We identified 267 heavy labeled proteins in the QuanCAT samples (Figure 4B). In comparison, we only identified 26 heavy labeled proteins in the input fraction and 34 in the flow through fraction (Supplementary Figure S3C-E). As expected, many of the QuanCAT enriched proteins are cytoskeletal proteins like ACTB and various tubulins. We also identify several ribosomal proteins and RNA binding proteins like hnRNPA1, hnRNPK, and IGF2BP1. Indeed, pathway analysis of these proteins shows terms related to ALS and Parkinson’s disease (Figure 4C), as was found for the neurite enriched transcripts (Figure 2B), and the cellular component terms are highly related to cytoskeleton and axons/cell projections (Figure 4D). As such, considering our short pulse time, it seems that the most frequently synthesized proteins in neurons are the expected ones, like cytoskeletal proteins, with some surprises including RNA binding proteins like FUS, MATR3, NONO, TAF15, IGF2BP1, and several hnRNPs, including hnRNPA1, hnRNPA2B1, and hnRNPK.

**Figure 4:**
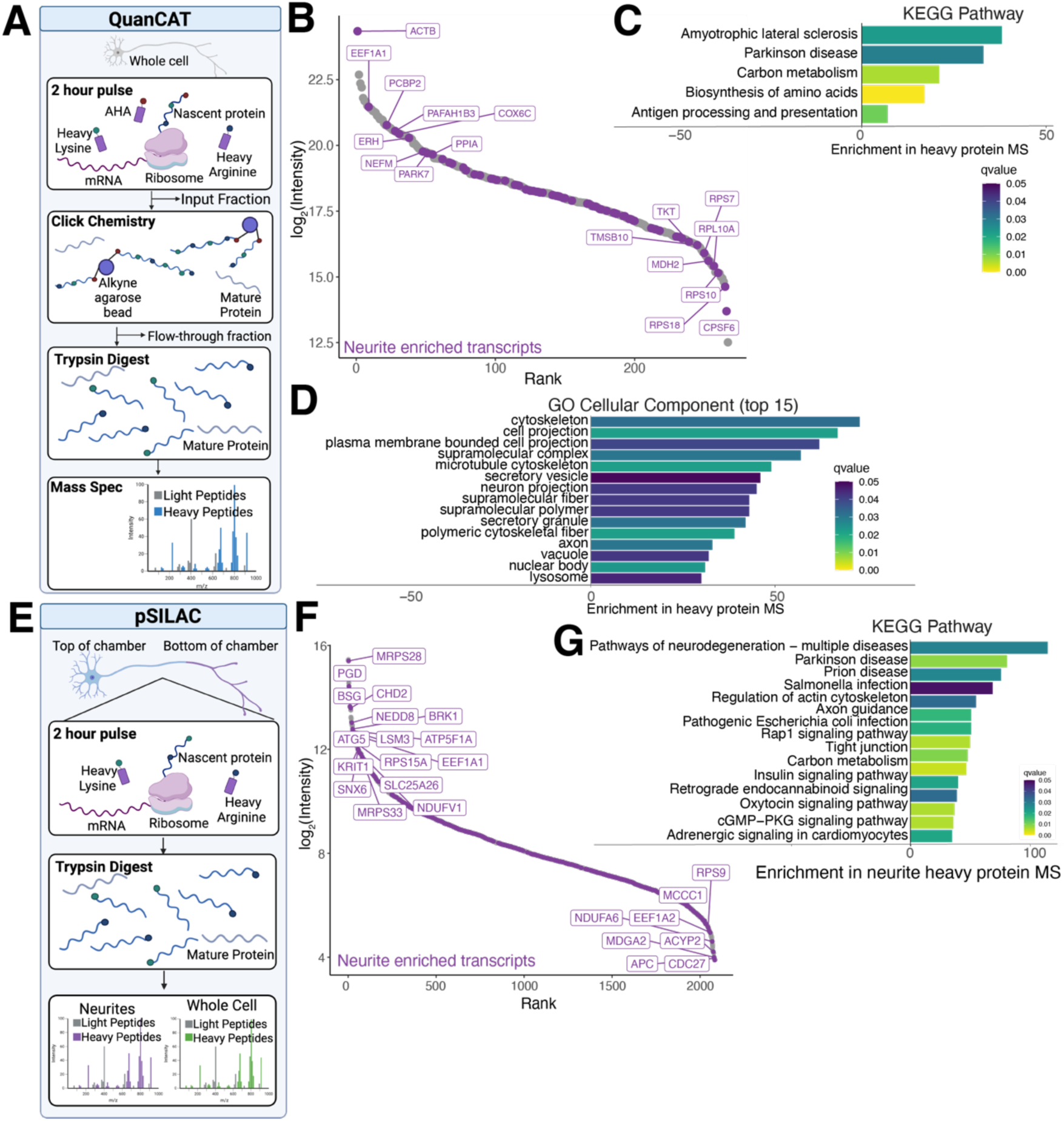
Identification of newly synthesized proteins identifies many neuritically localized mRNAs. A. Schematic of QuanCAT experiment. Neurons are treated with a 2-hour pulse of the methionine analog AHA as well as heavy arginine and lysine. 10% of the resulting lysate was kept as an input fraction, while the remaining lysate was bound to alkyne agarose beads using click chemistry. Following binding, the unbound proteins were retained as a flow through fraction. Beads were stringently washed and proteins were trypsin digested on beads prior to mass spec. B. Rank plot of the proteins containing heavy amino acids identified in neurons. Neurite enriched transcripts (Figure 2A) are indicated in purple. ACTB is the protein with the highest intensity, while several mitochondrial and ribosomal proteins are also identified as newly synthesized. n=3. C. KEGG pathway analysis of proteins containing heavy amino acids identify ALS and Parkinson’s disease as the highest enriched pathways. D. Top Cellular Component terms enriched in proteins that incorporate heavy amino acids include cytoskeletal and axon-related terms. E. Schematic of pSILAC experiment. Neurons were grown in Boyden chambers and exposed to a two hour pulse of heavy arginine and lysine prior to chamber scraping and cell lysis. Proteins were trypsin digested and mass spec performed on both fractions. F. Rank plot of proteins containing heavy amino acids identified in neurites. Neurite enriched transcripts (Figure 2A) are indicated in purple. MRPS28, a mitochondrial ribosomal protein, is the most highly abundantly translated protein detected. n=4 neurite samples. G. KEGG pathway analysis of proteins containing heavy amino acids in neurites identifies pathways found in the transcriptomics GO analysis (Figure 2B), including pathways of neurodegeneration, as well as terms related to axon growth, like axon guidance.

Given that almost a quarter of the newly synthesized proteins in whole cells are from transcripts enriched in neurites, we performed a pulsed stable isotope labeling by amino acids in cell culture (pSILAC) experiment on neurons grown in Boyden chambers to identify newly synthesized proteins in neurites. We used the same two-hour pulse of heavy arginine and lysine prior to neuron fractionation and trypsin digestion of proteins (Figure 4E). As expected, due to the lack of enrichment and short pulse, fewer heavy labeled proteins were detected than light proteins (Supplementary Figure S3F-G). However, in comparing the heavy labeled proteins identified in the neurite and whole cell fractions, we were able to detect heavy labeled proteins significantly enriched in both fractions (Supplementary Figure 3H). We find only 23 proteins enriched in the neurite fraction compared to the whole cell fraction; however, we can detect over 2000 heavy labeled proteins in the neurite fraction (Figure 4F). This seeming disparity is expected as the whole cell fraction contains neurites as well as cell bodies, and while some of the highly translated proteins detected by QuanCAT, like ACTB, are detected at high levels in neurites, their intensity is higher in the whole cell fraction. However, like QuanCAT, many of the proteins encoded by neurite enriched transcripts incorporate heavy amino acids in the neurites (Figure 2A, 4F), suggesting that to some degree, these transcripts are locally translated. Pathway analysis of neurite versus whole cell enriched proteins shows many nuclear and cell body related pathways and functions, reflective of the increased abundance of newly synthesized proteins in those compartments (Supplementary Figure S3I-J). However, pathway analysis of all proteins that were detected as newly synthesized in neurites shows pathways related to neuronal functions, including signaling pathways and axons, as well as pathways related to neurodegeneration (Figure 4G). As such, many of the proteins encoded by transcripts enriched in neurites are translated locally.

### RBP KDs cause few protein mislocalizations but hnRNPA3 compensates for hnRNPA1 KD

To understand the effect of loss-of-function of two FTD/ALS-associated RBPs on the neuritic proteome, we used CRISPRi (16) to knock down TDP-43 and hnRNPA1 by lentiviral sgRNA transduction of iPSCs prior to proceeding with neuron differentiation (Figure 5A). Western blots and immunofluorescence both confirm RBP KDs compared to a non-targeting (NT) control guide (Supplementary Figure S4A-D). We then fractionated RBP KD neurons and performed proteomics. Most of the variance between samples can be explained by the neurite versus whole cell separation, rather than RBP KD (Supplementary Figure S4E-F). We observe decreased abundance of the knocked down RBP in both the neuritic and whole cell fractions (Figure 5B-E), confirming the efficacy of our knockdowns. TDP-43 KD leads to decreased expression of more proteins in neurites (Figure 5B), while hnRNPA1 KD leads to increased expression of more proteins in neurites compared to those with decreased expression (Figure 5C). For both TDP-43 and hnRNPA1 KD, we observe fewer protein changes in the whole cell fraction than the neurites, suggesting that loss of either of these proteins mostly affects neuritic proteins (Figure 5D-E). We did not observe many proteins that were inversely affected in each compartment (decrease in neurite and corresponding increase in whole cell, or vice versa). Pathway analysis identified pathways related to the extracellular matrix in TDP-43 KD neurites and RNA processing in hnRNPA1 KD neurites (Supplementary Figure S4G-H). Surprisingly, for hnRNPA1 KD, we observed an increase in the related protein hnRNPA3 in both cellular compartments (Figure 5C,E). This increase suggests that hnRNPA3 may be compensating for the loss of expression of hnRNPA1. Indeed, KD of either hnRNPA1 or hnRNPA3 followed by immunofluorescence shows an approximately 50% increased protein level of the opposite protein (Figure 5F-H). Of note, hnRNPA1 has been shown to have a similar compensation mechanism with hnRNAP2 (52), suggesting these compensatory mechanisms may be more prevalent or complex than one protein compensating for another.

**Figure 5:**
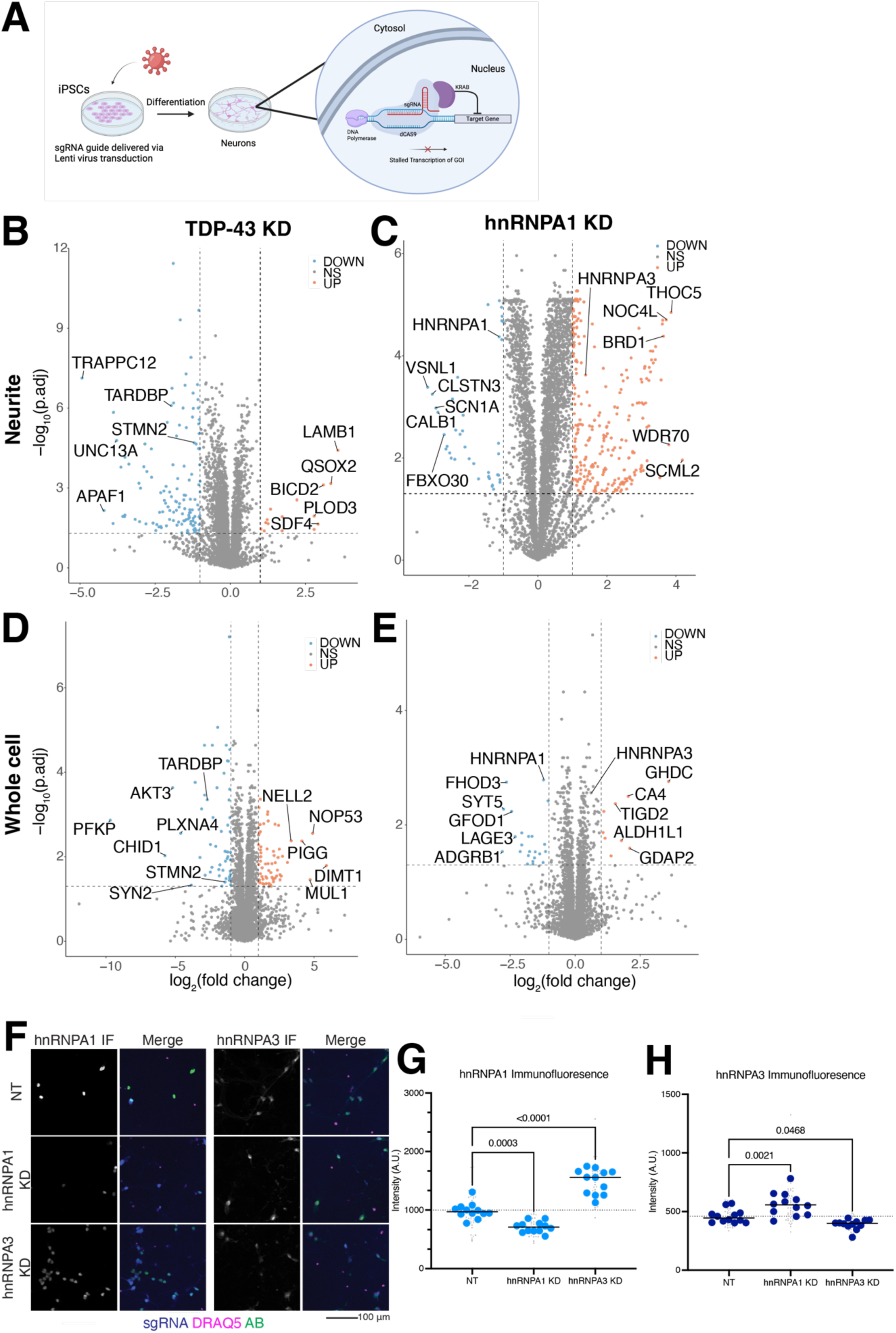
FTD/ALS-associated RBP KD causes moderate changes to the neuritic proteome, potentially modulated by compensation. A. Knockdowns are performed by lentiviral transduction of sgRNA into iPSCs expressing dCas9-KRAB, preventing transcription by binding to the promoter of the gene of interest. B. Volcano plot of TDP-43 KD vs NT proteins in the neurites shows a small number of proteins that are increased in neurites (right, orange) but a larger number decreased (left, blue), including known cryptic-exon containing genes STMN2 and UNC13A. Vertical dotted lines indicate a log2(fold change) of 1 or −1, horizontal dotted line indicates a p value of 0.05. n=6 samples each condition. C. Volcano plot of hnRNPA1 KD vs NT proteins in the neurites shows many proteins with increased abundance in neurites (right, orange) and a smaller number decreased (left, blue). The related hnRNP, hnRNPA3 is increased. Vertical dotted lines indicate a log2(fold change) of 1 or −1, horizontal dotted line indicates a p value of 0.05. n=5 NT, n=6 KD. D. Volcano plot of TDP-43 KD vs NT proteins in the whole cell fraction shows more proteins that are increased in whole cell compared to the neurite fraction (right, orange) but still a larger number of proteins with decreased abundance (left, blue), including known cryptic-exon containing gene STMN2. Vertical dotted lines indicate a log2(fold change) of 1 or −1, horizontal dotted line indicates a p value of 0.05. n=5 samples each condition. E. Volcano plot of hnRNPA1 KD vs NT proteins in the whole cell fraction shows fewer proteins with increased abundance than was observed in neurites (right, orange) and a similarly small number of proteins with decreased abundance (left, blue). Vertical dotted lines indicate a log2(fold change) of 1 or −1, horizontal dotted line indicates a p value of 0.05. n=6 samples each condition. F. Representative images of immunofluorescence (IF) of hnRNPA1 and hnRNPA3 levels in NT and KD neurons. sgRNA expression is visualized by co-expressed cytosolic BFP signal, immunofluorescence for the RBP of interest in green, and DRAQ5 nuclear stain in magenta. Scale bar 100 µm. G. Quantification of hnRNPA1 IF microscopy shows increased hnRNPA3 IF signal in hnRNPA1 KD neurons, suggesting compensatory changes in RBP levels upon knockdown. n=12 wells (large blue dots), 4 images per well (small gray dots). p-values from one-way ANOVA (p-value <0.0001) corrected by Šídák’s multiple comparisons test. H. Quantification of hnRNPA3 IF microscopy shows a slight increase in hnRNPA1 IF signal in hnRNPA3 KD neurons, suggesting compensatory changes in RBP levels upon knockdown. n=12 wells (large blue dots), 4 images per well (small gray dots). p-values from one-way ANOVA (p-value <0.0001) corrected by Šídák’s multiple comparisons test.

### RBP KD does not induce RNA mislocalization

To determine the effect of RBP KD on mRNA localization in our iPSC-derived neurons, we performed RNA sequencing on fractionated NT and KD neurons. Overall, samples show similar percent expression efficiencies, but neurite samples tend to have higher percentages of reads aligning to ribosomal RNA (rRNA) (Supplementary Figure S5A-B). Samples had similar numbers of genes detected within an experiment, although the hnRNPA1 experiment typically had more genes detected (Supplementary Figure S5C-D). To confirm separation of neurites from whole cell, we looked for expression of small nucleolar RNAs (snoRNAs) in all samples. As expected, higher snoRNA expression was observed in whole cell samples, but some snoRNAs and pre-mRNAs were detected in neurites, with more detected in the hnRNPA1 experiment compared to the TDP-43 experiment (Supplementary Figure S5E-F). Although some level of cross-contamination between compartments can occur in such experiments and explain our findings, snoRNAs are still increased in the whole cell fractions, supporting the validity of this dataset analysis.

Transcriptomics further validates our KDs, as both RBPs are significantly reduced in both the neuritic and whole cell fractions (Figure 6A-D). Further, as expected for decreased TDP-43 levels, STMN2 mRNA levels are significantly reduced in both fractions (Figure 6A,C) (53). Overall, TDP- 43 KD shows the most genes with expression changes in both compartments (Figure 6A,C). hnRNPA1 KD shows a more moderate effect on the neuritic and whole cell transcriptome, with some genes changing expression level in each compartment; interestingly, more genes have increased expression than decreased expression (Figure 6B,D). Interestingly, most of the difference in the samples (81% and 79% for TDP-43 KD and hnRNPA1 KD, respectively) comes from the cellular fraction, not the KD itself (Supplementary Figure S6A-B). Pathway analysis identifies axon guidance related transcripts decreased in TDP-43 KD neurons and neurodegeneration transcripts in hnRNPA1 KD neurons (Supplementary Figure S6C-E).

**Figure 6:**
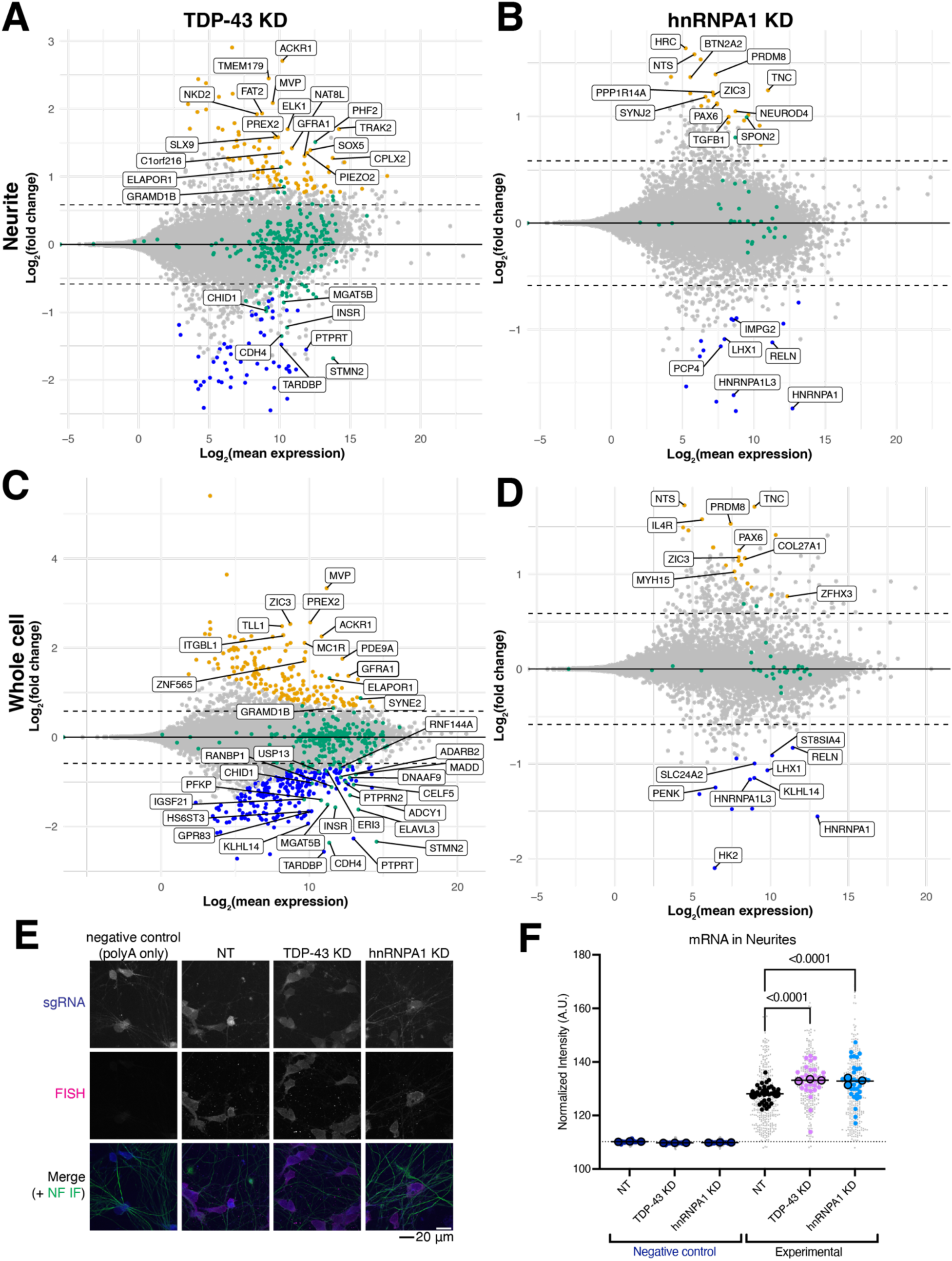
RBP KD does not drastically alter mRNA localization but increases total neuritic mRNA. A. MA plot comparing TDP-43 KD to NT for the neuritic fraction. 98 genes show increased expression while 59 show decreased expression, including TDP-43 and STMN2. log2(1.5) fold change cutoff. n=5 for each KD and NT. Genes that contain cryptic exons upon TDP- 43 KD are highlighted in green. B. MA plot comparing hnRNPA1 KD to NT for the neuritic fraction. 25 genes show increased expression while 16 show decreased expression, including hnRNPA1. log2(1.5) fold change cutoff. n=6 for each KD and NT. Genes that contain cryptic exons upon hnRNPA1 KD are highlighted in green. C. MA plot comparing TDP-43 KD to NT for the whole cell fraction. 179 genes show increased expression while 240 show decreased expression, including TDP-43 and STMN2. log2(1.5) fold change cutoff. n=5 for each KD and NT. Genes that contain cryptic exons upon TDP-43 KD are highlighted in green. D. MA plot comparing hnRNPA1 KD to NT for the whole cell fraction. 21 genes show increased expression while 13 show decreased expression, including hnRNPA1. log2(1.5) fold change cutoff. n=6 for each KD and NT. Genes that contain cryptic exons upon hnRNPA1 KD are highlighted in green. E. Representative images of polyA total mRNA FISH of NT and RBP KD neurons. mRNA is in magenta, anti-NEFL staining (axons) in green, and sgRNA in blue. Scale bar 20 µm. F. Quantification of polyA total mRNA FISH. TDP-43 and hnRNPA1 KD have increased neuritic mRNA compared to NT. Background signal is indicated by dotted line, from the non-fluorescent negative control. Neurites (NEFL) expressing sgRNA were used to mask FISH signal prior to intensity quantification. n=3 differentiations (replicates, means for each replicate indicated as large, outlined dots, with a line representing overall mean of condition), 12 wells per condition (mean of each well shown as a medium dot), 12 images per well (small gray dots). P-value from one-way ANOVA of replicate values (overall p < 0.0001), graph shows p-values corrected for multiple comparisons.

Loss of TDP-43 causes many changes to alternative splicing, like the inclusion of exons that are not normally expressed, called “cryptic” exons (54,55). hnRNPA1 is also involved in alternative splicing and altered splicing was recently found in a mouse model of hnRNPA1 mislocalization- associated multiple sclerosis (56). As such, we simultaneously analyzed our whole cell transcriptomics datasets for novel splice junctions. We observe over 500 novel splice junctions in the TDP-43 KD dataset and almost 150 in the hnRNPA1 KD neurons (Table S1 and 2). While most genes containing cryptic exons do not have significantly changed RNA levels in our transcriptomics for either TDP-43 or hnRNPA1 KD, a small number do. Interestingly, the only genes containing cryptic exons for hnRNPA1 KD that show significantly changed mRNA levels are increased in the neuritic fraction (Figure 6B). In contrast, genes containing cryptic exons in TDP-43 KD more often decrease mRNA levels in either compartment and these changes are consistent across compartment (Figure 6A,C).

To assess whether RBP KD alters total mRNA levels in neurites, we performed FISH against polyA tails on the NT and KD neurons. Quantification shows increased polyA transcripts in the neurites within TDP-43 and hnRNPA1 KD (Figure 6E-F). This increased RNA amount in the neurites, coupled with the differences in protein expression from proteomics, suggested to us that perhaps there was a defect in local translation in these knockdown neurites.

### Increased neuritic translation but decreased neurite outgrowth in TDP-43 KD

To address whether RBP KD alters neuritic translation, we used fluorescent non canonical amino acid tagging (FUNCAT) (21,57). FUNCAT is analogous to QuanCAT but uses click chemistry to attach a fluorophore to AHA after it is incorporated into newly synthesized proteins (Figure 7A). We confirmed the specificity of FUNCAT using methionine (Met) as a negative control and found decreased FUNCAT signal in cells treated with the protein synthesis inhibitors cycloheximide (CHX) and anisomycin (Aniso) (Figure 7B-C). Interestingly, TDP-43 KD shows increased translation compared to a NT control, but hnRNPA1 KD does not (Figure 7B-C). Given this increased translation and knowing that loss of TDP-43 can lead to decreased neurite outgrowth in other systems (58,59), likely due to decreased STMN2 (60), we wondered if our iNeurons also show decreased neurite outgrowth. We measured neurite outgrowth of KD neurons grown as neurospheres (Figure 7D-E). Quantification of the neurospheres show that across the 5 timepoints, our TDP-43 KD spheres show a significantly smaller size from our NT neurons. Additionally, hnRNPA1 KD remained significantly smaller than NT spheres throughout all time points tested (Figure 7D-E). However, the sphere area did not change in size substantially across the time course for any genotype, indicating neurons are not dying (Figure S7A). These results were confirmed by a two-dimensional neurite outgrowth assay, where 1% of cells were labeled with a cytoplasmic marker (Supplementary Figure S7B-F). These neurite outgrowth defects in TDP-43 and hnRNPA1 KD neurons demonstrate that these changes observed in the local transcriptome and proteome are correlated with functional deficits.

**Figure 7:**
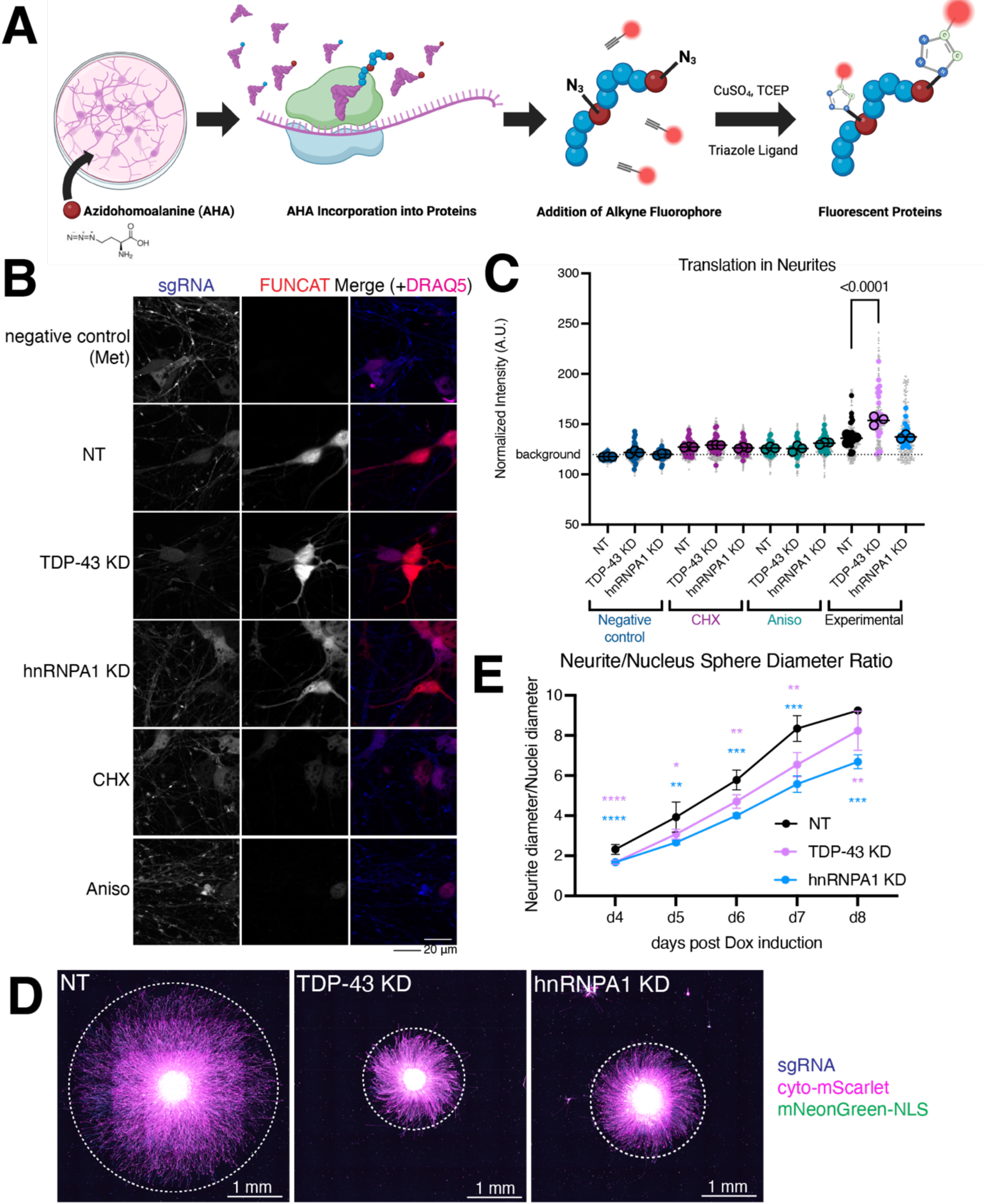
RBP KD alters neuritic translation and neurite outgrowth. A. Schematic outlining fluorescent non-canonical amino acid tagging (FUNCAT). Cells are starved of methionine before adding methionine analog azidohomoalanine (AHA). AHA is incorporated into new proteins, which, after fixation, is detected by addition of a fluorophore through click chemistry. B. Representative FUNCAT images. sgRNA in blue, FUNCAT signal in red, nuclear stain (DRAQ5) in magenta. Scale bar 20 µm. C. Quantification of FUNCAT. TDP-43 has increased translation compared to NT. Background signal is indicated by dotted line (from methionine negative control, Met). Protein synthesis inhibitors anisomycin (Aniso) and cycloheximide (CHX) decrease FUNCAT signal. FUNCAT signal masked by sgRNA prior to intensity quantification. n=3 differentiations (replicates, means for each replicate indicated as large, outlined dots, overall mean indicated by line), 12 wells (medium dots), 12 images per well (small grey dots). p-value from one-way ANOVA of replicate values (overall p < 0.0001), graph shows significant p-value corrected for multiple comparisons. D. Representative micrographs of NT and RBP KD neurospheres 5 days after dox induction. Dashed lines indicate the extent of neurite outgrowth. sgRNA in blue, cytoplasmic mScarlet marker in magenta, and nuclear mNeonGreen in green. Scale bar 1 mm. E. Quantification of neurite diameter (red channel) divided by cell body diameter (green channel) indicates decreased neurite outgrowth in TDP-43 and hnRNPA1 KD. n ≥ 3 spheres per genotype per day. Significant days indicated by stars, **** p<0.0001, *** 0.001 < p < 0.0001, ** 0.01 < p <0.001. n=3-10 for NT, 10-13 for TDP-43 KD, and 9 or 10 for hnRNPA1 KD. Number of spheres vary because spheres were removed from analysis when they touched the edge of the well. P-values from mixed-effects analysis (rather than ANOVA due to the inconsistent number of spheres per day) with Dunnett’s multiple comparison test.

## Discussion

mRNA transport and local translation are crucial processes that enable neurons to respond rapidly to synaptic stimuli and are crucial for maintaining neuronal health. We found thousands of transcripts enriched in neurites, but fewer neurite-enriched proteins. Indeed, many of the transcripts we identified in neurites were surprising, like the ribosomal protein subunits, which may be explained by local ribosome remodeling (61). The presence of other transcripts encoding other highly nuclear enriched proteins in the neurites, like hnRNPA1, is harder to explain and may lead one to believe that mRNAs are stochastically transported to neurites. However, our data indicate that, generally, neuritic transcripts are shorter and use the most proximal 3’ UTR, thus identifying a potential sorting mechanism for the cell to identify transcripts that should be transported to neurites that requires more work to validate. Interestingly, shorter mRNAs are more likely to be translated (48), which might explain why neuritic transcripts tend to be shorter, due to the energy the cell must expend to transport these transcripts. Indeed, many neuritic transcripts are detected as also newly synthesized in neurites and may be among the transcripts that are most often translated in neurons. There are likely more specific sorting mechanisms for transcripts that are transported to neurites from those that remain in the cell body, but more work is needed to identify them. Additionally, the most highly enriched neuritic transcripts are not necessarily the most highly translated proteins in neurites, so more work is needed to understand why some transcripts are translated more than others.

RBPs are required for the transport and translation of mRNAs in neuronal processes. Our work provides some evidence that the RBPs involved in mRNA transport or staying in the cell body might be different, as we observe binding sites for RBM8A enriched in neuritic transcripts but SRSF7 binding sites on transcripts enriched in the cell body. However, these are unlikely to be the only RBPs regulating this difference, or perhaps there are other, non-RNA binding, interacting proteins contributing to this difference that still need to be identified. Loss of FTD/ALS-associated RBPs TDP-43 and hnRNPA1 alter the local transcriptome and proteome in neurites. We find increased total transcripts in TDP-43 KD neurites, but fewer genes with altered expression in the neurites than the whole cell fraction. Conversely, we observe fewer changes in local transcriptome in hnRNPA1 KD but an increase in total neuritic mRNA, without a similar increase in local translation. However, for hnRNPA1 KD, we observed a compensatory increase in protein levels of hnRNPA3, a protein within the same family that binds to mRNA similarly (62), which may explain the moderate changes in the local transcriptome. This increase was only found at the protein level, we did not observe a similar change in transcriptomics. This protein-specific dose compensation suggests homeostatic changes to either the translation or protein degradation systems, potentially regulated by N-acetyltransferases (63). More work is needed to understand the role of protein degradation and associated posttranslational modifications in maintaining neuritic protein levels.

TDP-43 loss is well known to induce alternative splicing events, leading to inclusion of “cryptic” exons (54,55). Recently, some of these cryptic exons have been observed to be translated into protein products, paving the way for their use as potential biomarkers of disease (64,65). The identification of cryptic exons in hnRNPA1 KD neurons suggests that cryptic exons may form due to dysfunction in other RBPs besides TDP-43 in FTD/ALS and are not a TDP-43-specific phenomenon (66–68). As such, other diseases, like multiple sclerosis (56), may also benefit from biomarker identification using the pipelines developed for TDP-43 (64,65). In addition to the mRNA level changes observed in both compartments due to cryptic exon inclusion, we observed increases in both mRNA and translation levels in TDP-43 KD neurites and mRNA only in hnRNPA1 KD neurites. Both intracellular transport and translation are highly regulated cellular processes, and these increases may be the neuron attempting to compensate for altered mRNA transport and translation. The decreased neurite outgrowth observed in both TDP-43 and hnRNPA1 KD neurons suggests that neurons are not able to effectively compensate for these changes. In conclusion, these results provide transcriptomic and proteomic insights into mRNA transport and local translation in iPSC-derived neurons, as well as the effect of losing two FTD/ALS-associated RBPs, TDP-43 and hnRNPA1 on these processes.

## Data Availability

RNA sequencing and proteomics data files can be accessed at Alzheimer’s Disease Workbench (ADWB): https://fair.addi.addatainitiative.org/#/data/datasets/neurite_and_whole_cell_transcriptomics_on_ineurons. The mass spectrometry proteomics data have been deposited to the ProteomeXchange Consortium via the PRIDE partner repository with the dataset identifier PXD054548 (69). Code and processed transcriptomics and proteomics data are available at https://github.com/NIH-CARD/Neurite-WT-RBP-KD.

## Acknowledgements

We thank Kendall Van Keuren-Jenson, Mark Cookson, Michael Ward, and Miryam Gorospe for helpful comments. We thank the Sequencing Cores at the National Heart, Lung, and Blood Institute and the National Institute on Childhood Health and Development for performing RNA sequencing. We also thank the Proteomics Core at the National Institute of Neurological Disorders and Stroke for performing some of the mass spec.

## References

1. Idler, R.K. and Yan, W. (2012) Control of messenger RNA fate by RNA-binding proteins: an emphasis on mammalian spermatogenesis. J Androl, 33, 309–337.

2. Hentze, M.W., Castello, A., Schwarzl, T. and Preiss, T. (2018) A brave new world of RNA-binding proteins. Nat Rev Mol Cell Biol, 19, 327–341.

3. Donlin-Asp, P.G., Polisseni, C., Klimek, R., Heckel, A. and Schuman, E.M. (2021) Differential regulation of local mRNA dynamics and translation following long-term potentiation and depression. Proc Natl Acad Sci U S A, 118.

4. Tiruchinapalli, D.M., Oleynikov, Y., Kelic, S., Shenoy, S.M., Hartley, A., Stanton, P.K., Singer, R.H. and Bassell, G.J. (2003) Activity-dependent trafficking and dynamic localization of zipcode binding protein 1 and beta-actin mRNA in dendrites and spines of hippocampal neurons. J Neurosci, 23, 3251–3261.

5. Bassell, G.J., Zhang, H., Byrd, A.L., Femino, A.M., Singer, R.H., Taneja, K.L., Lifshitz, L.M., Herman, I.M. and Kosik, K.S. (1998) Sorting of beta-actin mRNA and protein to neurites and growth cones in culture. J Neurosci, 18, 251–265.

6. Zappulo, A., van den Bruck, D., Ciolli Mattioli, C., Franke, V., Imami, K., McShane, E., Moreno-Estelles, M., Calviello, L., Filipchyk, A., Peguero-Sanchez, E., et al. (2017) RNA localization is a key determinant of neurite-enriched proteome. Nat Commun, 8, 583.

7. Ling, S.C., Polymenidou, M. and Cleveland, D.W. (2013) Converging mechanisms in ALS and FTD: disrupted RNA and protein homeostasis. Neuron, 79, 416–438.

8. Wood, A., Gurfinkel, Y., Polain, N., Lamont, W. and Lyn Rea, S. (2021) Molecular Mechanisms Underlying TDP-43 Pathology in Cellular and Animal Models of ALS and FTLD. Int J Mol Sci, 22.

9. Kim, H.J., Kim, N.C., Wang, Y.D., Scarborough, E.A., Moore, J., Diaz, Z., MacLea, K.S., Freibaum, B., Li, S., Molliex, A. et al. (2013) Mutations in prion-like domains in hnRNPA2B1 and hnRNPA1 cause multisystem proteinopathy and ALS. Nature, 495, 467–473.

10. Mehta, P.R., Brown, A.L., Ward, M.E. and Fratta, P. (2023) The era of cryptic exons: implications for ALS-FTD. Mol Neurodegener, 18, 16.

11. Alami, N.H., Smith, R.B., Carrasco, M.A., Williams, L.A., Winborn, C.S., Han, S.S.W., Kiskinis, E., Winborn, B., Freibaum, B.D., Kanagaraj, A. et al. (2014) Axonal transport of TDP-43 mRNA granules is impaired by ALS-causing mutations. Neuron, 81, 536–543.

12. Imperatore, J.A., McAninch, D.S., Valdez-Sinon, A.N., Bassell, G.J. and Mihailescu, M.R. (2020) FUS Recognizes G Quadruplex Structures Within Neuronal mRNAs. Front Mol Biosci, 7, 6.

13. Burguete, A.S., Almeida, S., Gao, F.B., Kalb, R., Akins, M.R. and Bonini, N.M. (2015) GGGGCC microsatellite RNA is neuritically localized, induces branching defects, and perturbs transport granule function. Elife, 4, e08881.

14. Wang, C., Ward, M.E., Chen, R., Liu, K., Tracy, T.E., Chen, X., Xie, M., Sohn, P.D., Ludwig, C., Meyer-Franke, A. et al. (2017) Scalable Production of iPSC-Derived Human Neurons to Identify Tau-Lowering Compounds by High-Content Screening. Stem Cell Reports, 9, 1221–1233.

15. Fernandopulle, M.S., Prestil, R., Grunseich, C., Wang, C., Gan, L. and Ward, M.E. (2018) Transcription Factor-Mediated Differentiation of Human iPSCs into Neurons. Curr Protoc Cell Biol, 79, e51.

16. Tian, R., Gachechiladze, M.A., Ludwig, C.H., Laurie, M.T., Hong, J.Y., Nathaniel, D., Prabhu, A.V., Fernandopulle, M.S., Patel, R., Abshari, M. et al. (2019) CRISPR Interference-Based Platform for Multimodal Genetic Screens in Human iPSC-Derived Neurons. Neuron, 104, 239–255 e212.

17. Burgold, T., Karakoc, E., Gonçalves, E., Dwane, L., Barrio-Hernandez, I., Silva, R.O., Souster, E., Sharma, M., Beck, A., Koh, G. et al. (2024) Genetic interaction library screening with a next-generation dual guide CRISPR system. bioRxiv, 2024.2003.2028.587052.

18. Reilly, L., Lara, E., Ramos, D., Li, Z., Pantazis, C.B., Stadler, J., Santiana, M., Roberts, J., Faghri, F., Hao, Y. et al. (2023) A fully automated FAIMS-DIA mass spectrometry-based proteomic pipeline. Cell Rep Methods, 3, 100593.

19. Carpenter, A.E., Jones, T.R., Lamprecht, M.R., Clarke, C., Kang, I.H., Friman, O., Guertin, D.A., Chang, J.H., Lindquist, R.A., Moffat, J. et al. (2006) CellProfiler: image analysis software for identifying and quantifying cell phenotypes. Genome Biol, 7, R100.

20. Tsanov, N., Samacoits, A., Chouaib, R., Traboulsi, A.M., Gostan, T., Weber, C., Zimmer, C., Zibara, K., Walter, T., Peter, M. et al. (2016) smiFISH and FISH-quant - a flexible single RNA detection approach with super-resolution capability. Nucleic Acids Res, 44, e165.

21. Tom Dieck, S., Muller, A., Nehring, A., Hinz, F.I., Bartnik, I., Schuman, E.M. and Dieterich, D.C. (2012) Metabolic labeling with noncanonical amino acids and visualization by chemoselective fluorescent tagging. Curr Protoc Cell Biol, **Chapter** 7, 711 11 17–11 29.

22. Dobin, A., Davis, C.A., Schlesinger, F., Drenkow, J., Zaleski, C., Jha, S., Batut, P., Chaisson, M. and Gingeras, T.R. (2013) STAR: ultrafast universal RNA-seq aligner. Bioinformatics, 29, 15–21.

23. Danecek, P., Bonfield, J.K., Liddle, J., Marshall, J., Ohan, V., Pollard, M.O., Whitwham, A., Keane, T., McCarthy, S.A., Davies, R.M. (2021) Twelve years of SAMtools and BCFtools. Gigascience, 10.

24. Liao, Y., Smyth, G.K. and Shi, W. (2014) featureCounts: an efficient general purpose program for assigning sequence reads to genomic features. Bioinformatics, 30, 923–930.

25. Love, M.I., Huber, W. and Anders, S. (2014) Moderated estimation of fold change and dispersion for RNA-seq data with DESeq2. Genome Biol, 15, 550.

26. Yu, G., Wang, L.G., Han, Y. and He, Q.Y. (2012) clusterProfiler: an R package for comparing biological themes among gene clusters. OMICS, 16, 284–287.

27. Wu, T., Hu, E., Xu, S., Chen, M., Guo, P., Dai, Z., Feng, T., Zhou, L., Tang, W., Zhan, L. et al. (2021) clusterProfiler 4.0: A universal enrichment tool for interpreting omics data. Innovation (Camb), 2, 100141.

28. Subramanian, A., Tamayo, P., Mootha, V.K., Mukherjee, S., Ebert, B.L., Gillette, M.A., Paulovich, A., Pomeroy, S.L., Golub, T.R., Lander, E.S. (2005) Gene set enrichment analysis: a knowledge-based approach for interpreting genome-wide expression profiles. Proc Natl Acad Sci U S A, 102, 15545–15550.

29. Liberzon, A., Subramanian, A., Pinchback, R., Thorvaldsdottir, H., Tamayo, P. and Mesirov, J.P. (2011) Molecular signatures database (MSigDB) 3.0. Bioinformatics, 27, 1739–1740.

30. Ashburner, M., Ball, C.A., Blake, J.A., Botstein, D., Butler, H., Cherry, J.M., Davis, A.P., Dolinski, K., Dwight, S.S., Eppig, J.T. et al. (2000) Gene ontology: tool for the unification of biology. The Gene Ontology Consortium. Nat Genet, 25, 25–29.

31. Gene Ontology, C., Aleksander, S.A., Balhoff, J., Carbon, S., Cherry, J.M., Drabkin, H.J., Ebert, D., Feuermann, M., Gaudet, P., Harris, N.L., et al. (2023) The Gene Ontology knowledgebase in 2023. Genetics, 224.

32. Vaquero-Garcia, J., Aicher, J.K., Jewell, S., Gazzara, M.R., Radens, C.M., Jha, A., Norton, S.S., Lahens, N.F., Grant, G.R. and Barash, Y. (2023) RNA splicing analysis using heterogeneous and large RNA-seq datasets. Nat Commun, 14, 1230.

33. Vitting-Seerup, K. and Sandelin, A. (2019) IsoformSwitchAnalyzeR: analysis of changes in genome-wide patterns of alternative splicing and its functional consequences. Bioinformatics, 35, 4469–4471.

34. Patro, R., Duggal, G., Love, M.I., Irizarry, R.A. and Kingsford, C. (2017) Salmon provides fast and bias-aware quantification of transcript expression. Nat Methods, 14, 417–419.

35. Ha, K.C.H., Blencowe, B.J. and Morris, Q. (2018) QAPA: a new method for the systematic analysis of alternative polyadenylation from RNA-seq data. Genome Biol, 19, 45.

36. Kolberg, L., Raudvere, U., Kuzmin, I., Adler, P., Vilo, J. and Peterson, H. (2023) g:Profiler-interoperable web service for functional enrichment analysis and gene identifier mapping (2023 update). Nucleic Acids Res, 51, W207–W212.

37. Krismer, K., Bird, M.A., Varmeh, S., Handly, E.D., Gattinger, A., Bernwinkler, T., Anderson, D.A., Heinzel, A., Joughin, B.A., Kong, Y.W. et al. (2020) Transite: A Computational Motif-Based Analysis Platform That Identifies RNA-Binding Proteins Modulating Changes in Gene Expression. Cell Rep, 32, 108064.

38. Choi, Y., Um, B., Na, Y., Kim, J., Kim, J.S. and Kim, V.N. (2024) Time-resolved profiling of RNA binding proteins throughout the mRNA life cycle. Mol Cell, 84, 1764–1782 e1710.

39. Li, Z., Weller, C.A., Shah, S., Johnson, N., Hao, Y., Roberts, J., Bereda, C., Klaisner, S., Machado, P., Fratta, P. et al. (2023) ProtPipe: A Multifunctional Data Analysis Pipeline for Proteomics and Peptidomics. bioRxiv.

40. Demichev, V., Messner, C.B., Vernardis, S.I., Lilley, K.S. and Ralser, M. (2020) DIA-NN: neural networks and interference correction enable deep proteome coverage in high throughput. Nat Methods, 17, 41–44.

41. Willis, D.E. and Twiss, J.L. (2011) Profiling axonal mRNA transport. Methods Mol Biol, 714, 335–352.

42. Baas, P.W. and Lin, S. (2011) Hooks and comets: The story of microtubule polarity orientation in the neuron. Dev Neurobiol, 71, 403–418.

43. Sproviero, W., Shatunov, A., Stahl, D., Shoai, M., van Rheenen, W., Jones, A.R., Al-Sarraj, S., Andersen, P.M., Bonini, N.M., Conforti, F.L.,et al. (2017) ATXN2 trinucleotide repeat length correlates with risk of ALS. Neurobiol Aging, 51, 178 e171–178 e179.

44. Ganguly, A., Feldman, R.M. and Guo, M. (2008) ubiquilin antagonizes presenilin and promotes neurodegeneration in Drosophila. Hum Mol Genet, 17, 293–302.

45. Wang, S., Tatman, M. and Monteiro, M.J. (2020) Overexpression of UBQLN1 reduces neuropathology in the P497S UBQLN2 mouse model of ALS/FTD. Acta Neuropathol Commun, 8, 164.

46. Bae, B. and Miura, P. (2020) Emerging Roles for 3’ UTRs in Neurons. Int J Mol Sci, 21.

47. Andreassi, C., Luisier, R., Crerar, H., Darsinou, M., Blokzijl-Franke, S., Lenn, T., Luscombe, N.M., Cuda, G., Gaspari, M., Saiardi, A. (2021) Cytoplasmic cleavage of IMPA1 3’ UTR is necessary for maintaining axon integrity. Cell Rep, 34, 108778.

48. Middleton, S.A., Eberwine, J. and Kim, J. (2019) Comprehensive catalog of dendritically localized mRNA isoforms from sub-cellular sequencing of single mouse neurons. BMC Biol, 17, 5.

49. Alachkar, A., Jiang, D., Harrison, M., Zhou, Y., Chen, G. and Mao, Y. (2013) An EJC factor RBM8a regulates anxiety behaviors. Curr Mol Med, 13, 887–899.

50. Howden, A.J., Geoghegan, V., Katsch, K., Efstathiou, G., Bhushan, B., Boutureira, O., Thomas, B., Trudgian, D.C., Kessler, B.M., Dieterich, D.C. et al. (2013) QuaNCAT: quantitating proteome dynamics in primary cells. Nat Methods, 10, 343–346.

51. Hein, C.D., Liu, X.M. and Wang, D. (2008) Click chemistry, a powerful tool for pharmaceutical sciences. Pharm Res, 25, 2216–2230.

52. Chang, Y., Lu, X. and Qiu, J. (2021) Compensatory expression regulation of highly homologous proteins HNRNPA1 and HNRNPA2. Turk J Biol, 45, 187–195.

53. Prudencio, M., Humphrey, J., Pickles, S., Brown, A.L., Hill, S.E., Kachergus, J.M., Shi, J., Heckman, M.G., Spiegel, M.R., Cook, C. et al. (2020) Truncated stathmin-2 is a marker of TDP-43 pathology in frontotemporal dementia. J Clin Invest, 130, 6080–6092.

54. Arnold, E.S., Ling, S.C., Huelga, S.C., Lagier-Tourenne, C., Polymenidou, M., Ditsworth, D., Kordasiewicz, H.B., McAlonis-Downes, M., Platoshyn, O., Parone, P.A. et al. (2013) ALS-linked TDP-43 mutations produce aberrant RNA splicing and adult-onset motor neuron disease without aggregation or loss of nuclear TDP-43. Proc Natl Acad Sci U S A, 110, E736–745.

55. Highley, J.R., Kirby, J., Jansweijer, J.A., Webb, P.S., Hewamadduma, C.A., Heath, P.R., Higginbottom, A., Raman, R., Ferraiuolo, L., Cooper-Knock, J. et al. (2014) Loss of nuclear TDP-43 in amyotrophic lateral sclerosis (ALS) causes altered expression of splicing machinery and widespread dysregulation of RNA splicing in motor neurones. Neuropathol Appl Neurobiol, 40, 670–685.

56. Salapa, H.E., Thibault, P.A., Libner, C.D., Ding, Y., Clarke, J.W.E., Denomy, C., Hutchinson, C., Abidullah, H.M., Austin Hammond, S., Pastushok, L. et al. (2024) hnRNP A1 dysfunction alters RNA splicing and drives neurodegeneration in multiple sclerosis (MS). Nat Commun, 15, 356.

57. Dieterich, D.C., Hodas, J.J., Gouzer, G., Shadrin, I.Y., Ngo, J.T., Triller, A., Tirrell, D.A. and Schuman, E.M. (2010) In situ visualization and dynamics of newly synthesized proteins in rat hippocampal neurons. Nat Neurosci, 13, 897–905.

58. Fiesel, F.C., Schurr, C., Weber, S.S. and Kahle, P.J. (2011) TDP-43 knockdown impairs neurite outgrowth dependent on its target histone deacetylase 6. Mol Neurodegener, 6, 64.

59. Tripathi, V.B., Baskaran, P., Shaw, C.E. and Guthrie, S. (2014) Tar DNA-binding protein- 43 (TDP-43) regulates axon growth in vitro and in vivo. Neurobiol Dis, 65, 25–34.

60. Lopez-Erauskin, J., Bravo-Hernandez, M., Presa, M., Baughn, M.W., Melamed, Z., Beccari, M.S., Agra de Almeida Quadros, A.R., Arnold-Garcia, O., Zuberi, A., Ling, K., et al. (2024) Stathmin-2 loss leads to neurofilament-dependent axonal collapse driving motor and sensory denervation. Nat Neurosci, 27, 34–47.

61. Shigeoka, T., Koppers, M., Wong, H.H., Lin, J.Q., Cagnetta, R., Dwivedy, A., de Freitas Nascimento, J., van Tartwijk, F.W., Strohl, F., Cioni, J.M., et al. (2019) On-Site Ribosome Remodeling by Locally Synthesized Ribosomal Proteins in Axons. Cell Rep, 29, 3605–3619 e3610.

62. Lu, Y., Wang, X., Gu, Q., Wang, J., Sui, Y., Wu, J. and Feng, J. (2022) Heterogeneous nuclear ribonucleoprotein A/B: an emerging group of cancer biomarkers and therapeutic targets. Cell Death Discov, 8, 337.

63. Ishikawa, K., Ishihara, A. and Moriya, H. (2020) Exploring the Complexity of Protein-Level Dosage Compensation that Fine-Tunes Stoichiometry of Multiprotein Complexes. PLoS Genet, 16, e1009091.

64. Seddighi, S., Qi, Y.A., Brown, A.L., Wilkins, O.G., Bereda, C., Belair, C., Zhang, Y.J., Prudencio, M., Keuss, M.J., Khandeshi, A. et al. (2024) Mis-spliced transcripts generate de novo proteins in TDP-43-related ALS/FTD. Sci Transl Med, 16, eadg7162.

65. Irwin, K.E., Jasin, P., Braunstein, K.E., Sinha, I.R., Garret, M.A., Bowden, K.D., Chang, K., Troncoso, J.C., Moghekar, A., Oh, E.S. et al. (2024) A fluid biomarker reveals loss of TDP-43 splicing repression in presymptomatic ALS-FTD. Nat Med, 30, 382–393.

66. Tan, Q., Yalamanchili, H.K., Park, J., De Maio, A., Lu, H.C., Wan, Y.W., White, J.J., Bondar, V.V., Sayegh, L.S., Liu, X., et al. (2016) Extensive cryptic splicing upon loss of RBM17 and TDP43 in neurodegeneration models. Hum Mol Genet, 25, 5083–5093.

67. Ehrmann, I., Crichton, J.H., Gazzara, M.R., James, K., Liu, Y., Grellscheid, S.N., Curk, T., de Rooij, D., Steyn, J.S., Cockell, S., et al. (2019) An ancient germ cell-specific RNA-binding protein protects the germline from cryptic splice site poisoning. Elife, 8.

68. Zheng, R., Dunlap, M., Bobkov, G.O.M., Gonzalez-Figueroa, C., Patel, K.J., Lyu, J., Harvey, S.E., Chan, T.W., Quinones-Valdez, G., Choudhury, M. et al. (2024) hnRNPM protects against the dsRNA-mediated interferon response by repressing LINE-associated cryptic splicing. Mol Cell, 84, 2087–2103 e2088.

69. Perez-Riverol, Y., Bai, J., Bandla, C., Garcia-Seisdedos, D., Hewapathirana, S., Kamatchinathan, S., Kundu, D.J., Prakash, A., Frericks-Zipper, A., Eisenacher, M. et al. (2022) The PRIDE database resources in 2022: a hub for mass spectrometry-based proteomics evidences. Nucleic Acids Res, 50, D543–D552.

