## Supplemental Figures and Tables for "Altered mRNA transport and local translation in iNeurons with RNA binding protein knockdown"

### Supplementary figures and legends

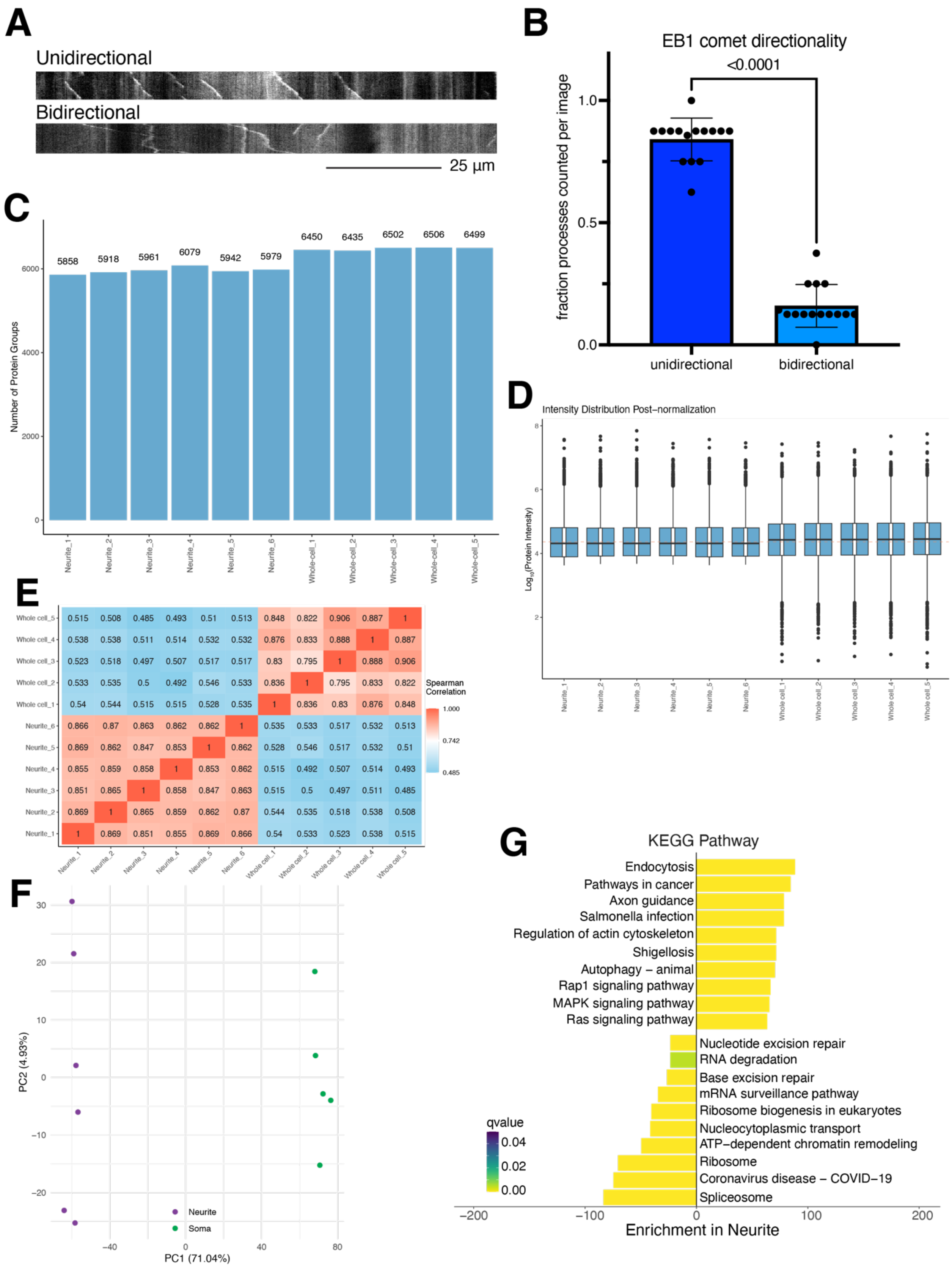

Figure S1: Neurites on underside of membrane are primarily axons. Related to Figure 1.

A. Representative kymographs of EB1 comets from 30 second videos. Processes with unidirectional EB1 comets (top, axons) and bidirectional comets (bottom, dendrites) are shown. Scale bar 25 μm.
B. Quantification of EB1 comet directionality. Neurites from 15 images taken from 2 chambers were quantified as either unidirectional or bidirectional and the fraction of total processes counted per image. 7-8 processes were quantified per image. p-value from two-tailed paired t-test.
C. Number of protein groups identified for each sample; neurite and whole cell samples had a similar number of protein groups.
D. Protein intensity distribution values after normalization for each sample shows similar distributions.
E. Correlation plot of proteomics data shows high correlation between samples from the same cellular fraction.
F. PCA plot of neurite and soma samples shows most of the difference between samples comes from the cellular fraction, not sample-to-sample difference.
G. KEGG analysis of proteins enriched or depleted in neurites shows axon and neurodegeneration-associated terms for proteins enriched in neurites, while proteins enriched in the whole cell fraction are related to nuclear functions (base repair, nucleocytoplasmic transport, splicing).

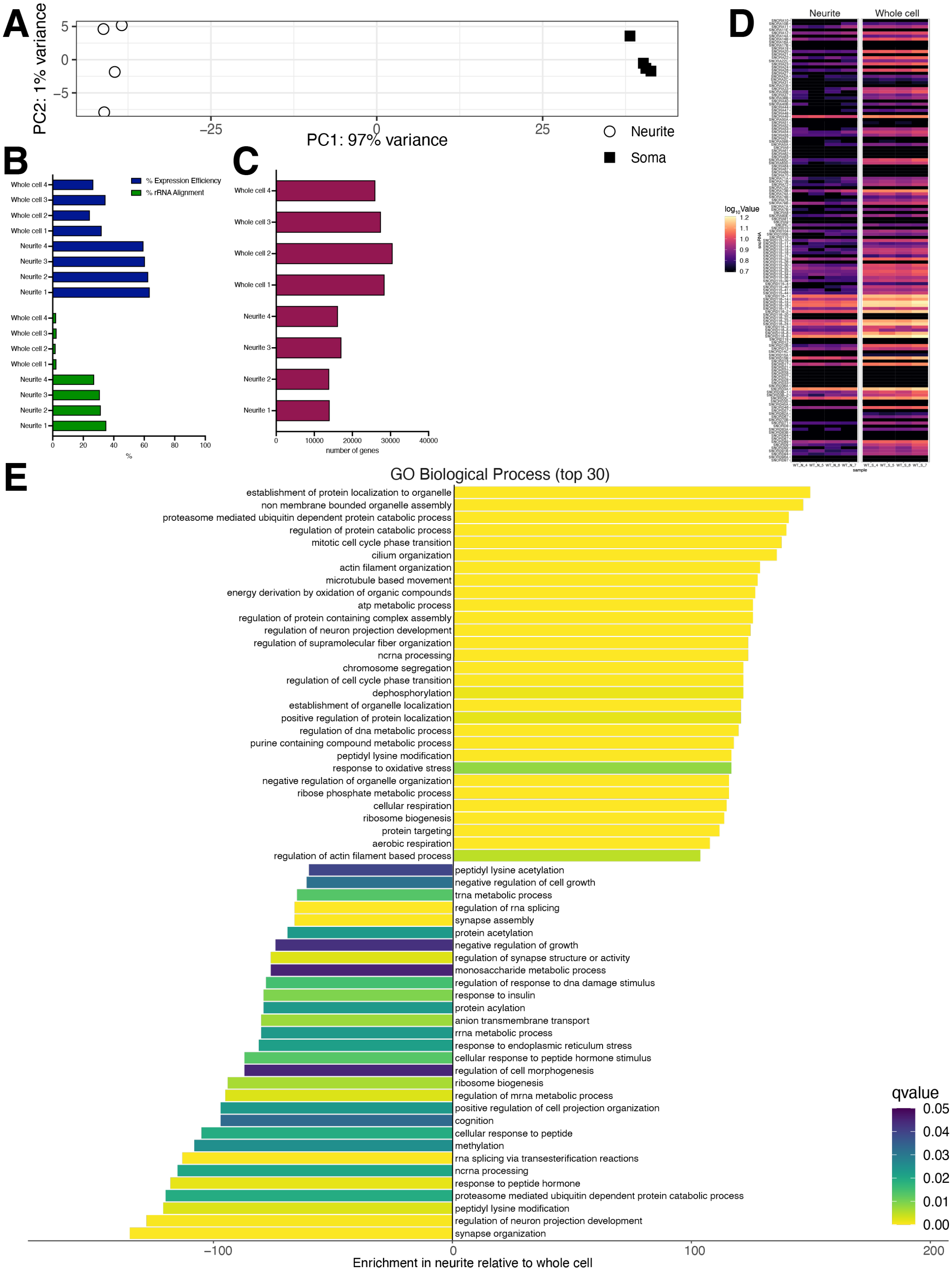
A
PC2: 1% variance
PC1: 97% variance
5
0
−5
−25
0
25
Neurite
Soma
B
Whole cell 4
Whole cell 3
Whole cell 2
Whole cell 1
Neurite 4
Neurite 3
Neurite 2
Neurite 1
% Expression Efficiency
% rRNA Alignment
0
20
40
60
80
100
%
C
number of genes
0
10000
20000
30000
40000
D
Neurite
Whole cell
log10Value
1.2
1.1
1.0
0.9
0.8
0.7
snoRNA
sample
E
GO Biological Process (top 30)
establishment of protein localization to organelle
non membrane bounded organelle assembly
proteasome mediated ubiquitin dependent protein catabolic process
regulation of protein catabolic process
mitotic cell cycle phase transition
cilium organization
actin filament organization
microtubule based movement
energy derivation by oxidation of organic compounds
atp metabolic process
regulation of protein containing complex assembly
regulation of neuron projection development
regulation of supramolecular fiber organization
ncrna processing
chromosome segregation
regulation of cell cycle phase transition
dephosphorylation
establishment of organelle localization
positive regulation of protein localization
regulation of dna metabolic process
purine containing compound metabolic process
peptidyl lysine modification
response to oxidative stress
negative regulation of organelle organization
ribose phosphate metabolic process
cellular respiration
ribosome biogenesis
protein targeting
aerobic respiration
regulation of actin filament based process
peptidyl lysine acetylation
negative regulation of cell growth
trna metabolic process
regulation of rna splicing
synapse assembly
protein acetylation
negative regulation of growth
regulation of synapse structure or activity
monosaccharide metabolic process
regulation of response to dna damage stimulus
response to insulin
protein acylation
anion transmembrane transport
rrna metabolic process
response to endoplasmic reticulum stress
cellular response to peptide hormone stimulus
regulation of cell morphogenesis
ribosome biogenesis
regulation of mrna metabolic process
positive regulation of cell projection organization
cognition
cellular response to peptide
methylation
rna splicing via transesterification reactions
ncrna processing
response to peptide hormone
proteasome mediated ubiquitin dependent protein catabolic process
peptidyl lysine modification
regulation of neuron projection development
synapse organization
qvalue
0.05
0.04
0.03
0.02
0.01
0.00
−100
0
100
200
Enrichment in neurite relative to whole cell

Figure S2: Differences between neurite and whole cell fractions by transcriptomics. Related to Figure 2.

A. Principal component analysis of transcriptomics data shows clear separation of neurite fractions from whole cell fractions along a single axis accounting for 97% of the variance between samples.
B. Percent expression efficiency and percent ribosomal RNA alignment for neurite and whole cell samples. rRNA alignment is higher for neurite samples.
C. Number of genes aligned to for neurite and whole cell samples.
D. Heatmap of snoRNA expression values in each sample for neurite and whole cell samples. As expected, whole cell samples have higher amounts of snoRNAs.
E. GO Biological Process terms enriched in neurite (top, positive values) or whole cell (bottom, negative values) show many terms related to ribosomes, transport, and neurons.

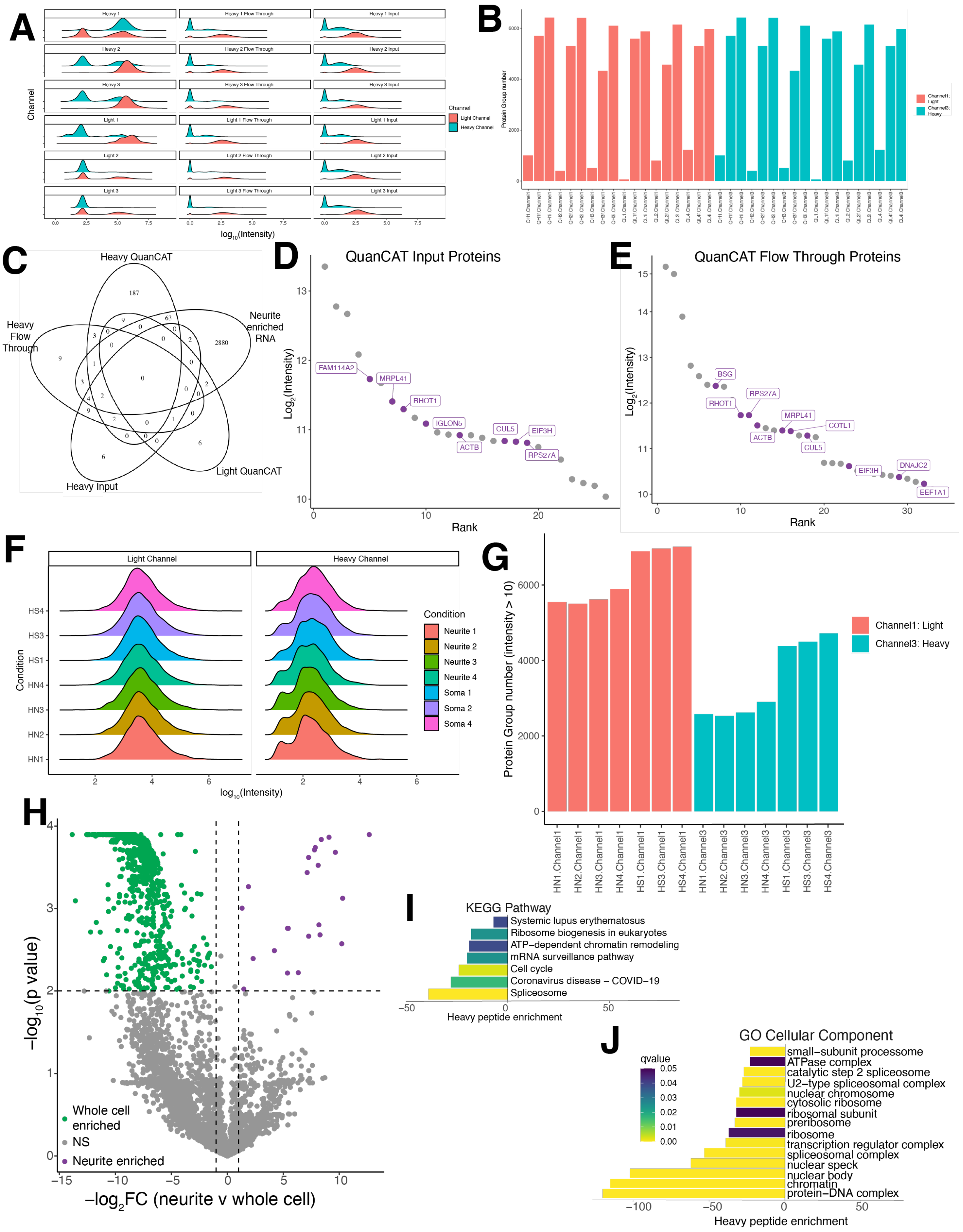

Figure S3: QuanCAT and pSILC quality control. Related to Figure 4.

A. Density plots of heavy and light channels for each QuanCAT sample, including inputs and controls.

B. Number of identified protein groups in both channels for all QuanCAT samples.
C. Venn diagram identifying overlap between the heavy QuanCAT samples, light QuanCAT samples (not exposed to heavy amino acids but treated with AHA), heavy flow through, and heavy input samples, as well as the neurite enriched RNAs. 8 proteins overlapped between the heavy QuanCAT and heavy flow through samples, while 4 overlapped between the input and heavy QuanCAT, demonstrating the specificity of the AHA isolation.
D. Rank plot of heavy proteins identified in the input fractions shows very few proteins, with some overlaps with the transcripts enriched in neurites (purple).
E. Rank plot of heavy proteins identified in the flow through fractions shows very few proteins, with some overlaps with the transcripts enriched in neurites (purple).
F. Density plots of heavy and light channels for each pSILAC sample.
G. Number of identified protein groups in both channels for all pSILAC experiments.
H. Volcano plot of neurite (purple) and soma (green) enriched newly translated proteins. More proteins are translated in the neuritic fraction than the soma fraction. n=4 neurite samples, n=3 whole cell samples.
I. KEGG pathway analysis of proteins containing heavy amino acids in neurites versus whole cell shows many pathways related to nuclear functions, like ribosome biogenesis and chromatin remodeling.
J. Cell component GO analysis of proteins containing heavy amino acids in neurites versus whole cell shows many nucleus-related terms.

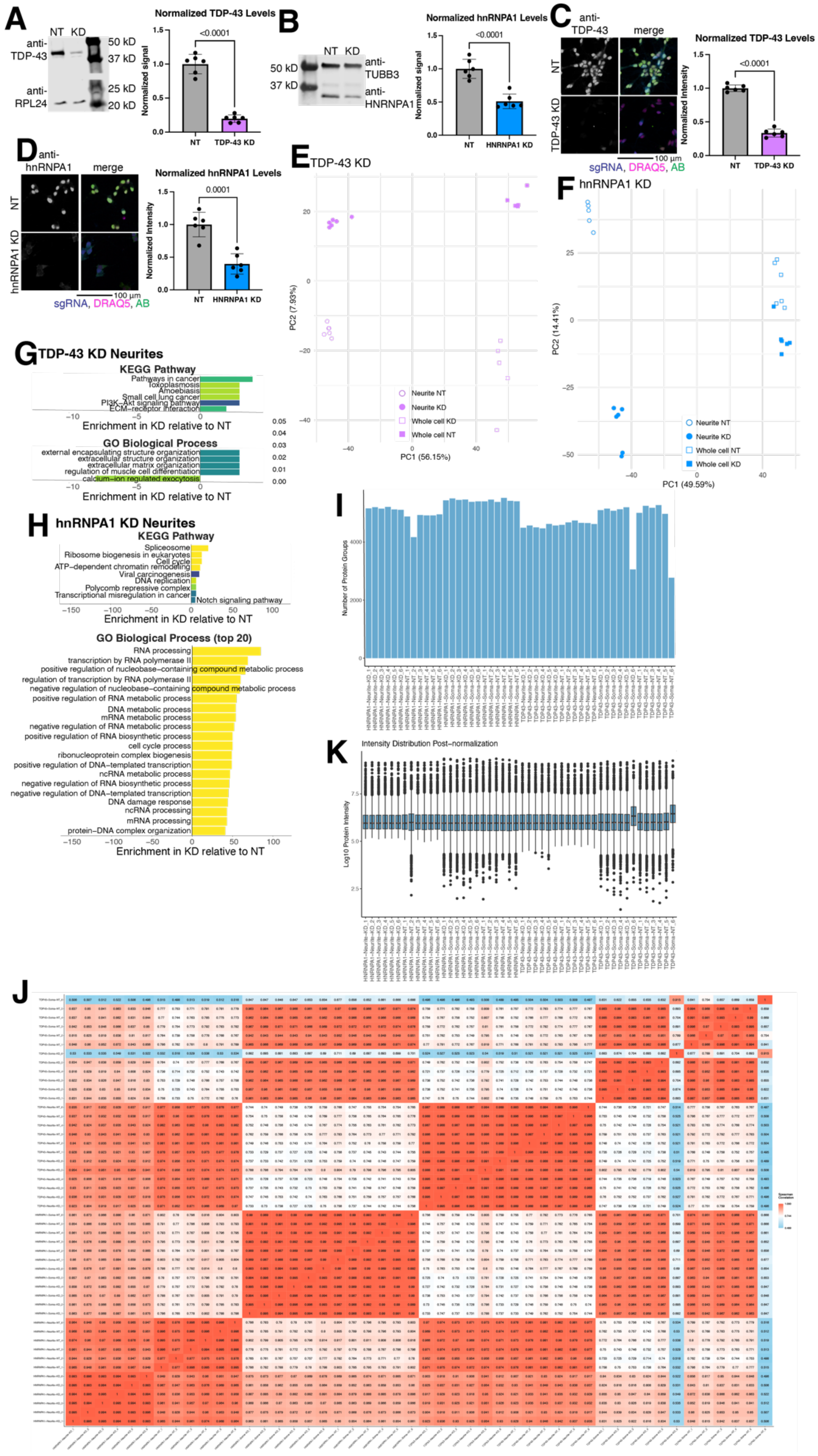

A
NT
KD
anti-TDP-43
anti-RPL24
50 kD
37 kD
25 kD
20 kD
Normalized TDP-43 Levels
<0.0001
Normalized signal
NT
TDP-43 KD
B
NT
KD
50 kD
37 kD
anti-TUBB3
anti-HNRNPA1
Normalized hnRNPA1 Levels
<0.0001
Normalized signal
NT
HNRNPA1 KD
C
anti-TDP-43
merge
NT
TDP-43 KD
100 µm
sgRNA, DRAQ5, AB
Normalized TDP-43 Levels
<0.0001
Normalized Intensity
NT
TDP-43 KD
D
anti-hnRNPA1
merge
NT
hnRNPA1 KD
100 µm
sgRNA, DRAQ5, AB
Normalized hnRNPA1 Levels
0.0001
Normalized Intensity
NT
HNRNPA1 KD
E
TDP-43 KD
PC2 (7.93%)
PC1 (56.15%)
Neurite NT
Neurite KD
Whole cell KD
Whole cell NT
F
hnRNPA1 KD
PC2 (14.41%)
PC1 (49.59%)
Neurite NT
Neurite KD
Whole cell NT
Whole cell KD
G
TDP-43 KD Neurites
KEGG Pathway
Pathways in cancer
Toxoplasmosis
Amoebiasis
Small cell lung cancer
PI3K–Akt signaling pathway
ECM–receptor interaction
Enrichment in KD relative to NT
GO Biological Process
external encapsulating structure organization
extracellular structure organization
extracellular matrix organization
regulation of muscle cell differentiation
calcium–ion regulated exocytosis
Enrichment in KD relative to NT
H
hnRNPA1 KD Neurites
KEGG Pathway
Spliceosome
Ribosome biogenesis in eukaryotes
Cell cycle
ATP–dependent chromatin remodeling
Viral carcinogenesis
DNA replication
Polycomb repressive complex
Transcriptional misregulation in cancer
Notch signaling pathway
Enrichment in KD relative to NT
GO Biological Process (top 20)
RNA processing
transcription by RNA polymerase II
positive regulation of nucleobase–containing compound metabolic process
regulation of transcription by RNA polymerase II
negative regulation of nucleobase–containing compound metabolic process
positive regulation of RNA metabolic process
DNA metabolic process
mRNA metabolic process
negative regulation of RNA metabolic process
positive regulation of RNA biosynthetic process
cell cycle process
ribonucleoprotein complex biogenesis
positive regulation of DNA–templated transcription
ncRNA metabolic process
negative regulation of RNA biosynthetic process
negative regulation of DNA–templated transcription
DNA damage response
ncRNA processing
mRNA processing
protein–DNA complex organization
Enrichment in KD relative to NT
I
Number of Protein Groups
K
Intensity Distribution Post–normalization
Log10 Protein Intensity
J

Figure S4: sgRNAs effectively knock down target proteins. Related to Figure 5.

A-B. Western blot validation of RBP KD for (A) TDP-43 and (B) hnRNPA1. Representative western blot is shown on the left, with quantification to the right. P-values from two-tailed unpaired t-test.

C-D. Immunofluorescence validation of RBP KD for (C) TDP-43 and (D) hnRNPA1. Representative images are shown on the left, with quantification to the right. IF shows similar levels of KD to western blots in A-B. P-values from two-tailed unpaired t-test.

E-F. PCA plots of proteomics experiments for (E) TDP-43 KD and (F) hnRNPA1 KD show the majority of variance (48-60%, PC1) between samples is due to the neurite to whole cell difference rather than KDs (<20%, PC2).

G. KEGG and GO BP analysis comparing TDP-43 KD to NT neurites shows various enriched pathways, including extracellular matrix organization.

H. KEGG and GO BP analysis comparing hnRNPA1 KD to NT neurites shows various enriched pathways, including many RNA and DNA metabolism related terms.

I. Number of non-zero proteins for each sample, used to identify samples with few proteins detected. Two samples, TDP-43 KD soma 6 and TDP-43 NT soma 6, had low protein counts and were excluded from the analysis.

J. Correlation plot of all samples identifies 3 samples (TDP43-Soma-NT_6, TDP43-Soma-KD_6, HNRNPA1-Neurite-NT_2) that are not well correlated with other samples of the same type and thus were excluded from the analysis.

K. Graph of peptide intensity distribution for each sample after normalization. Only 2 samples (TDP-43 KD soma 6 and TDP-43 NT soma 6) have different intensity distributions and were removed from the analysis.

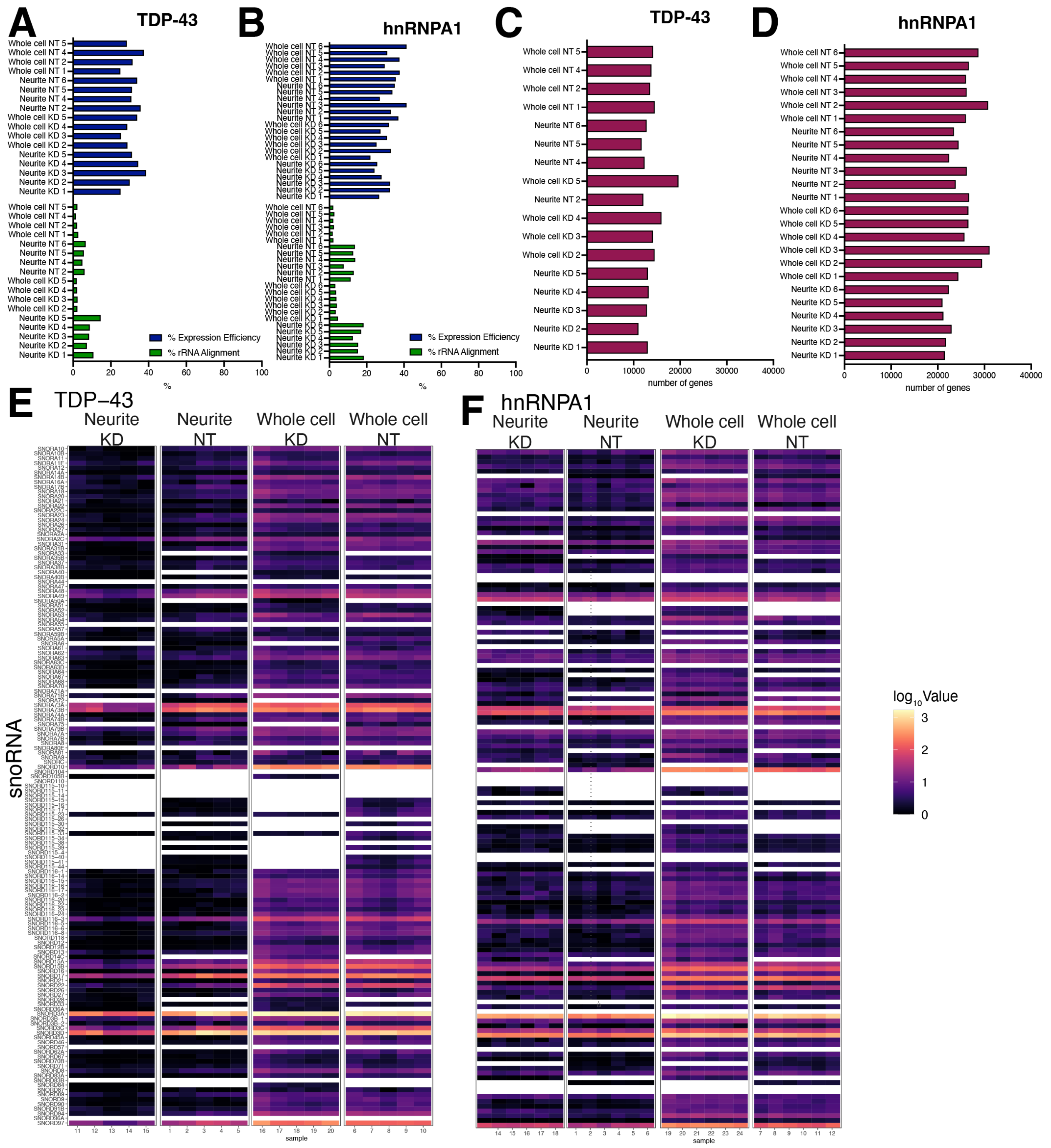

Figure S5: Quality control of RNA sequencing. Related to Figure 6.

A-B: Percent expression efficiency and percent ribosomal RNA alignment for (A) TDP-43 and (B) hnRNPA1 KD samples. rRNA alignment is higher for neurite samples.

C-D: Number of genes aligned to for (C) TDP-43 and (D) hnRNPA1 KD samples. hnRNPA1 samples align to the most genes.

E-F: Heatmap of snoRNA expression values in each sample for (E) TDP-43 and (F) hnRNPA1 KD samples. As expected, whole cell samples have higher amounts of snoRNAs.

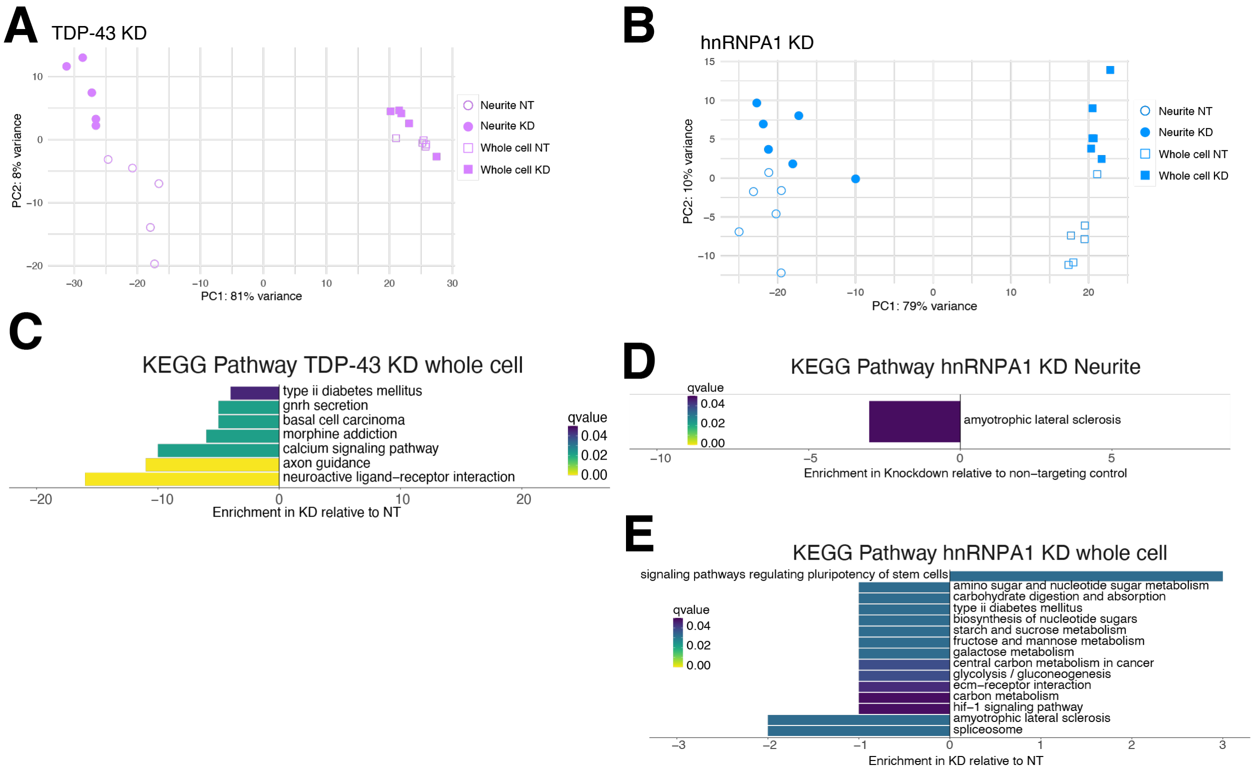

Figure S6: Neurite versus whole cell dominates difference between guides in transcriptomics. Related to Figure 6.

A. PCA plot of TDP-43 KD neurite and whole cell RNA seq shows that most of the difference (81%) between the samples comes from the cell compartment, while a minority (8%) can be explained by the difference between the KD and NT guides.
B. PCA plot of hnRNPA1 KD neurite and whole cell RNA seq shows that most of the difference (79%) between the samples comes from the cell compartment, while a minority (10%) can be explained by the difference between the KD and NT guides.
C. KEGG pathway analysis of transcripts changed in the whole cell in TDP-43 KD versus NT neurons. Axon guidance and neuroactive ligand-receptor interaction are decreased. No KEGG terms were significantly enriched in neurites.
D. KEGG pathway analysis of transcripts changed in hnRNPA1 KD versus NT neurites. The only significant term is a decrease in the ALS term.
E. KEGG pathway analysis of transcripts changed in the whole cell in hnRNPA1 KD versus NT neurons. Sugar metabolism and ALS terms are decreased.

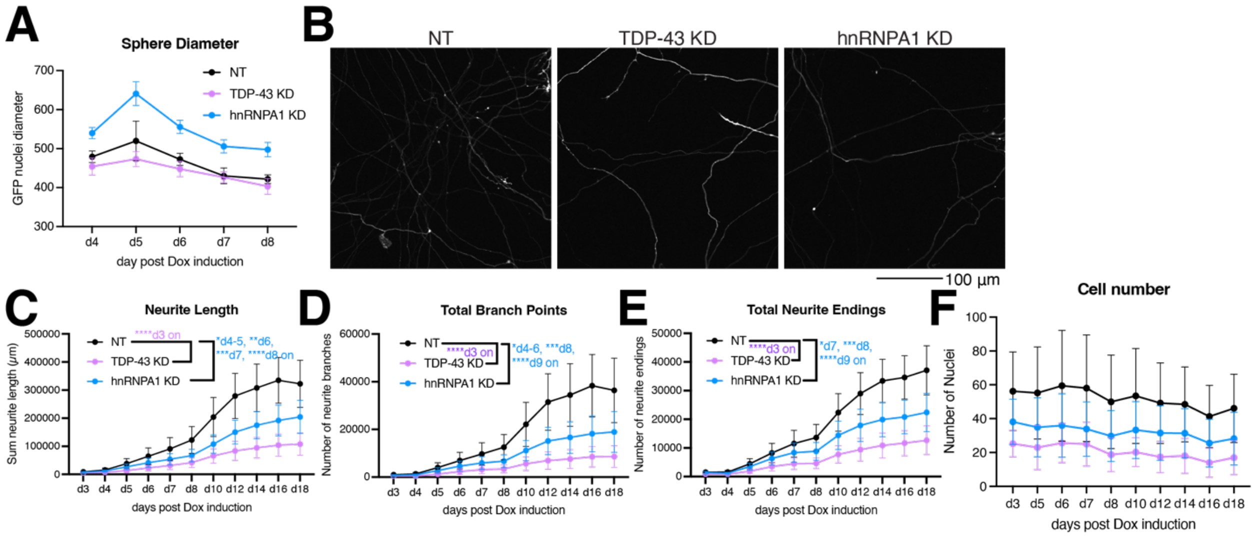

Figure S7: 2D neurite outgrowth shows similar neurite outgrowth defects to neurospheres. Related to Figure 7.

A. Plot of nucleus diameter of spheres in Figure 7D-E. hnRNPA1 KD spheres are slightly larger (have more cells), but NT and TDP-43 KD all have similar sizes. There is no significant change in the area taken up by the cell bodies day to day.
B. Representative images of 2D neurite outgrowth experiment. Image is cytosolic mScarlet channel. n=24 wells per genotype, scale bar 100 µm.
C. C-F: Quantification of 2D neurite outgrowth and neuron survival longitudinal imaging. TDP-43 and hnRNPA1 KD have decreased total neurite length (D), total branch points (E), and number of neurite endings (F). However, there is no significant change in cell counts over the course of the experiment (G). P-values from two-way ANOVA with Dunnett's multiple comparison test.

**Supplemental Tables**

Table S1: Identified cryptic exons in TDP-43 KD neurons.

Table S2: Identified cryptic exons in hnRNPA1 KD neurons.

Table S1: Identified cryptic exons in TDP-43 KD neurons.

| | module_id | gene_id | gene_name | seqid | strand | lsv_id | event_id | complex | module_ever | denovo | junction_nan | junctions_co | event_non_cl | event_chang | junction_cha | refPSI | KDSoma_me | deltaPSI | prob | padjust | event | abs(deltaPSI) |
|---|---|---|---|---|---|---|---|---|---|---|---|---|---|---|---|---|---|---|---|---|---|---|
| 1 | ENSG000001 | ENSG000001 | NUP210 | chr3 | - | | ENSG000001 | TRUE | tandem_cass | TRUE | Distal | chr3:133660 | FALSE | FALSE | TRUE | 0.014 | 0.304 | 0.279 | 1 | 0 | AFE/ALE | 0.279 |
| 2 | ENSG000001 | ENSG000001 | MICAL1 | chr6 | - | | ENSG000001 | TRUE | cassette^6\|t | TRUE | Distal | chr6:109447 | FALSE | FALSE | TRUE | 0.007 | 0.138 | 0.111 | 0.976 | 0.012 | AFE/ALE | 0.111 |
| 3 | ENSG000001 | ENSG000001 | UBC | chr12 | - | | ENSG000001 | TRUE | putative_alt3 | TRUE | Proximal | chr12:12491 | FALSE | FALSE | FALSE | 0.047 | 0.154 | 0.12 | 0.932 | 0.03 | AFE/ALE | 0.12 |
| 4 | ENSG000001 | ENSG000001 | UBC | chr12 | - | | ENSG000001 | TRUE | putative_alt3 | TRUE | Distal | chr12:12491 | FALSE | FALSE | TRUE | 0.003 | 0.182 | 0.157 | 1 | 0 | AFE/ALE | 0.157 |
| 5 | ENSG00000 | ENSG00000 | KLHL13 | chrX | - | ENSG00000 | ENSG00000 | TRUE | cassette^1\|a | TRUE | Distal | chrX:117914 | FALSE | TRUE | TRUE | 0.004 | 0.221 | 0.198 | 0.997 | 0.001 | AFE/ALE | 0.198 |
| 6 | ENSG00000 | ENSG00000 | SPPL2B | chr19 | + | ENSG00000 | ENSG00000 | TRUE | cassette^1\|i | FALSE | C2_A | chr19:23373 | FALSE | FALSE | TRUE | 0.041 | 0.244 | 0.205 | 1 | 0 | cassette | 0.205 |
| 7 | ENSG00000 | ENSG00000 | TSPOAP1 | chr17 | - | ENSG00000 | ENSG00000 | TRUE | alt5^1\|ir^2 | TRUE | Proximal | chr17:58305 | FALSE | FALSE | TRUE | 0.01 | 0.16 | 0.135 | 0.995 | 0.002 | AFE/ALE | 0.135 |
| 8 | ENSG00000 | ENSG00000 | ITGA3 | chr17 | + | ENSG00000 | ENSG00000 | FALSE | cassette^1 | FALSE | C1_A | chr17:50080 | FALSE | FALSE | TRUE | 0.043 | 0.301 | 0.258 | 1 | 0 | cassette | 0.258 |
| 9 | ENSG00000 | ENSG00000 | IFFO1 | chr12 | - | ENSG00000 | ENSG00000 | FALSE | cassette^1 | FALSE | C1_A | chr12:65494 | FALSE | FALSE | FALSE | 0.037 | 0.165 | 0.128 | 0.91 | 0.028 | cassette | 0.128 |
| 10 | ENSG00000 | ENSG00000 | SYT7 | chr11 | - | ENSG00000 | ENSG00000 | TRUE | cassette^4\|t | TRUE | C1_A | chr11:61547 | FALSE | TRUE | TRUE | 0 | 0.552 | 0.539 | 1 | 0 | cassette | 0.539 |
| 11 | ENSG00000 | ENSG00000 | MNAT1 | chr14 | + | ENSG00000 | ENSG00000 | TRUE | cassette^1\|i | TRUE | C1_A | chr14:60812 | FALSE | TRUE | TRUE | 0.047 | 0.673 | 0.633 | 1 | 0 | cassette | 0.633 |
| 12 | ENSG00000 | ENSG00000 | GRAMD1B | chr11 | + | ENSG00000 | ENSG00000 | TRUE | cassette^2\|r | FALSE | C1_A | chr11:12357 | FALSE | FALSE | TRUE | 0.031 | 0.149 | 0.119 | 0.961 | 0.018 | cassette | 0.119 |
| 13 | ENSG00000 | ENSG00000 | POU2F2 | chr19 | - | ENSG00000 | ENSG00000 | FALSE | putative_afe | TRUE | Proximal | chr19:42122 | FALSE | TRUE | TRUE | 0.002 | 0.344 | 0.32 | 1 | 0 | AFE/ALE | 0.32 |
| 14 | ENSG00000 | ENSG00000 | BRD9 | chr5 | - | ENSG00000 | ENSG00000 | TRUE | cassette^1\|i | FALSE | C1_C2 | chr5:879889 | FALSE | FALSE | FALSE | 0.002 | 0.143 | -0.113 | 0.938 | 0.029 | skipping | 0.113 |
| 15 | ENSG00000 | ENSG00000 | SYNE2 | chr14 | + | ENSG00000 | ENSG00000 | FALSE | putative_afe | TRUE | Proximal | chr14:63853 | FALSE | FALSE | FALSE | 0.039 | 0.164 | 0.123 | 0.932 | 0.027 | AFE/ALE | 0.123 |
| 16 | ENSG00000 | ENSG00000 | USP36 | chr17 | - | ENSG00000 | ENSG00000 | TRUE | ir^2\|putative | TRUE | Proximal | chr17:78836 | FALSE | FALSE | TRUE | 0.019 | 0.195 | 0.163 | 0.996 | 0.001 | AFE/ALE | 0.163 |
| 17 | ENSG00000 | ENSG00000 | USP13 | chr3 | + | ENSG00000 | ENSG00000 | FALSE | alt5^1 | FALSE | Proximal | chr3:179701 | FALSE | TRUE | TRUE | 0.05 | 0.651 | 0.62 | 1 | 0 | AFE/ALE | 0.62 |
| 18 | ENSG00000 | ENSG00000 | CAMK2B | chr7 | - | ENSG00000 | ENSG00000 | TRUE | cassette^4\|t | TRUE | C1_A | chr7:442585 | FALSE | FALSE | TRUE | 0.001 | 0.311 | 0.289 | 1 | 0 | cassette | 0.289 |
| 19 | ENSG00000 | ENSG00000 | CAMK2B | chr7 | - | ENSG00000 | ENSG00000 | TRUE | cassette^4\|t | TRUE | C1_A | chr7:442586 | FALSE | FALSE | TRUE | 0.002 | 0.227 | 0.204 | 1 | 0 | cassette | 0.204 |
| 20 | ENSG00000 | ENSG00000 | CAMK2B | chr7 | - | ENSG00000 | ENSG00000 | TRUE | ir^3\|ale^1\|a | TRUE | Proximal | chr7:442209 | FALSE | TRUE | TRUE | 0 | 0.395 | 0.378 | 1 | 0 | AFE/ALE | 0.378 |
| 21 | ENSG00000 | ENSG00000 | CAMK2B | chr7 | - | ENSG00000 | ENSG00000 | TRUE | cassette^4\|t | TRUE | Distal | chr7:442546 | FALSE | FALSE | TRUE | 0.004 | 0.5 | 0.48 | 1 | 0 | AFE/ALE | 0.48 |
| 22 | ENSG00000 | ENSG00000 | PSD | chr10 | - | ENSG00000 | ENSG00000 | TRUE | cassette^3\|t | TRUE | C1_A | chr10:10241 | FALSE | FALSE | TRUE | 0 | 0.408 | 0.394 | 1 | 0 | cassette | 0.394 |
| 23 | ENSG00000 | ENSG00000 | PSD | chr10 | - | ENSG00000 | ENSG00000 | TRUE | cassette^3\|t | TRUE | C2_A_Last | chr10:10241 | FALSE | FALSE | TRUE | 0 | 0.297 | 0.276 | 1 | 0 | cassette | 0.276 |
| 24 | ENSG00000 | ENSG00000 | CDON | chr11 | - | ENSG00000 | ENSG00000 | FALSE | alt3^1 | TRUE | Distal | chr11:12598 | FALSE | TRUE | TRUE | 0.002 | 0.332 | 0.314 | 1 | 0 | AFE/ALE | 0.314 |
| 25 | ENSG00000 | ENSG00000 | ZFAT | chr8 | - | ENSG00000 | ENSG00000 | FALSE | putative_afe | TRUE | Proximal | chr8:134637 | FALSE | TRUE | TRUE | 0.006 | 0.282 | 0.243 | 1 | 0 | AFE/ALE | 0.243 |
| 26 | ENSG00000 | ENSG00000 | PFKP | chr10 | + | ENSG00000 | ENSG00000 | TRUE | cassette^2\|a | TRUE | C1_A | chr10:30993 | FALSE | TRUE | TRUE | 0.012 | 0.689 | 0.685 | 1 | 0 | cassette | 0.685 |
| 27 | ENSG00000 | ENSG00000 | PFKP | chr10 | + | ENSG00000 | ENSG00000 | TRUE | cassette^2\|a | TRUE | C1_A | chr10:30993 | FALSE | FALSE | FALSE | 0.001 | 0.142 | 0.112 | 0.94 | 0.029 | cassette | 0.112 |
| 28 | ENSG00000 | ENSG00000 | ADGRL1 | chr19 | - | ENSG00000 | ENSG00000 | TRUE | cassette^1\|i | TRUE | C1_A | chr19:14176 | FALSE | TRUE | TRUE | 0 | 0.256 | 0.234 | 1 | 0 | cassette | 0.234 |
| 29 | ENSG00000 | ENSG00000 | ADGRL1 | chr19 | - | ENSG00000 | ENSG00000 | TRUE | cassette^1\|i | TRUE | Distal | chr19:14170 | FALSE | FALSE | TRUE | 0 | 0.217 | 0.198 | 1 | 0 | AFE/ALE | 0.198 |
| 30 | ENSG00000 | ENSG00000 | SPEG | chr2 | + | ENSG00000 | ENSG00000 | TRUE | ir^3\|ale^2\|p | TRUE | Proximal | chr2:219465 | FALSE | TRUE | TRUE | 0.014 | 0.326 | 0.309 | 1 | 0 | AFE/ALE | 0.309 |
| 31 | ENSG00000 | ENSG00000 | KCNQ2 | chr20 | - | ENSG00000 | ENSG00000 | TRUE | cassette^1\|i | TRUE | C1_C2 | chr20:63439 | FALSE | TRUE | TRUE | 0.028 | 0.261 | -0.242 | 1 | 0 | skipping | 0.242 |
| 32 | ENSG00000 | ENSG00000 | KCNQ2 | chr20 | - | ENSG00000 | ENSG00000 | TRUE | cassette^1\|i | TRUE | Distal | chr20:63439 | FALSE | TRUE | TRUE | 0.039 | 0.256 | 0.229 | 1 | 0 | AFE/ALE | 0.229 |
| 33 | ENSG00000 | ENSG00000 | CTTNBP2 | chr7 | - | ENSG00000 | ENSG00000 | TRUE | cassette^1\|i | TRUE | Proximal | chr7:117861 | FALSE | FALSE | TRUE | 0.013 | 0.168 | 0.145 | 0.999 | 0 | AFE/ALE | 0.145 |
| 34 | ENSG00000 | ENSG00000 | ACTL6B | chr7 | - | ENSG00000 | ENSG00000 | TRUE | cassette^1\|i | TRUE | C1_A | chr7:100650 | FALSE | TRUE | TRUE | 0.021 | 0.636 | 0.626 | 1 | 0 | cassette | 0.626 |
| 35 | ENSG00000 | ENSG00000 | PIAS2 | chr18 | - | ENSG00000 | ENSG00000 | FALSE | cassette^1 | FALSE | C2_C1 | chr18:46855 | FALSE | FALSE | FALSE | 0.033 | 0.159 | -0.125 | 0.911 | 0.03 | skipping | 0.125 |
| 36 | ENSG00000 | ENSG00000 | UIMC1 | chr5 | - | ENSG00000 | ENSG00000 | FALSE | cassette^1 | TRUE | C1_A | chr5:176980 | FALSE | FALSE | TRUE | 0.021 | 0.123 | 0.1 | 0.957 | 0.022 | cassette | 0.1 |
| 37 | ENSG00000 | ENSG00000 | PHACTR3 | chr20 | + | ENSG00000 | ENSG00000 | TRUE | putative_afe | TRUE | Proximal | chr20:59742 | FALSE | FALSE | FALSE | 0.001 | 0.131 | 0.103 | 0.939 | 0.031 | AFE/ALE | 0.103 |
| 38 | ENSG00000 | ENSG00000 | CRLS1 | chr20 | + | ENSG00000 | ENSG00000 | TRUE | cassette^2\|i | TRUE | C1_A | chr20:60154 | FALSE | TRUE | TRUE | 0.002 | 0.229 | 0.202 | 1 | 0 | cassette | 0.202 |
| 39 | ENSG00000 | ENSG00000 | CRLS1 | chr20 | + | ENSG00000 | ENSG00000 | TRUE | cassette^2\|i | TRUE | Distal | chr20:60262 | FALSE | FALSE | TRUE | 0.008 | 0.213 | 0.187 | 1 | 0 | AFE/ALE | 0.187 |
| 40 | ENSG00000 | ENSG00000 | DNAAF9 | chr20 | - | ENSG00000 | ENSG00000 | TRUE | cassette^1\|i | TRUE | C1_A | chr20:33446 | FALSE | TRUE | TRUE | 0.002 | 0.441 | 0.426 | 1 | 0 | cassette | 0.426 |
| 41 | ENSG00000 | ENSG00000 | GRAMD1A | chr19 | + | ENSG00000 | ENSG00000 | TRUE | cassette^1\|i | TRUE | C1_A | chr19:35000 | FALSE | TRUE | TRUE | 0.006 | 0.185 | 0.161 | 1 | 0 | cassette | 0.161 |
| 42 | ENSG00000 | ENSG00000 | GRAMD1A | chr19 | + | ENSG00000 | ENSG00000 | TRUE | cassette^1\|i | FALSE | Distal | chr19:35001 | FALSE | FALSE | TRUE | 0.025 | 0.314 | 0.293 | 1 | 0 | AFE/ALE | 0.293 |
| 43 | ENSG00000 | ENSG00000 | AARS1 | chr16 | - | ENSG00000 | ENSG00000 | TRUE | cassette^1\|i | TRUE | C1_A | chr16:70272 | FALSE | TRUE | TRUE | 0 | 0.286 | 0.27 | 1 | 0 | cassette | 0.27 |
| 44 | ENSG00000 | ENSG00000 | G2E3 | chr14 | + | ENSG00000 | ENSG00000 | TRUE | cassette^1\|i | TRUE | Proximal | chr14:30559 | FALSE | FALSE | TRUE | 0.001 | 0.136 | 0.11 | 0.956 | 0.021 | AFE/ALE | 0.11 |
| 45 | ENSG00000 | ENSG00000 | NUP188 | chr9 | + | ENSG00000 | ENSG00000 | FALSE | cassette^1 | TRUE | C1_A | chr9:128952 | FALSE | TRUE | TRUE | 0.003 | 0.783 | 0.781 | 1 | 0 | cassette | 0.781 |
| 46 | ENSG00000 | ENSG00000 | RANBP1 | chr22 | + | ENSG00000 | ENSG00000 | FALSE | cassette^1 | FALSE | C2_A | chr22:20122 | FALSE | FALSE | TRUE | 0.003 | 0.123 | 0.102 | 0.981 | 0.01 | cassette | 0.102 |
| 47 | ENSG00000 | ENSG00000 | BCL2L13 | chr22 | + | ENSG00000 | ENSG00000 | TRUE | cassette^7\|t | TRUE | J3 | chr22:17702 | FALSE | FALSE | TRUE | 0.006 | 0.134 | 0.107 | 0.963 | 0.019 | cassette | 0.107 |
| 48 | ENSG000001 | ENSG000001 | SNRPD3 | chr22 | + | ENSG000001 | ENSG000001 | TRUE | ir^2\|putative | TRUE | Proximal | chr22:24572 | FALSE | FALSE | TRUE | 0.004 | 0.188 | 0.162 | 0.993 | 0.002 | AFE/ALE | 0.162 |
| 49 | ENSG000001 | ENSG000001 | HDAC10 | chr22 | - | ENSG000001 | ENSG000001 | TRUE | cassette^1\|t | TRUE | C2_A_Last | chr22:50248 | FALSE | FALSE | TRUE | 0.043 | 0.224 | 0.174 | 0.991 | 0.002 | cassette | 0.174 |
| 50 | ENSG000001 | ENSG000001 | GSS | chr20 | - | ENSG000001 | ENSG000001 | FALSE | ale^1 | FALSE | Proximal | chr20:34950 | FALSE | TRUE | TRUE | 0.001 | 0.231 | 0.202 | 1 | 0 | AFE/ALE | 0.202 |
| 51 | ENSG000001 | ENSG000001 | PRELID3B | chr20 | - | ENSG000001 | ENSG000001 | TRUE | cassette^2\|a | FALSE | C1_C2 | chr20:59036 | FALSE | FALSE | TRUE | 0.001 | 0.157 | -0.132 | 0.998 | 0.001 | skipping | 0.132 |
| 52 | ENSG000001 | ENSG000001 | ATP11C | chrX | - | ENSG000001 | ENSG000001 | TRUE | cassette^5\|p | FALSE | C1_A | chrX:139737 | FALSE | FALSE | TRUE | 0.041 | 0.291 | 0.241 | 0.997 | 0 | cassette | 0.241 |
| 53 | ENSG000001 | ENSG000001 | EEA1 | chr12 | - | ENSG000001 | ENSG000001 | FALSE | cassette^1 | TRUE | C1_A | chr12:92846 | FALSE | TRUE | TRUE | 0.014 | 0.255 | 0.231 | 1 | 0 | cassette | 0.231 |
| 54 | ENSG000001 | ENSG000001 | ZNF423 | chr16 | - | ENSG000001 | ENSG000001 | FALSE | cassette^1 | TRUE | C1_A | chr16:49640 | FALSE | TRUE | TRUE | 0.001 | 0.41 | 0.391 | 1 | 0 | cassette | 0.391 |
| 55 | ENSG000001 | ENSG000001 | NECAB2 | chr16 | + | ENSG000001 | ENSG000001 | TRUE | cassette^1\|i | TRUE | C1_A | chr16:83990 | FALSE | TRUE | TRUE | 0.001 | 0.409 | 0.389 | 1 | 0 | cassette | 0.389 |
| 56 | ENSG000001 | ENSG000001 | USP10 | chr16 | + | ENSG000001 | ENSG000001 | FALSE | cassette^1 | TRUE | C2_A | chr16:84735 | FALSE | FALSE | TRUE | 0.001 | 0.192 | 0.169 | 1 | 0 | cassette | 0.169 |
| 57 | ENSG000001 | ENSG000001 | CORO7-PAM | chr16 | - | ENSG000001 | ENSG000001 | FALSE | putative_ale | TRUE | Proximal | chr16:43674 | FALSE | TRUE | TRUE | 0.019 | 0.347 | 0.323 | 1 | 0 | AFE/ALE | 0.323 |
| 58 | ENSG000001 | ENSG000001 | STMN2 | chr8 | + | ENSG000001 | ENSG000001 | TRUE | ir^2\|ale^2\|p | TRUE | Distal | chr8:796112 | FALSE | FALSE | TRUE | 0.003 | 0.637 | 0.624 | 1 | 0 | AFE/ALE | 0.624 |
| 59 | ENSG000001 | ENSG000001 | SLC1A6 | chr19 | - | ENSG000001 | ENSG000001 | TRUE | cassette^2\|a | TRUE | C1_A | chr19:14979 | FALSE | FALSE | TRUE | 0 | 0.139 | 0.113 | 0.985 | 0.007 | cassette | 0.113 |
| 60 | ENSG000001 | ENSG000001 | GPI | chr19 | + | ENSG000001 | ENSG000001 | TRUE | alt3^1\|alt3_ | TRUE | C2_A2 | chr19:34392 | FALSE | FALSE | TRUE | 0 | 0.127 | 0.101 | 0.965 | 0.018 | cassette | 0.101 |
| 61 | ENSG000001 | ENSG000001 | CCDC9 | chr19 | + | ENSG000001 | ENSG000001 | FALSE | alt5^1 | TRUE | Proximal | chr19:47271 | FALSE | FALSE | TRUE | 0.004 | 0.187 | 0.16 | 0.994 | 0.002 | AFE/ALE | 0.16 |
| 62 | ENSG000001 | ENSG000001 | DENND3 | chr8 | + | ENSG000001 | ENSG000001 | FALSE | putative_alt5 | FALSE | Proximal | chr8:141173 | FALSE | FALSE | FALSE | 0.023 | 0.176 | 0.132 | 0.923 | 0.02 | AFE/ALE | 0.132 |
| 63 | ENSG000001 | ENSG000001 | MAST1 | chr19 | + | ENSG000001 | ENSG000001 | TRUE | cassette^2\|a | TRUE | C1_A | chr19:12841 | FALSE | FALSE | TRUE | 0 | 0.275 | 0.254 | 1 | 0 | cassette | 0.254 |
| 64 | ENSG000001 | ENSG000001 | MAST1 | chr19 | + | ENSG000001 | ENSG000001 | TRUE | cassette^2\|a | TRUE | Distal | chr19:12842 | FALSE | FALSE | TRUE | 0 | 0.151 | 0.124 | 1 | 0 | AFE/ALE | 0.124 |
| 65 | ENSG000001 | ENSG000001 | SIPA1L3 | chr19 | + | ENSG000001 | ENSG000001 | TRUE | ale^2\|putati | TRUE | Distal | chr19:37907 | FALSE | FALSE | TRUE | 0.003 | 0.377 | 0.353 | 1 | 0 | AFE/ALE | 0.353 |
| 66 | ENSG000001 | ENSG000001 | RUNDC3B | chr7 | + | ENSG000001 | ENSG000001 | TRUE | cassette^3\|a | TRUE | Distal | chr7:876285 | FALSE | TRUE | TRUE | 0.001 | 0.261 | 0.238 | 1 | 0 | AFE/ALE | 0.238 |
| 67 | ENSG000001 | ENSG000001 | IQCE | chr7 | + | ENSG000001 | ENSG000001 | TRUE | cassette^3\|i | TRUE | C1_A | chr7:255921 | FALSE | TRUE | TRUE | 0.001 | 0.621 | 0.608 | 1 | 0 | cassette | 0.608 |
| 68 | ENSG000001 | ENSG000001 | IQCE | chr7 | + | ENSG000001 | ENSG000001 | TRUE | cassette^3\|i | TRUE | Proximal | chr7:255921 | FALSE | TRUE | TRUE | 0.013 | 0.202 | 0.181 | 1 | 0 | AFE/ALE | 0.181 |
| 69 | ENSG000001 | ENSG000001 | WASL | chr7 | - | ENSG000001 | ENSG000001 | TRUE | cassette^2\|a | TRUE | C1_C2 | chr7:123684 | FALSE | TRUE | TRUE | 0.001 | 0.554 | -0.547 | 1 | 0 | skipping | 0.547 |
| 70 | ENSG000001 | ENSG000001 | WASL | chr7 | - | ENSG000001 | ENSG000001 | TRUE | cassette^2\|a | TRUE | C2_A | chr7:123684 | FALSE | FALSE | FALSE | 0 | 0.119 | 0.096 | 0.922 | 0.042 | cassette | 0.096 |
| 71 | ENSG000001 | ENSG000001 | DNM1 | chr9 | + | ENSG000001 | ENSG000001 | TRUE | alt3^1\|ir^2 | TRUE | Proximal | chr9:128250 | FALSE | TRUE | TRUE | 0.001 | 0.204 | 0.181 | 1 | 0 | AFE/ALE | 0.181 |
| 72 | ENSG000001 | ENSG000001 | NSMCE4A | chr10 | - | ENSG000001 | ENSG000001 | TRUE | cassette^1\|i | TRUE | C1_A | chr10:12197 | FALSE | FALSE | TRUE | 0.019 | 0.261 | 0.232 | 1 | 0 | cassette | 0.232 |
| 73 | ENSG000001 | ENSG000001 | SPOCK2 | chr10 | - | ENSG000001 | ENSG000001 | TRUE | alt3_5^1\|ir^ | TRUE | E1_E2_J2 | chr10:72088 | FALSE | FALSE | FALSE | 0.01 | 0.131 | 0.103 | 0.931 | 0.034 | cassette | 0.103 |
| 74 | ENSG000001 | ENSG000001 | DNAJC12 | chr10 | - | ENSG000001 | ENSG000001 | TRUE | cassette^1\|p | TRUE | C2_A | chr10:67823 | FALSE | FALSE | FALSE | 0.036 | 0.153 | 0.116 | 0.94 | 0.027 | cassette | 0.116 |
| 75 | ENSG000001 | ENSG000001 | MPP2 | chr17 | - | ENSG000001 | ENSG000001 | TRUE | cassette^1\|a | FALSE | C1_A | chr17:43884 | FALSE | FALSE | FALSE | 0.026 | 0.117 | 0.094 | 0.917 | 0.043 | cassette | 0.094 |

| | | | | | | | | | | | | | | | | | | | | | | |
|---|---|---|---|---|---|---|---|---|---|---|---|---|---|---|---|---|---|---|---|---|---|---|
| 76 | ENSG000001 | ENSG000001 | MADD | chr11 | + | ENSG000001 | ENSG000001 | TRUE | cassette^4\| | TRUE | C1_A | chr11:47325 | FALSE | FALSE | TRUE | 0.005 | 0.195 | 0.17 | 0.999 | 0 | cassette | 0.17 |
| 77 | ENSG000001 | ENSG000001 | MADD | chr11 | + | ENSG000001 | ENSG000001 | TRUE | cassette^4\| | FALSE | C1_A | chr11:47324 | FALSE | FALSE | TRUE | 0.003 | 0.326 | 0.3 | 1 | 0 | cassette | 0.3 |
| 78 | ENSG000001 | ENSG000001 | MADD | chr11 | + | ENSG000001 | ENSG000001 | TRUE | cassette^4\| | TRUE | Proximal | chr11:47325 | FALSE | TRUE | TRUE | 0.009 | 0.248 | 0.222 | 1 | 0 | AFE/ALE | 0.222 |
| 79 | ENSG000001 | ENSG000001 | MADD | chr11 | + | ENSG000001 | ENSG000001 | TRUE | cassette^4\| | TRUE | Proximal | chr11:47325 | FALSE | TRUE | TRUE | 0.003 | 0.281 | 0.259 | 1 | 0 | AFE/ALE | 0.259 |
| 80 | ENSG000001 | ENSG000001 | RNGTT | chr6 | - | ENSG000001 | ENSG000001 | FALSE | alt5^1 | TRUE | Proximal | chr6:889049 | FALSE | FALSE | TRUE | 0.039 | 0.179 | 0.141 | 0.97 | 0.011 | AFE/ALE | 0.141 |
| 81 | ENSG000001 | ENSG000001 | FRMD4B | chr3 | - | ENSG000001 | ENSG000001 | TRUE | ir^1\|ale^1\|a | TRUE | Proximal | chr3:693024 | FALSE | FALSE | TRUE | 0.009 | 0.221 | 0.186 | 0.993 | 0.001 | AFE/ALE | 0.186 |
| 82 | ENSG000001 | ENSG000001 | CCDC88A | chr2 | - | ENSG000001 | ENSG000001 | TRUE | cassette^3\| | FALSE | A_C2 | chr2:553006 | FALSE | FALSE | TRUE | 0.003 | 0.29 | 0.265 | 1 | 0 | cassette | 0.265 |
| 83 | ENSG000001 | ENSG000001 | EHBP1 | chr2 | + | ENSG000001 | ENSG000001 | FALSE | cassette^1 | FALSE | C1_C2 | chr2:628311 | FALSE | FALSE | TRUE | 0.038 | 0.209 | -0.171 | 0.997 | 0.001 | skipping | 0.171 |
| 84 | ENSG000001 | ENSG000001 | ELAPOR1 | chr1 | + | ENSG000001 | ENSG000001 | TRUE | cassette^1\|i | TRUE | C1_C2 | chr1:109197 | FALSE | TRUE | TRUE | 0.018 | 0.241 | -0.216 | 1 | 0 | skipping | 0.216 |
| 85 | ENSG000001 | ENSG000001 | DLGAP3 | chr1 | - | ENSG000001 | ENSG000001 | TRUE | cassette^1\| | TRUE | C1_A | chr1:349290 | FALSE | FALSE | TRUE | 0.001 | 0.232 | 0.206 | 1 | 0 | cassette | 0.206 |
| 86 | ENSG000001 | ENSG000001 | AKT3 | chr1 | - | ENSG000001 | ENSG000001 | FALSE | tandem_cass | TRUE | C1_A | chr1:243527 | FALSE | TRUE | TRUE | 0.002 | 0.228 | 0.203 | 1 | 0 | cassette | 0.203 |
| 87 | ENSG000001 | ENSG000001 | IGSF21 | chr1 | + | ENSG000001 | ENSG000001 | FALSE | cassette^1 | TRUE | C1_A | chr1:182280 | FALSE | FALSE | TRUE | 0.003 | 0.494 | 0.481 | 1 | 0 | cassette | 0.481 |
| 88 | ENSG000001 | ENSG000001 | ERI3 | chr1 | - | ENSG000001 | ENSG000001 | TRUE | cassette^2\| | FALSE | C1_A | chr1:442420 | FALSE | FALSE | TRUE | 0.011 | 0.171 | 0.149 | 1 | 0 | cassette | 0.149 |
| 89 | ENSG000001 | ENSG000001 | PROSER1 | chr13 | - | ENSG000001 | ENSG000001 | TRUE | cassette^2\| | TRUE | C1_A | chr13:39018 | FALSE | FALSE | TRUE | 0.001 | 0.14 | 0.118 | 0.988 | 0.005 | cassette | 0.118 |
| 90 | ENSG000001 | ENSG000001 | NFYB | chr12 | - | ENSG000001 | ENSG000001 | TRUE | cassette^1\| | TRUE | Proximal | chr12:10411 | FALSE | FALSE | TRUE | 0.001 | 0.145 | 0.119 | 0.968 | 0.015 | AFE/ALE | 0.119 |
| 91 | ENSG000001 | ENSG000001 | WASHC3 | chr12 | - | ENSG000001 | ENSG000001 | TRUE | cassette^2\| | FALSE | C2_A | chr12:10204 | FALSE | FALSE | FALSE | 0.018 | 0.125 | 0.1 | 0.921 | 0.04 | cassette | 0.1 |
| 92 | ENSG000001 | ENSG000001 | APAF1 | chr12 | + | ENSG000001 | ENSG000001 | TRUE | alt5^3 | TRUE | Proximal | chr12:98649 | FALSE | FALSE | TRUE | 0.021 | 0.179 | 0.145 | 0.97 | 0.01 | AFE/ALE | 0.145 |
| 93 | ENSG000001 | ENSG000001 | ADCY7 | chr16 | + | ENSG000001 | ENSG000001 | TRUE | cassette^1\| | TRUE | C1_A | chr16:50246 | FALSE | TRUE | TRUE | 0.002 | 0.317 | 0.296 | 1 | 0 | cassette | 0.296 |
| 94 | ENSG000001 | ENSG000001 | GLIPR2 | chr9 | + | ENSG000001 | ENSG000001 | FALSE | ale^1 | FALSE | Distal | chr9:361486 | FALSE | FALSE | FALSE | 0.026 | 0.211 | 0.154 | 0.918 | 0.017 | AFE/ALE | 0.154 |
| 95 | ENSG000001 | ENSG000001 | CNTFR | chr9 | - | ENSG000001 | ENSG000001 | TRUE | cassette^2\|afe^4\|putativ | | C1_A | chr9: | FALSE | FALSE | TRUE | 0.004 | 0.165 | 0.138 | 0.992 | 0.003 | cassette | 0.138 |
| 96 | ENSG000001 | ENSG000001 | CYREN | chr7 | - | ENSG000001 | ENSG000001 | TRUE | alt5^1\|multi | FALSE | C1_A | chr7:135167 | FALSE | FALSE | TRUE | 0.006 | 0.17 | 0.138 | 0.978 | 0.008 | cassette | 0.138 |
| 97 | ENSG000001 | ENSG000001 | MED13L | chr12 | - | ENSG000001 | ENSG000001 | FALSE | cassette^1 | TRUE | C1_A | chr12:11605 | FALSE | FALSE | TRUE | 0.001 | 0.147 | 0.124 | 0.996 | 0.002 | cassette | 0.124 |
| 98 | ENSG000001 | ENSG000001 | STX16 | chr20 | + | ENSG000001 | ENSG000001 | FALSE | cassette^1 | FALSE | C2_C1 | chr20:58670 | FALSE | FALSE | TRUE | 0.014 | 0.126 | -0.101 | 0.951 | 0.025 | skipping | 0.101 |
| 99 | ENSG000001 | ENSG000001 | ATXN1 | chr6 | - | ENSG000001 | ENSG000001 | TRUE | cassette^3\| | TRUE | C1_A | chr6:165895 | FALSE | FALSE | TRUE | 0.001 | 0.366 | 0.347 | 1 | 0 | cassette | 0.347 |
| 100 | ENSG000001 | ENSG000001 | ATXN1 | chr6 | - | ENSG000001 | ENSG000001 | TRUE | cassette^3\|alt5^3\|ir^2\|a | | A_C2 | chr6: | FALSE | FALSE | TRUE | 0.001 | 0.184 | 0.161 | 1 | 0 | cassette | 0.161 |
| 101 | ENSG000001 | ENSG000001 | TM9SF2 | chr13 | + | ENSG000001 | ENSG000001 | TRUE | alt3^1\|ir^5\| | TRUE | Proximal | chr13:99536 | FALSE | FALSE | TRUE | 0 | 0.149 | 0.127 | 0.96 | 0.017 | AFE/ALE | 0.127 |
| 102 | ENSG000001 | ENSG000001 | TM9SF2 | chr13 | + | ENSG000001 | ENSG000001 | TRUE | alt3^1\|ir^5\| | TRUE | Distal | chr13:99536 | FALSE | FALSE | TRUE | 0.001 | 0.143 | 0.119 | 0.953 | 0.021 | AFE/ALE | 0.119 |
| 103 | ENSG000001 | ENSG000001 | TM9SF2 | chr13 | + | ENSG000001 | ENSG000001 | TRUE | alt3^1\|ir^5\| | TRUE | Distal | chr13:99536 | FALSE | FALSE | FALSE | 0.001 | 0.139 | 0.115 | 0.949 | 0.023 | AFE/ALE | 0.115 |
| 104 | ENSG000001 | ENSG000001 | TMEM175 | chr4 | + | ENSG000001 | ENSG000001 | TRUE | cassette^1\|i | FALSE | C1_A | chr4:932540 | FALSE | FALSE | TRUE | 0 | 0.165 | 0.145 | 1 | 0 | cassette | 0.145 |
| 105 | ENSG000001 | ENSG000001 | MACF1 | chr1 | + | ENSG000001 | ENSG000001 | TRUE | cassette^1\|i | FALSE | C1_A | chr1:394357 | FALSE | FALSE | TRUE | 0.036 | 0.13 | 0.099 | 0.957 | 0.023 | cassette | 0.099 |
| 106 | ENSG000001 | ENSG000001 | LRFN1 | chr19 | - | ENSG000001 | ENSG000001 | TRUE | cassette^1\|i | TRUE | C1_A | chr19:39316 | FALSE | FALSE | TRUE | 0.001 | 0.144 | 0.119 | 0.996 | 0.002 | cassette | 0.119 |
| 107 | ENSG000001 | ENSG000001 | EPB41L4A | chr5 | - | ENSG000001 | ENSG000001 | FALSE | cassette^1 | TRUE | C1_A | chr5:112267 | FALSE | TRUE | TRUE | 0.024 | 0.895 | 0.882 | 1 | 0 | cassette | 0.882 |
| 108 | ENSG000001 | ENSG000001 | CDO1 | chr5 | - | ENSG000001 | ENSG000001 | TRUE | cassette^1\|i | FALSE | C1_A | chr5:115813 | FALSE | TRUE | TRUE | 0.022 | 0.315 | 0.295 | 1 | 0 | cassette | 0.295 |
| 109 | ENSG000001 | ENSG000001 | UNC13A | chr19 | - | ENSG000001 | ENSG000001 | TRUE | cassette^2\| | TRUE | C1_A | chr19:17642 | FALSE | TRUE | TRUE | 0 | 0.492 | 0.479 | 1 | 0 | cassette | 0.479 |
| 110 | ENSG000001 | ENSG000001 | UNC13A | chr19 | - | ENSG000001 | ENSG000001 | TRUE | cassette^2\| | TRUE | C2_A | chr19:17641 | FALSE | TRUE | TRUE | 0 | 0.42 | 0.403 | 1 | 0 | cassette | 0.403 |
| 111 | ENSG000001 | ENSG000001 | PXDN | chr2 | - | ENSG000001 | ENSG000001 | TRUE | cassette^1\|i | TRUE | C1_A | chr2:165088 | FALSE | TRUE | TRUE | 0.003 | 0.449 | 0.436 | 1 | 0 | cassette | 0.436 |
| 112 | ENSG000001 | ENSG000001 | CAMSAP1 | chr9 | - | ENSG000001 | ENSG000001 | TRUE | putative_afe | TRUE | Proximal | chr9:135838 | FALSE | FALSE | TRUE | 0.003 | 0.151 | 0.124 | 0.972 | 0.012 | AFE/ALE | 0.124 |
| 113 | ENSG000001 | ENSG000001 | SYNE1 | chr6 | - | ENSG000001 | ENSG000001 | FALSE | cassette^1 | TRUE | C1_A | chr6:152247 | FALSE | TRUE | TRUE | 0.001 | 0.216 | 0.188 | 1 | 0 | cassette | 0.188 |
| 114 | ENSG000001 | ENSG000001 | ZNF141 | chr4 | + | ENSG000001 | ENSG000001 | TRUE | ir^1\|putative | TRUE | Proximal | chr4:373234 | FALSE | TRUE | TRUE | 0.003 | 0.346 | 0.32 | 1 | 0 | AFE/ALE | 0.32 |
| 115 | ENSG000001 | ENSG000001 | GSE1 | chr16 | + | ENSG000001 | ENSG000001 | TRUE | cassette^2\| | TRUE | Orphan | chr16:85463 | FALSE | FALSE | TRUE | 0.003 | 0.167 | 0.145 | 0.995 | 0.002 | cassette | 0.145 |
| 116 | ENSG000001 | ENSG000001 | MGAT1 | chr5 | - | ENSG000001 | ENSG000001 | TRUE | alt5^3\|afe^1 | TRUE | Distal | chr5:180809 | FALSE | TRUE | TRUE | 0.001 | 0.3 | 0.278 | 1 | 0 | AFE/ALE | 0.278 |
| 117 | ENSG000001 | ENSG000001 | NUP210 | chr3 | - | ENSG000001 | ENSG000001 | TRUE | tandem_cass | FALSE | C1_A | chr3:133738 | FALSE | TRUE | TRUE | 0.014 | 0.304 | 0.279 | 1 | 0 | cassette | 0.279 |
| 118 | ENSG000001 | ENSG000001 | RIN2 | chr20 | + | ENSG000001 | ENSG000001 | TRUE | cassette^1\|i | TRUE | C1_A | chr20:19965 | FALSE | TRUE | TRUE | 0.001 | 0.241 | 0.222 | 1 | 0 | cassette | 0.222 |
| 119 | ENSG000001 | ENSG000001 | XPO4 | chr13 | - | ENSG000001 | ENSG000001 | FALSE | alt5^1 | TRUE | Proximal | chr13:20799 | FALSE | FALSE | TRUE | 0.013 | 0.205 | 0.18 | 0.994 | 0.002 | AFE/ALE | 0.18 |
| 120 | ENSG000001 | ENSG000001 | CDK7 | chr5 | + | ENSG000001 | ENSG000001 | TRUE | cassette^1\|i | TRUE | C2_A | chr5:692614 | FALSE | FALSE | TRUE | 0.001 | 0.28 | 0.258 | 1 | 0 | cassette | 0.258 |
| 121 | ENSG000001 | ENSG000001 | KATNBL1 | chr15 | - | ENSG000001 | ENSG000001 | TRUE | ale^4\|multi_ | FALSE | Proximal | chr15:34143 | FALSE | FALSE | TRUE | 0.013 | 0.174 | 0.15 | 0.988 | 0.004 | AFE/ALE | 0.15 |
| 122 | ENSG000001 | ENSG000001 | ARHGAP32 | chr11 | - | ENSG000001 | ENSG000001 | FALSE | putative_ale | TRUE | Proximal | chr11:12899 | FALSE | TRUE | TRUE | 0.009 | 0.709 | 0.703 | 1 | 0 | AFE/ALE | 0.703 |
| 123 | ENSG000001 | ENSG000001 | TAOK3 | chr12 | - | ENSG000001 | ENSG000001 | FALSE | cassette^1 | TRUE | C1_A | chr12:11815 | FALSE | FALSE | TRUE | 0.019 | 0.165 | 0.144 | 1 | 0 | cassette | 0.144 |
| 124 | ENSG000001 | ENSG000001 | MICAL1 | chr6 | - | ENSG000001 | ENSG000001 | TRUE | cassette^6\| | TRUE | C2_A | chr6:109447 | FALSE | FALSE | TRUE | 0.007 | 0.138 | 0.111 | 0.976 | 0.012 | cassette | 0.111 |
| 125 | ENSG000001 | ENSG000001 | TGFBRAP1 | chr2 | - | ENSG000001 | ENSG000001 | FALSE | alt5^1 | TRUE | Proximal | chr2:105298 | FALSE | FALSE | TRUE | 0.007 | 0.137 | 0.113 | 0.955 | 0.021 | AFE/ALE | 0.113 |
| 126 | ENSG000001 | ENSG000001 | ANKRD36 | chr2 | + | ENSG000001 | ENSG000001 | TRUE | cassette^1\|i | TRUE | C1_C2 | chr2:972476 | FALSE | FALSE | TRUE | 0.007 | 0.249 | -0.221 | 1 | 0 | skipping | 0.221 |
| 127 | ENSG000001 | ENSG000001 | LMO7 | chr13 | + | ENSG000001 | ENSG000001 | TRUE | cassette^1\| | FALSE | Distal | chr13:75804 | FALSE | FALSE | FALSE | 0.022 | 0.2 | 0.152 | 0.921 | 0.016 | AFE/ALE | 0.152 |
| 128 | ENSG000001 | ENSG000001 | CBWD2 | chr2 | + | ENSG000001 | ENSG000001 | TRUE | ir^2\|putative | TRUE | Proximal | chr2:113465 | FALSE | FALSE | TRUE | 0.02 | 0.2 | 0.162 | 0.988 | 0.003 | AFE/ALE | 0.162 |
| 129 | ENSG000001 | ENSG000001 | SEMA6D | chr15 | + | ENSG000001 | ENSG000001 | TRUE | cassette^6\| | TRUE | C1_A | chr15:47412 | FALSE | TRUE | TRUE | 0.002 | 0.217 | 0.197 | 1 | 0 | cassette | 0.197 |
| 130 | ENSG000001 | ENSG000001 | ACTR1A | chr10 | - | ENSG000001 | ENSG000001 | TRUE | ir^1\|putative | TRUE | Proximal | chr10:10247 | FALSE | FALSE | TRUE | 0.011 | 0.181 | 0.152 | 0.984 | 0.004 | AFE/ALE | 0.152 |
| 131 | ENSG000001 | ENSG000001 | SEPTIN11 | chr4 | + | ENSG000001 | ENSG000001 | TRUE | cassette^1\|i | TRUE | C1_A | chr4:770309 | FALSE | FALSE | TRUE | 0.005 | 0.171 | 0.152 | 1 | 0 | cassette | 0.152 |
| 132 | ENSG000001 | ENSG000001 | KIF21A | chr12 | - | ENSG000001 | ENSG000001 | TRUE | cassette^1\|i | FALSE | C1_C2 | chr12:39326 | FALSE | FALSE | FALSE | 0.03 | 0.119 | -0.096 | 0.932 | 0.036 | skipping | 0.096 |
| 133 | ENSG000001 | ENSG000001 | KIF21A | chr12 | - | ENSG000001 | ENSG000001 | TRUE | cassette^1\| | TRUE | C1_A | chr12:39389 | FALSE | FALSE | TRUE | 0.001 | 0.248 | 0.224 | 1 | 0 | cassette | 0.224 |
| 134 | ENSG000001 | ENSG000001 | KIF21A | chr12 | - | ENSG000001 | ENSG000001 | TRUE | cassette^1\| | TRUE | Proximal | chr12:39370 | FALSE | FALSE | TRUE | 0.001 | 0.135 | 0.11 | 1 | 0 | AFE/ALE | 0.11 |
| 135 | ENSG000001 | ENSG000001 | TMEM117 | chr12 | + | ENSG000001 | ENSG000001 | FALSE | ale^1 | TRUE | Proximal | chr12:43836 | FALSE | TRUE | TRUE | 0.006 | 0.812 | 0.796 | 1 | 0 | AFE/ALE | 0.796 |
| 136 | ENSG000001 | ENSG000001 | SRRM4 | chr12 | + | ENSG000001 | ENSG000001 | TRUE | cassette^2\|ir^6\|multi_ex | | A_C2 | chr12: | FALSE | FALSE | TRUE | 0.013 | 0.195 | 0.162 | 0.997 | 0.001 | cassette | 0.162 |
| 137 | ENSG000001 | ENSG000001 | POLG | chr15 | - | ENSG000001 | ENSG000001 | FALSE | cassette^1 | FALSE | C2_C1 | chr15:89322 | FALSE | FALSE | TRUE | 0.02 | 0.189 | -0.154 | 0.972 | 0.007 | skipping | 0.154 |
| 138 | ENSG000001 | ENSG000001 | NTRK3 | chr15 | - | ENSG000001 | ENSG000001 | TRUE | alt5^1\|ir^2 | TRUE | Proximal | chr15:88256 | FALSE | FALSE | TRUE | 0.001 | 0.132 | 0.105 | 0.955 | 0.023 | AFE/ALE | 0.105 |
| 139 | ENSG000001 | ENSG000001 | NKD1 | chr16 | + | ENSG000001 | ENSG000001 | TRUE | cassette^2\| | TRUE | C1_A | chr16:50608 | FALSE | FALSE | TRUE | 0.001 | 0.408 | 0.391 | 1 | 0 | cassette | 0.391 |
| 140 | ENSG000001 | ENSG000001 | NKD1 | chr16 | + | ENSG000001 | ENSG000001 | TRUE | cassette^2\| | TRUE | Proximal | chr16:50620 | FALSE | FALSE | TRUE | 0.001 | 0.456 | 0.443 | 1 | 0 | AFE/ALE | 0.443 |
| 141 | ENSG000001 | ENSG000001 | PRELID3A | chr18 | + | ENSG000001 | ENSG000001 | TRUE | cassette^1\| | TRUE | C1_A | chr18:12429 | FALSE | TRUE | TRUE | 0.003 | 0.529 | 0.512 | 1 | 0 | cassette | 0.512 |
| 142 | ENSG000001 | ENSG000001 | CEP131 | chr17 | - | ENSG000001 | ENSG000001 | TRUE | putative_afe | TRUE | Distal | chr17:81207 | FALSE | FALSE | TRUE | 0.001 | 0.17 | 0.144 | 0.995 | 0.002 | AFE/ALE | 0.144 |
| 143 | ENSG000001 | ENSG000001 | FBN3 | chr19 | - | ENSG000001 | ENSG000001 | TRUE | cassette^1\|i | TRUE | C1_A | chr19:80844 | FALSE | TRUE | TRUE | 0.002 | 0.74 | 0.733 | 1 | 0 | cassette | 0.733 |
| 144 | ENSG000001 | ENSG000001 | IGLON5 | chr19 | + | ENSG000001 | ENSG000001 | FALSE | cassette^1 | TRUE | C1_A | chr19:51311 | FALSE | TRUE | TRUE | 0.001 | 0.24 | 0.215 | 1 | 0 | cassette | 0.215 |
| 145 | ENSG000001 | ENSG000001 | NVL | chr1 | - | ENSG000001 | ENSG000001 | TRUE | cassette^8\| | TRUE | C2_A | chr1:224311 | FALSE | FALSE | TRUE | 0.036 | 0.185 | 0.151 | 0.974 | 0.008 | cassette | 0.151 |
| 146 | ENSG000001 | ENSG000001 | CNGA3 | chr2 | + | ENSG000001 | ENSG000001 | FALSE | putative_afe | TRUE | Proximal | chr2:983760 | FALSE | TRUE | TRUE | 0.038 | 0.301 | 0.253 | 0.995 | 0 | AFE/ALE | 0.253 |
| 147 | ENSG000001 | ENSG000001 | GOLGA4 | chr3 | + | ENSG000001 | ENSG000001 | TRUE | cassette^2\| | TRUE | C1_A | chr3:372514 | FALSE | FALSE | FALSE | 0.008 | 0.133 | 0.103 | 0.903 | 0.049 | cassette | 0.103 |
| 148 | ENSG000001 | ENSG000001 | SHQ1 | chr3 | - | ENSG000001 | ENSG000001 | TRUE | cassette^1\| | TRUE | Proximal | chr3:727508 | FALSE | FALSE | TRUE | 0.001 | 0.306 | 0.286 | 1 | 0 | AFE/ALE | 0.286 |
| 149 | ENSG000001 | ENSG000001 | STXBP5L | chr3 | + | ENSG000001 | ENSG000001 | TRUE | alt5^3\|afe^3 | TRUE | Proximal | chr3:120908 | FALSE | TRUE | TRUE | 0.001 | 0.222 | 0.193 | 1 | 0 | AFE/ALE | 0.193 |
| 150 | ENSG000001 | ENSG000001 | STXBP5L | chr3 | + | ENSG000001 | ENSG000001 | TRUE | alt5^3\|afe^3 | TRUE | Proximal | chr3:120908 | FALSE | TRUE | TRUE | 0.001 | 0.409 | 0.382 | 1 | 0 | AFE/ALE | 0.382 |
| 151 | ENSG000001 | ENSG000001 | G3BP1 | chr5 | + | ENSG000001 | ENSG000001 | FALSE | cassette^1 | FALSE | C1_A | chr5:151786 | FALSE | TRUE | TRUE | 0.033 | 0.571 | 0.566 | 1 | 0 | cassette | 0.566 |

| | | | | | | | | | | | | | | | | | | | | | | |
|---|---|---|---|---|---|---|---|---|---|---|---|---|---|---|---|---|---|---|---|---|---|---|
| 152 | ENSG000001 | ENSG000001 | SCUBE3 | chr6 | + | ENSG000001 | ENSG000001 | TRUE | cassette^3\|a | TRUE | C2_A | chr6:352181 | FALSE | FALSE | TRUE | 0.045 | 0.23 | 0.191 | 1 | 0 | cassette | 0.191 |
| 153 | ENSG000001 | ENSG000001 | WASH2P | chr2 | + | ENSG000001 | ENSG000001 | FALSE | alt5^1 | TRUE | Proximal | chr2:113598 | FALSE | FALSE | FALSE | 0.009 | 0.167 | 0.137 | 0.946 | 0.017 | AFE/ALE | 0.137 |
| 154 | ENSG000001 | ENSG000001 | UTP23 | chr8 | + | ENSG000001 | ENSG000001 | TRUE | ir^2\|ale^2\|r | FALSE | Distal | chr8:116770 | FALSE | FALSE | FALSE | 0.045 | 0.253 | 0.184 | 0.932 | 0.008 | AFE/ALE | 0.184 |
| 155 | ENSG000001 | ENSG000001 | POLR3A | chr10 | - | ENSG000001 | ENSG000001 | FALSE | cassette^1 | FALSE | C1_C2 | chr10:78007 | FALSE | FALSE | TRUE | 0.028 | 0.214 | -0.184 | 0.999 | 0 | skipping | 0.184 |
| 156 | ENSG000001 | ENSG000001 | INTS4 | chr11 | - | ENSG000001 | ENSG000001 | TRUE | putative_alt5 | TRUE | C1_A | chr11:77883 | FALSE | FALSE | TRUE | 0.031 | 0.742 | 0.72 | 1 | 0 | cassette | 0.72 |
| 157 | ENSG000001 | ENSG000001 | HYOU1 | chr11 | - | ENSG000001 | ENSG000001 | FALSE | cassette^1 | FALSE | C1_C2 | chr11:11905 | FALSE | FALSE | TRUE | 0.02 | 0.176 | -0.155 | 1 | 0 | skipping | 0.155 |
| 158 | ENSG000001 | ENSG000001 | FEZ1 | chr11 | - | ENSG000001 | ENSG000001 | TRUE | alt5^1\|afe^2 | FALSE | Proximal | chr11:12548 | FALSE | TRUE | TRUE | 0.027 | 0.278 | 0.258 | 1 | 0 | AFE/ALE | 0.258 |
| 159 | ENSG000001 | ENSG000001 | UBC | chr12 | - | ENSG000001 | ENSG000001 | TRUE | putative_alt3 | FALSE | C1_A | chr12:12491 | FALSE | FALSE | TRUE | 0.003 | 0.182 | 0.157 | 1 | 0 | cassette | 0.157 |
| 160 | ENSG000001 | ENSG000001 | UBC | chr12 | - | ENSG000001 | ENSG000001 | TRUE | putative_alt3 | FALSE | A_C2 | chr12:12491 | FALSE | FALSE | TRUE | 0.024 | 0.119 | 0.102 | 0.991 | 0.005 | cassette | 0.102 |
| 161 | ENSG000001 | ENSG000001 | DLG5 | chr10 | - | ENSG000001 | ENSG000001 | TRUE | cassette^1\|i | TRUE | C1_A | chr10:77793 | FALSE | FALSE | TRUE | 0.001 | 0.14 | 0.116 | 0.999 | 0 | cassette | 0.116 |
| 162 | ENSG000001 | ENSG000001 | RNF144A | chr2 | + | ENSG000001 | ENSG000001 | TRUE | ale^2\|putati | TRUE | Proximal | chr2:703021 | FALSE | TRUE | TRUE | 0.032 | 0.289 | 0.261 | 1 | 0 | AFE/ALE | 0.261 |
| 163 | ENSG000001 | ENSG000001 | RNF144A | chr2 | + | ENSG000001 | ENSG000001 | TRUE | ale^2\|putati | TRUE | Distal | chr2:703021 | FALSE | FALSE | FALSE | 0.036 | 0.152 | 0.112 | 0.902 | 0.039 | AFE/ALE | 0.112 |
| 164 | ENSG000001 | ENSG000001 | GUF1 | chr4 | + | ENSG000001 | ENSG000001 | TRUE | ir^2\|putative | TRUE | Proximal | chr4:446927 | FALSE | FALSE | TRUE | 0.034 | 0.19 | 0.157 | 0.983 | 0.005 | AFE/ALE | 0.157 |
| 165 | ENSG000001 | ENSG000001 | PTPRD | chr9 | - | ENSG000001 | ENSG000001 | TRUE | cassette^4\|t | TRUE | C1_A | chr9:912720 | FALSE | FALSE | TRUE | 0.006 | 0.612 | 0.602 | 1 | 0 | cassette | 0.602 |
| 166 | ENSG000001 | ENSG000001 | PTPRD | chr9 | - | ENSG000001 | ENSG000001 | TRUE | cassette^4\|t | TRUE | C1_A | chr9:912720 | FALSE | TRUE | TRUE | 0.019 | 0.645 | 0.634 | 1 | 0 | cassette | 0.634 |
| 167 | ENSG000001 | ENSG000001 | PTPRD | chr9 | - | ENSG000001 | ENSG000001 | TRUE | cassette^4\|t | TRUE | C2_A_Last | chr9:901873 | FALSE | FALSE | TRUE | 0.026 | 0.575 | 0.566 | 1 | 0 | cassette | 0.566 |
| 168 | ENSG000001 | ENSG000001 | PTPRD | chr9 | - | ENSG000001 | ENSG000001 | TRUE | cassette^3\|a | TRUE | E2_E1_J1 | chr9:939658 | FALSE | FALSE | TRUE | 0 | 0.173 | 0.144 | 0.958 | 0.008 | cassette | 0.144 |
| 169 | ENSG000001 | ENSG000001 | PTPRD | chr9 | - | ENSG000001 | ENSG000001 | TRUE | cassette^3\|a | TRUE | Proximal | chr9:944222 | FALSE | TRUE | TRUE | 0 | 0.314 | 0.29 | 1 | 0 | AFE/ALE | 0.29 |
| 170 | ENSG000001 | ENSG000001 | UBASH3B | chr11 | + | ENSG000001 | ENSG000001 | FALSE | cassette^1 | TRUE | C1_A | chr11:12265 | FALSE | TRUE | TRUE | 0.002 | 0.573 | 0.56 | 1 | 0 | cassette | 0.56 |
| 171 | ENSG000001 | ENSG000001 | PTPRN2 | chr7 | - | ENSG000001 | ENSG000001 | TRUE | putative_ale | TRUE | Proximal | chr7:157875 | FALSE | TRUE | TRUE | 0.003 | 0.245 | 0.208 | 0.996 | 0.001 | AFE/ALE | 0.208 |
| 172 | ENSG000001 | ENSG000001 | PTPRN2 | chr7 | - | ENSG000001 | ENSG000001 | TRUE | ale^3\|afe^1\|multi_exon_s | | C1_A | chr7: | FALSE | FALSE | TRUE | 0.001 | 0.328 | 0.304 | 1 | 0 | cassette | 0.304 |
| 173 | ENSG000001 | ENSG000001 | PTPRN2 | chr7 | - | ENSG000001 | ENSG000001 | TRUE | cassette^3\|p | TRUE | C1_A | chr7:158249 | FALSE | FALSE | TRUE | 0.02 | 0.19 | 0.165 | 1 | 0 | cassette | 0.165 |
| 174 | ENSG000001 | ENSG000001 | GOLGA7B | chr10 | + | ENSG000001 | ENSG000001 | TRUE | cassette^1\|i | TRUE | C1_A | chr10:97866 | FALSE | FALSE | TRUE | 0.003 | 0.349 | 0.325 | 1 | 0 | cassette | 0.325 |
| 175 | ENSG000001 | ENSG000001 | GOLGA7B | chr10 | + | ENSG000001 | ENSG000001 | TRUE | cassette^1\|i | TRUE | Proximal | chr10:97866 | FALSE | FALSE | TRUE | 0.015 | 0.401 | 0.382 | 1 | 0 | AFE/ALE | 0.382 |
| 176 | ENSG000001 | ENSG000001 | ADCY8 | chr8 | - | ENSG000001 | ENSG000001 | FALSE | putative_ale | TRUE | Proximal | chr8:130909 | FALSE | TRUE | TRUE | 0.011 | 0.358 | 0.325 | 1 | 0 | AFE/ALE | 0.325 |
| 177 | ENSG000001 | ENSG000001 | KCNMA1 | chr10 | - | ENSG000001 | ENSG000001 | TRUE | cassette^1\|i | FALSE | C1_C2 | chr10:77019 | FALSE | FALSE | TRUE | 0.003 | 0.187 | -0.163 | 0.988 | 0.003 | skipping | 0.163 |
| 178 | ENSG000001 | ENSG000001 | ETS2 | chr21 | + | ENSG000001 | ENSG000001 | FALSE | cassette^1 | FALSE | C2_A | chr21:38806 | FALSE | FALSE | TRUE | 0.002 | 0.2 | 0.175 | 1 | 0 | cassette | 0.175 |
| 179 | ENSG000001 | ENSG000001 | TSPAN18 | chr11 | + | ENSG000001 | ENSG000001 | FALSE | putative_alt5 | TRUE | Intron | chr11:44920 | FALSE | FALSE | TRUE | 0.048 | 0.239 | 0.181 | 0.985 | 0.003 | cassette | 0.181 |
| 180 | ENSG000001 | ENSG000001 | WIPI2 | chr7 | + | ENSG000001 | ENSG000001 | FALSE | cassette^1 | TRUE | C2_C1 | chr7:519049 | FALSE | FALSE | FALSE | 0.03 | 0.143 | -0.111 | 0.935 | 0.029 | skipping | 0.111 |
| 181 | ENSG000001 | ENSG000001 | UBXN11 | chr1 | - | ENSG000001 | ENSG000001 | TRUE | cassette^5\|t | FALSE | A_C2 | chr1:262943 | FALSE | FALSE | FALSE | 0.046 | 0.188 | 0.13 | 0.905 | 0.03 | cassette | 0.13 |
| 182 | ENSG000001 | ENSG000001 | EYA3 | chr1 | - | ENSG000001 | ENSG000001 | FALSE | cassette^1 | TRUE | C1_A | chr1:280566 | FALSE | TRUE | TRUE | 0.002 | 0.217 | 0.195 | 1 | 0 | cassette | 0.195 |
| 183 | ENSG000001 | ENSG000001 | ISL2 | chr15 | + | ENSG000001 | ENSG000001 | TRUE | afe^4\|multi_ | FALSE | A_C2 | chr15:76338 | FALSE | FALSE | TRUE | 0.004 | 0.278 | 0.258 | 1 | 0 | cassette | 0.258 |
| 184 | ENSG000001 | ENSG000001 | KALRN | chr3 | + | ENSG000001 | ENSG000001 | TRUE | cassette^2\|a | TRUE | C1_A | chr3:124700 | FALSE | FALSE | TRUE | 0.006 | 0.141 | 0.116 | 0.994 | 0.003 | cassette | 0.116 |
| 185 | ENSG000001 | ENSG000001 | KALRN | chr3 | + | ENSG000001 | ENSG000001 | TRUE | cassette^2\|a | TRUE | C1_A | chr3:124700 | FALSE | TRUE | TRUE | 0.014 | 0.308 | 0.289 | 1 | 0 | cassette | 0.289 |
| 186 | ENSG000001 | ENSG000001 | WDR4 | chr21 | - | ENSG000001 | ENSG000001 | TRUE | ir^6\|putative | FALSE | Distal | chr21:42843 | FALSE | FALSE | TRUE | 0.012 | 0.174 | 0.149 | 0.993 | 0.002 | AFE/ALE | 0.149 |
| 187 | ENSG000001 | ENSG000001 | LSS | chr21 | - | ENSG000001 | ENSG000001 | TRUE | putative_alt3 | TRUE | Distal | chr21:46193 | FALSE | FALSE | TRUE | 0.028 | 0.288 | 0.264 | 1 | 0 | AFE/ALE | 0.264 |
| 188 | ENSG000001 | ENSG000001 | LSS | chr21 | - | ENSG000001 | ENSG000001 | TRUE | cassette^1\|ir^2\|multi_ex | | C1_A | chr21: | FALSE | FALSE | TRUE | 0.031 | 0.145 | 0.12 | 0.985 | 0.007 | cassette | 0.12 |
| 189 | ENSG000001 | ENSG000001 | TAOK1 | chr17 | + | ENSG000001 | ENSG000001 | FALSE | cassette^1 | TRUE | C1_A | chr17:29534 | FALSE | FALSE | TRUE | 0 | 0.166 | 0.149 | 1 | 0 | cassette | 0.149 |
| 190 | ENSG000001 | ENSG000001 | CELF5 | chr19 | + | ENSG000001 | ENSG000001 | TRUE | cassette^5\|a | TRUE | C1_A | chr19:32249 | FALSE | FALSE | TRUE | 0.003 | 0.146 | 0.121 | 0.999 | 0.001 | cassette | 0.121 |
| 191 | ENSG000001 | ENSG000001 | CELF5 | chr19 | + | ENSG000001 | ENSG000001 | TRUE | cassette^5\|a | TRUE | C1_A | chr19:32249 | FALSE | FALSE | TRUE | 0.001 | 0.173 | 0.149 | 1 | 0 | cassette | 0.149 |
| 192 | ENSG000001 | ENSG000001 | CELF5 | chr19 | + | ENSG000001 | ENSG000001 | TRUE | cassette^5\|a | TRUE | C1_A | chr19:32256 | FALSE | FALSE | TRUE | 0.001 | 0.231 | 0.201 | 1 | 0 | cassette | 0.201 |
| 193 | ENSG000001 | ENSG000001 | CELF5 | chr19 | + | ENSG000001 | ENSG000001 | TRUE | cassette^5\|a | TRUE | C1_A | chr19:32256 | FALSE | FALSE | TRUE | 0.009 | 0.347 | 0.329 | 1 | 0 | cassette | 0.329 |
| 194 | ENSG000001 | ENSG000001 | CELF5 | chr19 | + | ENSG000001 | ENSG000001 | TRUE | cassette^2\|a | TRUE | E1_E2_J1 | chr19:32781 | FALSE | FALSE | TRUE | 0 | 0.149 | 0.123 | 0.999 | 0 | cassette | 0.123 |
| 195 | ENSG000001 | ENSG000001 | CELF5 | chr19 | + | ENSG000001 | ENSG000001 | TRUE | cassette^2\|a | TRUE | C1_A | chr19:32781 | FALSE | FALSE | TRUE | 0 | 0.16 | 0.14 | 1 | 0 | cassette | 0.14 |
| 196 | ENSG000001 | ENSG000001 | CELF5 | chr19 | + | ENSG000001 | ENSG000001 | TRUE | cassette^2\|a | TRUE | C1_A | chr19:32781 | FALSE | FALSE | TRUE | 0 | 0.163 | 0.141 | 1 | 0 | cassette | 0.141 |
| 197 | ENSG000001 | ENSG000001 | TAL1 | chr1 | - | ENSG000001 | ENSG000001 | FALSE | afe^1 | TRUE | Distal | chr1:472240 | FALSE | TRUE | TRUE | 0.021 | 0.27 | 0.247 | 1 | 0 | AFE/ALE | 0.247 |
| 198 | ENSG000001 | ENSG000001 | USP24 | chr1 | - | ENSG000001 | ENSG000001 | FALSE | cassette^1 | TRUE | C1_C2 | chr1:550786 | FALSE | TRUE | TRUE | 0.017 | 0.238 | -0.214 | 1 | 0 | skipping | 0.214 |
| 199 | ENSG000001 | ENSG000001 | HFM1 | chr1 | - | ENSG000001 | ENSG000001 | TRUE | cassette^1\|a | TRUE | E1_E2_J1 | chr1:913874 | FALSE | FALSE | FALSE | 0.001 | 0.232 | 0.185 | 0.93 | 0.011 | cassette | 0.185 |
| 200 | ENSG000001 | ENSG000001 | HFM1 | chr1 | - | ENSG000001 | ENSG000001 | TRUE | cassette^1\|a | TRUE | E1_E2_J1 | chr1:913875 | FALSE | FALSE | TRUE | 0.001 | 0.196 | 0.172 | 1 | 0 | cassette | 0.172 |
| 201 | ENSG000001 | ENSG000001 | HFM1 | chr1 | - | ENSG000001 | ENSG000001 | TRUE | cassette^1\|alt3^20\|alt5^ | | C1_A | chr1: | FALSE | FALSE | TRUE | 0.001 | 0.237 | 0.211 | 1 | 0 | cassette | 0.211 |
| 202 | ENSG000001 | ENSG000001 | HFM1 | chr1 | - | ENSG000001 | ENSG000001 | TRUE | cassette^1\|a | TRUE | Distal | chr1:913873 | FALSE | FALSE | TRUE | 0.001 | 0.187 | 0.163 | 0.994 | 0.002 | AFE/ALE | 0.163 |
| 203 | ENSG000001 | ENSG000001 | KCNT2 | chr1 | - | ENSG000001 | ENSG000001 | TRUE | cassette^1\|p | TRUE | Proximal | chr1:196305 | FALSE | FALSE | TRUE | 0.004 | 0.152 | 0.123 | 0.966 | 0.015 | AFE/ALE | 0.123 |
| 204 | ENSG000001 | ENSG000001 | KCNT2 | chr1 | - | ENSG000001 | ENSG000001 | TRUE | ir^2\|afe^4\|r | TRUE | A_C2 | chr1:196316 | FALSE | FALSE | TRUE | 0.011 | 0.209 | 0.181 | 0.999 | 0 | cassette | 0.181 |
| 205 | ENSG000001 | ENSG000001 | KIF26B | chr1 | + | ENSG000001 | ENSG000001 | TRUE | putative_ale | TRUE | Proximal | chr1:245419 | FALSE | TRUE | TRUE | 0.002 | 0.274 | 0.251 | 1 | 0 | AFE/ALE | 0.251 |
| 206 | ENSG000001 | ENSG000001 | SANBR | chr2 | + | ENSG000001 | ENSG000001 | TRUE | cassette^1\|i | TRUE | C1_A | chr2:610884 | FALSE | FALSE | TRUE | 0.006 | 0.14 | 0.116 | 0.965 | 0.016 | cassette | 0.116 |
| 207 | ENSG000001 | ENSG000001 | ANKRD30BL | chr2 | - | ENSG000001 | ENSG000001 | TRUE | alt5^3\|orpha | TRUE | Distal | chr2:132255 | FALSE | FALSE | FALSE | 0 | 0.137 | 0.109 | 0.938 | 0.03 | AFE/ALE | 0.109 |
| 208 | ENSG000001 | ENSG000001 | CLASP2 | chr3 | - | ENSG000001 | ENSG000001 | TRUE | cassette^2\|t | TRUE | C1_A | chr3:335887 | FALSE | FALSE | TRUE | 0.043 | 0.203 | 0.172 | 1 | 0 | cassette | 0.172 |
| 209 | ENSG000001 | ENSG000001 | IFT122 | chr3 | + | ENSG000001 | ENSG000001 | TRUE | ir^2\|ale^2\|p | TRUE | Proximal | chr3:129483 | FALSE | FALSE | TRUE | 0.003 | 0.366 | 0.335 | 1 | 0 | AFE/ALE | 0.335 |
| 210 | ENSG000001 | ENSG000001 | IFT122 | chr3 | + | ENSG000001 | ENSG000001 | TRUE | ir^2\|ale^2\|p | TRUE | Proximal | chr3:129483 | FALSE | FALSE | TRUE | 0.003 | 0.178 | 0.157 | 0.994 | 0.002 | AFE/ALE | 0.157 |
| 211 | ENSG000001 | ENSG000001 | SLC9B2 | chr4 | - | ENSG000001 | ENSG000001 | TRUE | cassette^1\|a | FALSE | Proximal | chr4:103065 | FALSE | FALSE | TRUE | 0.007 | 0.221 | 0.197 | 1 | 0 | AFE/ALE | 0.197 |
| 212 | ENSG000001 | ENSG000001 | MAP9 | chr4 | - | ENSG000001 | ENSG000001 | TRUE | cassette^2\|a | FALSE | C2_C1 | chr4:155375 | FALSE | FALSE | TRUE | 0.019 | 0.163 | -0.132 | 0.957 | 0.015 | skipping | 0.132 |
| 213 | ENSG000001 | ENSG000001 | STARD4 | chr5 | - | ENSG000001 | ENSG000001 | FALSE | cassette^1 | FALSE | C1_C2 | chr5:111501 | FALSE | FALSE | TRUE | 0.028 | 0.22 | -0.182 | 0.986 | 0.003 | skipping | 0.182 |
| 214 | ENSG000001 | ENSG000001 | ADCY1 | chr7 | + | ENSG000001 | ENSG000001 | FALSE | cassette^1 | TRUE | C1_A | chr7:456227 | FALSE | TRUE | TRUE | 0.004 | 0.56 | 0.548 | 1 | 0 | cassette | 0.548 |
| 215 | ENSG000001 | ENSG000001 | INTS8 | chr8 | + | ENSG000001 | ENSG000001 | TRUE | cassette^1\|i | TRUE | C1_A | chr8:948321 | FALSE | FALSE | FALSE | 0.001 | 0.121 | 0.097 | 0.912 | 0.048 | cassette | 0.097 |
| 216 | ENSG000001 | ENSG000001 | SLITRK5 | chr13 | + | ENSG000001 | ENSG000001 | TRUE | cassette^1\|i | TRUE | C1_A | chr13:87672 | FALSE | FALSE | TRUE | 0.002 | 0.136 | 0.107 | 0.951 | 0.024 | cassette | 0.107 |
| 217 | ENSG000001 | ENSG000001 | SLC39A13 | chr11 | + | ENSG000001 | ENSG000001 | TRUE | alt5^1\|ir^2 | TRUE | Proximal | chr11:47414 | FALSE | FALSE | FALSE | 0.025 | 0.141 | 0.107 | 0.924 | 0.032 | AFE/ALE | 0.107 |
| 218 | ENSG000001 | ENSG000001 | RIMKLB | chr12 | + | ENSG000001 | ENSG000001 | TRUE | alt5^3 | TRUE | Proximal | chr12:87770 | FALSE | FALSE | FALSE | 0.031 | 0.192 | 0.151 | 0.936 | 0.014 | AFE/ALE | 0.151 |
| 219 | ENSG000001 | ENSG000001 | ISLR2 | chr15 | + | ENSG000001 | ENSG000001 | TRUE | cassette^4\|a | TRUE | C2_A | chr15:74130 | FALSE | FALSE | TRUE | 0.03 | 0.161 | 0.143 | 1 | 0 | cassette | 0.143 |
| 220 | ENSG000001 | ENSG000001 | LENG8 | chr19 | + | ENSG000001 | ENSG000001 | TRUE | cassette^2\|p | TRUE | Distal | chr19:54459 | FALSE | FALSE | TRUE | 0.03 | 0.138 | 0.108 | 0.973 | 0.013 | AFE/ALE | 0.108 |
| 221 | ENSG000001 | ENSG000001 | HDGFL2 | chr19 | + | ENSG000001 | ENSG000001 | TRUE | cassette^1\|a | TRUE | C1_A | chr19:44918 | FALSE | TRUE | TRUE | 0.001 | 0.744 | 0.736 | 1 | 0 | cassette | 0.736 |
| 222 | ENSG000001 | ENSG000001 | HDGFL2 | chr19 | + | ENSG000001 | ENSG000001 | TRUE | cassette^1\|a | TRUE | Proximal | chr19:44921 | FALSE | FALSE | TRUE | 0.001 | 0.522 | 0.505 | 1 | 0 | AFE/ALE | 0.505 |
| 223 | ENSG000001 | ENSG000001 | SLC43A2 | chr17 | - | ENSG000001 | ENSG000001 | TRUE | ir^1\|ale^2\|a | TRUE | Distal | chr17:15840 | FALSE | FALSE | TRUE | 0.001 | 0.391 | 0.369 | 1 | 0 | AFE/ALE | 0.369 |
| 224 | ENSG000001 | ENSG000001 | MGAT5B | chr17 | + | ENSG000001 | ENSG000001 | TRUE | cassette^1\|i | TRUE | C1_A | chr17:76933 | FALSE | TRUE | TRUE | 0.001 | 0.273 | 0.241 | 1 | 0 | cassette | 0.241 |
| 225 | ENSG000001 | ENSG000001 | ANKS3 | chr16 | - | ENSG000001 | ENSG000001 | TRUE | cassette^5\|a | FALSE | C1_A | chr16:47141 | FALSE | FALSE | FALSE | 0.02 | 0.145 | 0.109 | 0.905 | 0.04 | cassette | 0.109 |
| 226 | ENSG000001 | ENSG000001 | ANKS3 | chr16 | - | ENSG000001 | ENSG000001 | TRUE | cassette^5\|a | TRUE | C2_A1 | chr16:47052 | FALSE | FALSE | TRUE | 0.024 | 0.185 | 0.152 | 0.977 | 0.007 | cassette | 0.152 |
| 227 | ENSG000001 | ENSG000001 | SETD5 | chr3 | + | ENSG000001 | ENSG000001 | TRUE | cassette^2\|a | TRUE | C2_A | chr3:946861 | FALSE | FALSE | TRUE | 0.007 | 0.332 | 0.315 | 1 | 0 | cassette | 0.315 |

| | | | | | | | | | | | | | | | | | | | | | | |
|---|---|---|---|---|---|---|---|---|---|---|---|---|---|---|---|---|---|---|---|---|---|---|
| 228 | ENSG000001 | ENSG000001 | ATG4B | chr2 | + | ENSG000001 | ENSG000001 | TRUE | cassette^1\|i | TRUE | C1_A | chr2:241668 | FALSE | TRUE | TRUE | 0.001 | 0.283 | 0.263 | 1 | 0 | cassette | 0.263 |
| 229 | ENSG000001 | ENSG000001 | LINGO1 | chr15 | - | ENSG000001 | ENSG000001 | TRUE | afe^1\|putati | TRUE | Proximal | chr15:77641 | FALSE | TRUE | TRUE | 0.002 | 0.222 | 0.203 | 1 | 0 | AFE/ALE | 0.203 |
| 230 | ENSG000001 | ENSG000001 | LINGO1 | chr15 | - | ENSG000001 | ENSG000001 | TRUE | cassette^3\|p | TRUE | C1_A | chr15:77756 | FALSE | FALSE | TRUE | 0.002 | 0.136 | 0.112 | 0.994 | 0.003 | cassette | 0.112 |
| 231 | ENSG000001 | ENSG000001 | ONECUT1 | chr15 | - | ENSG000001 | ENSG000001 | TRUE | putative_alt5 | TRUE | Distal | chr15:52777 | FALSE | FALSE | TRUE | 0 | 0.227 | 0.207 | 1 | 0 | AFE/ALE | 0.207 |
| 232 | ENSG000001 | ENSG000001 | ONECUT1 | chr15 | - | ENSG000001 | ENSG000001 | TRUE | putative_alt5^4\|ir^7\|ale^ | A_C2 | | chr15: | FALSE | FALSE | TRUE | 0.006 | 0.244 | 0.225 | 1 | 0 | cassette | 0.225 |
| 233 | ENSG000001 | ENSG000001 | DLGAP1 | chr18 | - | ENSG000001 | ENSG000001 | FALSE | putative_ale | TRUE | Proximal | chr18:41476 | FALSE | TRUE | TRUE | 0.004 | 0.214 | 0.185 | 0.998 | 0 | AFE/ALE | 0.185 |
| 234 | ENSG000001 | ENSG000001 | INSR | chr19 | - | ENSG000001 | ENSG000001 | TRUE | ir^2\|putative | TRUE | Proximal | chr19:71698 | FALSE | TRUE | TRUE | 0.004 | 0.432 | 0.407 | 1 | 0 | AFE/ALE | 0.407 |
| 235 | ENSG000001 | ENSG000001 | KNDC1 | chr10 | + | ENSG000001 | ENSG000001 | TRUE | ir^3\|putative | TRUE | Proximal | chr10:13320 | FALSE | TRUE | TRUE | 0.002 | 0.254 | 0.226 | 1 | 0 | AFE/ALE | 0.226 |
| 236 | ENSG000001 | ENSG000001 | TRAPPC12 | chr2 | + | ENSG000001 | ENSG000001 | TRUE | cassette^1\|i | TRUE | C1_A | chr2:345769 | FALSE | FALSE | TRUE | 0.001 | 0.201 | 0.174 | 0.998 | 0.001 | cassette | 0.174 |
| 237 | ENSG000001 | ENSG000001 | TRAPPC12 | chr2 | + | ENSG000001 | ENSG000001 | TRUE | cassette^1\|i | FALSE | Proximal | chr2:345769 | FALSE | FALSE | TRUE | 0.028 | 0.369 | 0.339 | 1 | 0 | AFE/ALE | 0.339 |
| 238 | ENSG000001 | ENSG000001 | B3GALT1 | chr2 | + | ENSG000001 | ENSG000001 | TRUE | alt5^1\|putat | TRUE | Proximal | chr2:167868 | FALSE | FALSE | TRUE | 0.005 | 0.146 | 0.123 | 0.991 | 0.004 | AFE/ALE | 0.123 |
| 239 | ENSG000001 | ENSG000001 | CRLF3 | chr17 | - | ENSG000001 | ENSG000001 | TRUE | ir^5\|putative | TRUE | Proximal | chr17:30786 | FALSE | FALSE | FALSE | 0.022 | 0.153 | 0.117 | 0.929 | 0.026 | AFE/ALE | 0.117 |
| 240 | ENSG000001 | ENSG000001 | RIMS2 | chr8 | + | ENSG000001 | ENSG000001 | TRUE | cassette^3\|a | FALSE | C2_A | chr8:104094 | FALSE | FALSE | TRUE | 0.035 | 0.18 | 0.144 | 0.98 | 0.007 | cassette | 0.144 |
| 241 | ENSG000001 | ENSG000001 | CHID1 | chr11 | - | ENSG000001 | ENSG000001 | FALSE | alt5^1 | FALSE | Proximal | chr11:90098 | FALSE | TRUE | TRUE | 0.014 | 0.458 | 0.449 | 1 | 0 | AFE/ALE | 0.449 |
| 242 | ENSG000001 | ENSG000001 | CDH4 | chr20 | + | ENSG000001 | ENSG000001 | TRUE | putative_ale | TRUE | Proximal | chr20:61254 | FALSE | TRUE | TRUE | 0.003 | 0.253 | 0.226 | 1 | 0 | AFE/ALE | 0.226 |
| 243 | ENSG000001 | ENSG000001 | EHMT1 | chr9 | + | ENSG000001 | ENSG000001 | TRUE | cassette^2\|i | FALSE | C1_A | chr9:137778 | FALSE | FALSE | TRUE | 0.004 | 0.153 | 0.127 | 0.993 | 0.003 | cassette | 0.127 |
| 244 | ENSG000001 | ENSG000001 | ADGRB1 | chr8 | + | ENSG000001 | ENSG000001 | TRUE | cassette^1\|a | TRUE | Proximal | chr8:142529 | FALSE | FALSE | FALSE | 0.005 | 0.122 | 0.096 | 0.93 | 0.037 | AFE/ALE | 0.096 |
| 245 | ENSG000001 | ENSG000001 | ADGRB1 | chr8 | + | ENSG000001 | ENSG000001 | TRUE | cassette^1\|a | TRUE | Proximal | chr8:142529 | FALSE | TRUE | TRUE | 0.005 | 0.259 | 0.237 | 1 | 0 | AFE/ALE | 0.237 |
| 246 | ENSG000001 | ENSG000001 | KCNIP1 | chr5 | + | ENSG000001 | ENSG000001 | TRUE | cassette^1\|c | FALSE | C2_A | chr5:170669 | FALSE | FALSE | TRUE | 0.003 | 0.204 | 0.172 | 0.997 | 0.001 | cassette | 0.172 |
| 247 | ENSG000001 | ENSG000001 | PLCXD1 | chrY | + | ENSG000001 | ENSG000001 | TRUE | ale^1\|afe^1 | TRUE | Proximal | chrY:281684 | FALSE | TRUE | TRUE | 0.005 | 0.262 | 0.238 | 1 | 0 | AFE/ALE | 0.238 |
| 248 | ENSG000001 | ENSG000001 | EP400 | chr12 | + | ENSG000001 | ENSG000001 | TRUE | cassette^1\|i | TRUE | C1_C2 | chr12:13199 | FALSE | TRUE | TRUE | 0.006 | 0.216 | -0.195 | 1 | 0 | skipping | 0.195 |
| 249 | ENSG000001 | ENSG000001 | PCBP3 | chr21 | + | ENSG000001 | ENSG000001 | TRUE | cassette^2\|i | TRUE | Proximal | chr21:45755 | FALSE | TRUE | TRUE | 0.027 | 0.609 | 0.581 | 1 | 0 | AFE/ALE | 0.581 |
| 250 | ENSG000001 | ENSG000001 | WDR27 | chr6 | - | ENSG000001 | ENSG000001 | TRUE | cassette^2 | FALSE | C2_C1 | chr6:169658 | FALSE | FALSE | TRUE | 0.045 | 0.22 | -0.17 | 0.99 | 0.003 | skipping | 0.17 |
| 251 | ENSG000001 | ENSG000001 | SLC24A3 | chr20 | + | ENSG000001 | ENSG000001 | TRUE | cassette^2\|a | TRUE | C1_A | chr20:19681 | FALSE | TRUE | TRUE | 0.029 | 0.356 | 0.309 | 1 | 0 | cassette | 0.309 |
| 252 | ENSG000001 | ENSG000001 | ARL15 | chr5 | - | ENSG000001 | ENSG000001 | TRUE | ale^2\|putati | TRUE | Distal | chr5:542853 | FALSE | FALSE | TRUE | 0.025 | 0.183 | 0.156 | 0.965 | 0.008 | AFE/ALE | 0.156 |
| 253 | ENSG000001 | ENSG000001 | ARL15 | chr5 | - | ENSG000001 | ENSG000001 | TRUE | ale^2\|putati | TRUE | Proximal | chr5:541719 | FALSE | FALSE | FALSE | 0.002 | 0.142 | 0.113 | 0.926 | 0.034 | AFE/ALE | 0.113 |
| 254 | ENSG000001 | ENSG000001 | ADARB2 | chr10 | - | ENSG000001 | ENSG000001 | TRUE | cassette^1\|r | TRUE | C1_A | chr10:13762 | FALSE | TRUE | TRUE | 0.002 | 0.61 | 0.603 | 1 | 0 | cassette | 0.603 |
| 255 | ENSG000001 | ENSG000001 | ADARB2 | chr10 | - | ENSG000001 | ENSG000001 | TRUE | alt3^7\|putat | TRUE | Proximal | chr10:16459 | FALSE | FALSE | TRUE | 0.015 | 0.208 | 0.186 | 1 | 0 | AFE/ALE | 0.186 |
| 256 | ENSG000001 | ENSG000001 | GNB1L | chr22 | - | ENSG000001 | ENSG000001 | TRUE | cassette^1\|a | TRUE | Proximal | chr22:19824 | FALSE | FALSE | TRUE | 0.003 | 0.538 | 0.523 | 1 | 0 | AFE/ALE | 0.523 |
| 257 | ENSG000001 | ENSG000001 | RASA3 | chr13 | - | ENSG000001 | ENSG000001 | TRUE | putative_ale | TRUE | Proximal | chr13:11399 | FALSE | TRUE | TRUE | 0.001 | 0.261 | 0.237 | 1 | 0 | AFE/ALE | 0.237 |
| 258 | ENSG000001 | ENSG000001 | ZNF529 | chr19 | - | ENSG000001 | ENSG000001 | TRUE | cassette^1\|a | TRUE | Proximal | chr19:36572 | FALSE | TRUE | TRUE | 0.011 | 0.39 | 0.362 | 1 | 0 | AFE/ALE | 0.362 |
| 259 | ENSG000001 | ENSG000001 | ZFP91 | chr11 | + | ENSG000001 | ENSG000001 | FALSE | cassette^1 | TRUE | C1_A | chr11:58616 | FALSE | FALSE | TRUE | 0.001 | 0.21 | 0.19 | 1 | 0 | cassette | 0.19 |
| 260 | ENSG000001 | ENSG000001 | ZSCAN30 | chr18 | - | ENSG000001 | ENSG000001 | TRUE | alt3^1\|ale^2 | TRUE | Distal | chr18:35254 | FALSE | FALSE | FALSE | 0.045 | 0.202 | 0.145 | 0.929 | 0.017 | AFE/ALE | 0.145 |
| 261 | ENSG000001 | ENSG000001 | MAPK12 | chr22 | - | ENSG000001 | ENSG000001 | FALSE | alt5^1 | TRUE | Distal | chr22:50248 | FALSE | FALSE | TRUE | 0.046 | 0.235 | 0.184 | 0.994 | 0.001 | AFE/ALE | 0.184 |
| 262 | ENSG000001 | ENSG000001 | AGRN | chr1 | + | ENSG000001 | ENSG000001 | TRUE | ale^1\|afe^1 | TRUE | Proximal | chr1:102246 | FALSE | TRUE | TRUE | 0.009 | 0.186 | 0.165 | 1 | 0 | AFE/ALE | 0.165 |
| 263 | ENSG000001 | ENSG000001 | RALGAPA2 | chr20 | - | ENSG000001 | ENSG000001 | FALSE | cassette^1 | TRUE | C1_A | chr20:20491 | FALSE | TRUE | TRUE | 0.004 | 0.416 | 0.398 | 1 | 0 | cassette | 0.398 |
| 264 | ENSG000001 | ENSG000001 | GREB1 | chr2 | + | ENSG000001 | ENSG000001 | TRUE | cassette^2\|a | TRUE | C1_A | chr2:115808 | FALSE | FALSE | TRUE | 0.011 | 0.18 | 0.159 | 1 | 0 | cassette | 0.159 |
| 265 | ENSG000001 | ENSG000001 | GREB1 | chr2 | + | ENSG000001 | ENSG000001 | TRUE | cassette^2\|a | TRUE | Proximal | chr2:114833 | FALSE | FALSE | FALSE | 0.012 | 0.117 | 0.096 | 0.927 | 0.04 | AFE/ALE | 0.096 |
| 266 | ENSG000001 | ENSG000001 | ELAVL3 | chr19 | - | ENSG000001 | ENSG000001 | FALSE | cassette^1 | TRUE | C1_A | chr19:11463 | FALSE | FALSE | TRUE | 0 | 0.141 | 0.115 | 0.993 | 0.003 | cassette | 0.115 |
| 267 | ENSG000001 | ENSG000001 | TRRAP | chr7 | + | ENSG000001 | ENSG000001 | TRUE | cassette^1\|i | TRUE | C1_A | chr7:988812 | FALSE | FALSE | TRUE | 0 | 0.149 | 0.126 | 1 | 0 | cassette | 0.126 |
| 268 | ENSG000001 | ENSG000001 | ARMCX4 | chrX | + | ENSG000001 | ENSG000001 | FALSE | ale^1 | FALSE | Distal | chrX:101488 | FALSE | FALSE | FALSE | 0.026 | 0.183 | 0.14 | 0.911 | 0.019 | AFE/ALE | 0.14 |
| 269 | ENSG000001 | ENSG000001 | NCOR2 | chr12 | - | ENSG000001 | ENSG000001 | TRUE | cassette^1\|p | TRUE | C1_A | chr12:12447 | FALSE | FALSE | TRUE | 0.001 | 0.129 | 0.106 | 0.966 | 0.017 | cassette | 0.106 |
| 270 | ENSG000001 | ENSG000001 | NCOR2 | chr12 | - | ENSG000001 | ENSG000001 | TRUE | cassette^1\|p | TRUE | Distal | chr12:12447 | FALSE | FALSE | TRUE | 0.005 | 0.169 | 0.147 | 1 | 0 | AFE/ALE | 0.147 |
| 271 | ENSG000001 | ENSG000001 | MYO18A | chr17 | - | ENSG000001 | ENSG000001 | TRUE | cassette^4\|a | TRUE | C1_A | chr17:29131 | FALSE | FALSE | TRUE | 0.001 | 0.371 | 0.361 | 1 | 0 | cassette | 0.361 |
| 272 | ENSG000001 | ENSG000001 | MYO18A | chr17 | - | ENSG000001 | ENSG000001 | TRUE | cassette^4\|a | TRUE | Distal | chr17:29124 | FALSE | FALSE | TRUE | 0.004 | 0.536 | 0.528 | 1 | 0 | AFE/ALE | 0.528 |
| 273 | ENSG000001 | ENSG000001 | NF1 | chr17 | + | ENSG000001 | ENSG000001 | FALSE | cassette^1 | FALSE | C1_A | chr17:31249 | FALSE | FALSE | FALSE | 0.026 | 0.124 | 0.096 | 0.925 | 0.04 | cassette | 0.096 |
| 274 | ENSG000001 | ENSG000001 | DAPK1 | chr9 | + | ENSG000001 | ENSG000001 | TRUE | cassette^1\|a | TRUE | C1_A | chr9:874991 | FALSE | FALSE | TRUE | 0.001 | 0.147 | 0.121 | 0.998 | 0.001 | cassette | 0.121 |
| 275 | ENSG000001 | ENSG000001 | DAPK1 | chr9 | + | ENSG000001 | ENSG000001 | TRUE | cassette^1\|a | FALSE | Proximal | chr9:874991 | FALSE | FALSE | TRUE | 0.046 | 0.222 | 0.176 | 1 | 0 | AFE/ALE | 0.176 |
| 276 | ENSG000001 | ENSG000001 | RGPD4 | chr2 | + | ENSG000001 | ENSG000001 | FALSE | putative_afe | TRUE | Proximal | chr2:107878 | FALSE | TRUE | TRUE | 0.007 | 0.274 | 0.254 | 1 | 0 | AFE/ALE | 0.254 |
| 277 | ENSG000001 | ENSG000001 | ZGPAT | chr20 | + | ENSG000001 | ENSG000001 | FALSE | cassette^1 | TRUE | C1_A | chr20:63709 | FALSE | TRUE | TRUE | 0.005 | 0.433 | 0.414 | 1 | 0 | cassette | 0.414 |
| 278 | ENSG000001 | ENSG000001 | PHF2 | chr9 | + | ENSG000001 | ENSG000001 | TRUE | cassette^2\|t | TRUE | Proximal | chr9:936605 | FALSE | TRUE | TRUE | 0 | 0.495 | 0.481 | 1 | 0 | AFE/ALE | 0.481 |
| 279 | ENSG000001 | ENSG000001 | MYO1C | chr17 | - | ENSG000001 | ENSG000001 | FALSE | cassette^1 | TRUE | C1_C2 | chr17:14722 | FALSE | TRUE | TRUE | 0.006 | 0.234 | -0.204 | 1 | 0 | skipping | 0.204 |
| 280 | ENSG000001 | ENSG000001 | SDAD1 | chr4 | - | ENSG000001 | ENSG000001 | TRUE | cassette^1\|a | TRUE | C1_A | chr4:759848 | FALSE | FALSE | TRUE | 0.001 | 0.15 | 0.123 | 0.99 | 0.004 | cassette | 0.123 |
| 281 | ENSG000001 | ENSG000001 | UVRAG | chr11 | + | ENSG000001 | ENSG000001 | TRUE | cassette^1\|i | FALSE | C1_A | chr11:76008 | FALSE | FALSE | TRUE | 0.031 | 0.181 | 0.139 | 0.951 | 0.014 | cassette | 0.139 |
| 282 | ENSG000001 | ENSG000001 | RYR2 | chr1 | + | ENSG000001 | ENSG000001 | TRUE | alt3^1\|ale^2 | TRUE | Distal | chr1:237585 | FALSE | FALSE | TRUE | 0.012 | 0.294 | 0.246 | 0.986 | 0.002 | AFE/ALE | 0.246 |
| 283 | ENSG000001 | ENSG000001 | CEP290 | chr12 | - | ENSG000001 | ENSG000001 | FALSE | alt3^1 | TRUE | Proximal | chr12:88086 | FALSE | TRUE | TRUE | 0.004 | 0.469 | 0.455 | 1 | 0 | AFE/ALE | 0.455 |
| 284 | ENSG000001 | ENSG000001 | CEP290 | chr12 | - | ENSG000001 | ENSG000001 | FALSE | cassette^1 | FALSE | C2_C1 | chr12:88111 | FALSE | FALSE | FALSE | 0.046 | 0.185 | -0.128 | 0.922 | 0.027 | skipping | 0.128 |
| 285 | ENSG000001 | ENSG000001 | UNC13B | chr9 | + | ENSG000001 | ENSG000001 | FALSE | cassette^1 | FALSE | C1_A | chr9:353139 | FALSE | TRUE | TRUE | 0.044 | 0.419 | 0.377 | 1 | 0 | cassette | 0.377 |
| 286 | ENSG000001 | ENSG000001 | COLGALT2 | chr1 | - | ENSG000001 | ENSG000001 | FALSE | alt5^1 | TRUE | Distal | chr1:183978 | FALSE | FALSE | FALSE | 0.02 | 0.152 | 0.12 | 0.939 | 0.024 | AFE/ALE | 0.12 |
| 287 | ENSG000001 | ENSG000001 | SCAMP5 | chr15 | + | ENSG000001 | ENSG000001 | TRUE | cassette^7\|a | TRUE | C1_A | chr15:75007 | FALSE | FALSE | TRUE | 0.036 | 0.354 | 0.323 | 1 | 0 | cassette | 0.323 |
| 288 | ENSG000001 | ENSG000001 | SCAMP5 | chr15 | + | ENSG000001 | ENSG000001 | TRUE | cassette^7\|a | FALSE | C2_A | chr15:75009 | FALSE | FALSE | TRUE | 0.047 | 0.203 | 0.167 | 1 | 0 | cassette | 0.167 |
| 289 | ENSG000001 | ENSG000001 | ZNF521 | chr18 | - | ENSG000001 | ENSG000001 | TRUE | putative_ale | TRUE | Proximal | chr18:25160 | FALSE | FALSE | TRUE | 0.001 | 0.162 | 0.138 | 0.996 | 0.002 | AFE/ALE | 0.138 |
| 290 | ENSG000001 | ENSG000001 | ZNF521 | chr18 | - | ENSG000001 | ENSG000001 | TRUE | putative_ale | TRUE | Proximal | chr18:25162 | FALSE | FALSE | TRUE | 0.008 | 0.2 | 0.177 | 1 | 0 | AFE/ALE | 0.177 |
| 291 | ENSG000001 | ENSG000001 | KIAA0753 | chr17 | - | ENSG000001 | ENSG000001 | FALSE | cassette^1 | TRUE | C1_A | chr17:65809 | FALSE | TRUE | TRUE | 0.004 | 0.483 | 0.461 | 1 | 0 | cassette | 0.461 |
| 292 | ENSG000002 | ENSG000002 | LINC00963 | chr9 | + | ENSG000002 | ENSG000002 | TRUE | cassette^2\|a | FALSE | A_C2 | chr9:129503 | FALSE | FALSE | TRUE | 0.014 | 0.144 | 0.117 | 0.98 | 0.009 | cassette | 0.117 |
| 293 | ENSG000002 | ENSG000002 | STARD7-AS1 | chr2 | + | ENSG000002 | ENSG000002 | TRUE | cassette^1\|a | FALSE | Distal | chr2:962089 | FALSE | FALSE | TRUE | 0.042 | 0.253 | 0.193 | 0.958 | 0.006 | AFE/ALE | 0.193 |
| 294 | ENSG000002 | ENSG000002 | PCDHA2 | chr5 | + | ENSG000002 | ENSG000002 | TRUE | alt3^1\|alt3_ | TRUE | E1_E2_J2 | chr5:140796 | FALSE | FALSE | TRUE | 0.003 | 0.156 | 0.134 | 1 | 0 | cassette | 0.134 |
| 295 | ENSG000002 | ENSG000002 | LIN52 | chr14 | + | ENSG000002 | ENSG000002 | FALSE | ale^1 | FALSE | Proximal | chr14:74101 | FALSE | FALSE | TRUE | 0.018 | 0.181 | 0.152 | 0.988 | 0.004 | AFE/ALE | 0.152 |
| 296 | ENSG000002 | ENSG000002 | NYNRIN | chr14 | + | ENSG000002 | ENSG000002 | TRUE | cassette^1\|i | TRUE | C1_A | chr14:24411 | FALSE | FALSE | FALSE | 0.009 | 0.151 | 0.122 | 0.946 | 0.021 | cassette | 0.122 |
| 297 | ENSG000002 | ENSG000002 | ZDHHC11B | chr5 | - | ENSG000002 | ENSG000002 | TRUE | cassette^1\|i | TRUE | Proximal | chr5:733156 | FALSE | FALSE | TRUE | 0.002 | 0.32 | 0.299 | 1 | 0 | AFE/ALE | 0.299 |
| 298 | ENSG000002 | ENSG000002 | ANKRD28 | chr3 | - | ENSG000002 | ENSG000002 | TRUE | cassette^3\|a | TRUE | E2_E1_J2 | chr3:156673 | FALSE | FALSE | TRUE | 0 | 0.296 | 0.27 | 1 | 0 | cassette | 0.27 |
| 299 | ENSG000002 | ENSG000002 | ANKRD28 | chr3 | - | ENSG000002 | ENSG000002 | TRUE | cassette^3\|a | TRUE | E2_E1_J1 | chr3:156715 | FALSE | FALSE | TRUE | 0 | 0.174 | 0.152 | 0.997 | 0.001 | cassette | 0.152 |
| 300 | ENSG000002 | ENSG000002 | ANKRD28 | chr3 | - | ENSG000002 | ENSG000002 | TRUE | cassette^3\|a | TRUE | E2_E1_J1 | chr3:156764 | FALSE | FALSE | TRUE | 0.009 | 0.266 | 0.24 | 0.975 | 0.004 | cassette | 0.24 |
| 301 | ENSG000002 | ENSG000002 | ANKRD28 | chr3 | - | ENSG000002 | ENSG000002 | TRUE | cassette^3\|a | FALSE | A_C2 | chr3:156770 | FALSE | FALSE | TRUE | 0.001 | 0.219 | 0.192 | 0.991 | 0.002 | cassette | 0.192 |
| 302 | ENSG000002 | ENSG000002 | ANKRD28 | chr3 | - | ENSG000002 | ENSG000002 | TRUE | cassette^3\|a | TRUE | E2_E1_J1 | chr3:157131 | FALSE | FALSE | TRUE | 0 | 0.304 | 0.284 | 1 | 0 | cassette | 0.284 |
| 303 | ENSG000002 | ENSG000002 | ANKRD28 | chr3 | - | ENSG000002 | ENSG000002 | TRUE | cassette^3\|a | TRUE | Distal | chr3:157062 | FALSE | TRUE | TRUE | 0.005 | 0.438 | 0.393 | 1 | 0 | AFE/ALE | 0.393 |

| | | | | | | | | | | | | | | | | | | | | | | |
|---|---|---|---|---|---|---|---|---|---|---|---|---|---|---|---|---|---|---|---|---|---|---|
| 304 | ENSG000002 | ENSG000002 | DENND1B | chr1 | - | ENSG000002 | ENSG000002 | FALSE | putative_ale | TRUE | Proximal | chr1:197560 | FALSE | FALSE | FALSE | 0.023 | 0.159 | 0.129 | 0.944 | 0.019 | AFE/ALE | 0.129 |
| 305 | ENSG000002 | ENSG000002 | RPS29 | chr14 | - | ENSG000002 | ENSG000002 | TRUE | alt3^162\|ale | TRUE | Proximal | chr14:49583 | FALSE | FALSE | FALSE | 0 | 0.151 | 0.122 | 0.923 | 0.026 | AFE/ALE | 0.122 |
| 306 | ENSG000002 | ENSG000002 | RPS29 | chr14 | - | ENSG000002 | ENSG000002 | TRUE | alt3^162\|ale | TRUE | Proximal | chr14:49585 | FALSE | FALSE | FALSE | 0 | 0.136 | 0.11 | 0.944 | 0.027 | AFE/ALE | 0.11 |
| 307 | ENSG000002 | ENSG000002 | RPS29 | chr14 | - | ENSG000002 | ENSG000002 | TRUE | alt3^162\|ale | TRUE | Distal | chr14:49573 | FALSE | FALSE | TRUE | 0 | 0.657 | 0.646 | 1 | 0 | AFE/ALE | 0.646 |
| 308 | ENSG000002 | ENSG000002 | RPS29 | chr14 | - | ENSG000002 | ENSG000002 | TRUE | alt3^162\|ale | TRUE | Distal | chr14:49585 | FALSE | FALSE | TRUE | 0 | 0.212 | 0.189 | 1 | 0 | AFE/ALE | 0.189 |
| 309 | ENSG000002 | ENSG000002 | SEPTIN7P2 | chr7 | - | ENSG000002 | ENSG000002 | TRUE | cassette^1\|a | TRUE | C1_A | chr7:457356 | FALSE | FALSE | TRUE | 0.029 | 0.356 | 0.328 | 1 | 0 | cassette | 0.328 |
| 310 | ENSG000002 | ENSG000002 | SEPTIN7P2 | chr7 | - | ENSG000002 | ENSG000002 | TRUE | cassette^1\|a | TRUE | Distal | chr7:457283 | FALSE | FALSE | TRUE | 0.006 | 0.281 | 0.262 | 1 | 0 | AFE/ALE | 0.262 |
| 311 | ENSG000002 | ENSG000002 | NPEPL1 | chr20 | + | ENSG000002 | ENSG000002 | TRUE | ir^2\|putative | TRUE | Proximal | chr20:58694 | FALSE | TRUE | TRUE | 0.001 | 0.247 | 0.22 | 1 | 0 | AFE/ALE | 0.22 |
| 312 | ENSG000002 | ENSG000002 | LINC00863 | chr10 | + | ENSG000002 | ENSG000002 | TRUE | cassette^1\|i | FALSE | A_C2 | chr10:87354 | FALSE | FALSE | TRUE | 0.018 | 0.212 | 0.173 | 0.963 | 0.006 | cassette | 0.173 |
| 313 | ENSG000002 | ENSG000002 | FAM66C | chr12 | + | ENSG000002 | ENSG000002 | TRUE | cassette^3\|a | TRUE | C1_A | chr12:81885 | FALSE | TRUE | TRUE | 0.015 | 0.447 | 0.436 | 1 | 0 | cassette | 0.436 |
| 314 | ENSG000002 | ENSG000002 | FAM66C | chr12 | + | ENSG000002 | ENSG000002 | TRUE | cassette^3\|a | FALSE | C1_A | chr12:81886 | FALSE | TRUE | TRUE | 0.034 | 0.383 | 0.357 | 1 | 0 | cassette | 0.357 |
| 315 | ENSG000002 | ENSG000002 | CROCCP4 | chr1 | + | ENSG000002 | ENSG000002 | TRUE | alt3^1\|putat | TRUE | Distal | chr1:167406 | FALSE | FALSE | TRUE | 0.001 | 0.21 | 0.186 | 1 | 0 | AFE/ALE | 0.186 |
| 316 | ENSG000002 | ENSG000002 | ZNF826P | chr19 | - | ENSG000002 | ENSG000002 | TRUE | cassette^2\|a | FALSE | C1_A | chr19:20409 | FALSE | FALSE | TRUE | 0.045 | 0.642 | 0.611 | 1 | 0 | cassette | 0.611 |
| 317 | ENSG000002 | ENSG000002 | ZNF826P | chr19 | - | ENSG000002 | ENSG000002 | TRUE | cassette^2\|a | FALSE | Proximal | chr19:20409 | FALSE | FALSE | TRUE | 0.023 | 0.522 | 0.508 | 1 | 0 | AFE/ALE | 0.508 |
| 318 | ENSG000002 | ENSG000002 | LINC01122 | chr2 | + | ENSG000002 | ENSG000002 | TRUE | cassette^2\|a | FALSE | Distal | chr2:586567 | FALSE | FALSE | FALSE | 0.028 | 0.184 | 0.147 | 0.947 | 0.014 | AFE/ALE | 0.147 |
| 319 | ENSG000002 | ENSG000002 | RPL23AP7 | chr2 | - | ENSG000002 | ENSG000002 | TRUE | alt5^3\|ir^3 | TRUE | Proximal | chr2:113625 | FALSE | FALSE | TRUE | 0.038 | 0.241 | 0.197 | 0.999 | 0 | AFE/ALE | 0.197 |
| 320 | ENSG000002 | ENSG000002 | PEDS1 | chr20 | - | ENSG000002 | ENSG000002 | FALSE | cassette^1 | FALSE | C1_C2 | chr20:50129 | FALSE | FALSE | TRUE | 0.033 | 0.262 | -0.23 | 1 | 0 | skipping | 0.23 |
| 321 | ENSG000002 | ENSG000002 | ZNF286B | chr17 | - | ENSG000002 | ENSG000002 | TRUE | cassette^1\|i | TRUE | C1_C2 | chr17:18680 | FALSE | FALSE | FALSE | 0.027 | 0.14 | -0.11 | 0.921 | 0.034 | skipping | 0.11 |
| 322 | ENSG000002 | ENSG000002 | LINC02506 | chr4 | + | ENSG000002 | ENSG000002 | TRUE | cassette^6\|i | TRUE | C2_A | chr4:321571 | FALSE | FALSE | TRUE | 0.031 | 0.146 | 0.121 | 0.987 | 0.006 | cassette | 0.121 |
| 323 | ENSG000002 | ENSG000002 | ENSG000002 | chr9 | + | ENSG000002 | ENSG000002 | FALSE | cassette^1 | TRUE | C1_A | chr9:128952 | FALSE | TRUE | TRUE | 0.003 | 0.783 | 0.781 | 1 | 0 | cassette | 0.781 |
| 324 | ENSG000002 | ENSG000002 | STX16-NPEPL | chr20 | + | ENSG000002 | ENSG000002 | TRUE | ir^2\|putative | TRUE | Proximal | chr20:58694 | FALSE | TRUE | TRUE | 0.001 | 0.248 | 0.224 | 1 | 0 | AFE/ALE | 0.224 |
| 325 | ENSG000002 | ENSG000002 | FAM66D | chr8 | + | ENSG000002 | ENSG000002 | TRUE | cassette^1\|a | TRUE | C1_A | chr8:121220 | FALSE | FALSE | TRUE | 0.021 | 0.594 | 0.584 | 1 | 0 | cassette | 0.584 |
| 326 | ENSG000002 | ENSG000002 | ZFP91-CNTF | chr11 | + | ENSG000002 | ENSG000002 | FALSE | cassette^1 | TRUE | C1_A | chr11:58616 | FALSE | FALSE | TRUE | 0.001 | 0.21 | 0.19 | 1 | 0 | cassette | 0.19 |
| 327 | ENSG000002 | ENSG000002 | ENSG000002 | chr15 | + | ENSG000002 | ENSG000002 | TRUE | orphan_junc | TRUE | Orphan | chr15:67840 | FALSE | FALSE | TRUE | 0.001 | 0.209 | 0.181 | 0.994 | 0.002 | cassette | 0.181 |
| 328 | ENSG000002 | ENSG000002 | ENSG000002 | chr15 | + | ENSG000002 | ENSG000002 | TRUE | orphan_junc | TRUE | Orphan | chr15:67840 | FALSE | FALSE | TRUE | 0.02 | 0.198 | 0.173 | 0.987 | 0.003 | cassette | 0.173 |
| 329 | ENSG000002 | ENSG000002 | CORO7 | chr16 | - | ENSG000002 | ENSG000002 | FALSE | putative_ale | TRUE | Proximal | chr16:43674 | FALSE | TRUE | TRUE | 0.019 | 0.347 | 0.323 | 1 | 0 | AFE/ALE | 0.323 |
| 330 | ENSG000002 | ENSG000002 | SUZ12P1 | chr17 | + | ENSG000002 | ENSG000002 | TRUE | putative_alt3 | TRUE | Proximal | chr17:30786 | FALSE | FALSE | FALSE | 0.021 | 0.146 | 0.112 | 0.922 | 0.031 | AFE/ALE | 0.112 |
| 331 | ENSG000002 | ENSG000002 | ENSG000002 | chr19 | - | ENSG000002 | ENSG000002 | TRUE | alt5^3 | TRUE | Distal | chr19:43900 | FALSE | FALSE | TRUE | 0.047 | 0.219 | 0.167 | 0.962 | 0.008 | AFE/ALE | 0.167 |
| 332 | ENSG000002 | ENSG000002 | ENSG000002 | chr20 | + | ENSG000002 | ENSG000002 | FALSE | cassette^1 | TRUE | C1_A | chr20:63709 | FALSE | TRUE | TRUE | 0.005 | 0.433 | 0.414 | 1 | 0 | cassette | 0.414 |
| 333 | ENSG000002 | ENSG000002 | SEC22B4P | chr1 | - | ENSG000002 | ENSG000002 | TRUE | alt5^2\|alt3_ | TRUE | E1_E2_J1 | chr1:146376 | FALSE | FALSE | TRUE | 0.002 | 0.165 | 0.145 | 0.981 | 0.007 | cassette | 0.145 |
| 334 | ENSG000002 | ENSG000002 | SEC22B4P | chr1 | - | ENSG000002 | ENSG000002 | TRUE | alt5^2\|alt3_ | TRUE | E1_E2_J2 | chr1:146376 | FALSE | FALSE | FALSE | 0.002 | 0.136 | 0.114 | 0.94 | 0.028 | cassette | 0.114 |
| 335 | ENSG000002 | ENSG000002 | ENSG000002 | chr21 | + | ENSG000002 | ENSG000002 | TRUE | cassette^127\|alt3^84\|alt | | C1_A | chr21: | FALSE | FALSE | TRUE | 0.048 | 0.295 | 0.248 | 1 | 0 | cassette | 0.248 |
| 336 | ENSG000002 | ENSG000002 | ENSG000002 | chr21 | + | ENSG000002 | ENSG000002 | TRUE | cassette^127\|alt3^84\|alt | | C1_A | chr21: | FALSE | FALSE | FALSE | 0 | 0.125 | 0.101 | 0.928 | 0.037 | cassette | 0.101 |
| 337 | ENSG000002 | ENSG000002 | ENSG000002 | chr21 | + | ENSG000002 | ENSG000002 | TRUE | cassette^127\|alt3^84\|alt | | C1_A | chr21: | FALSE | FALSE | TRUE | 0 | 0.161 | 0.134 | 0.983 | 0.007 | cassette | 0.134 |
| 338 | ENSG000002 | ENSG000002 | ENSG000002 | chr21 | + | ENSG000002 | ENSG000002 | TRUE | cassette^127\|alt3^84\|alt | | C1_A | chr21: | FALSE | TRUE | TRUE | 0.004 | 0.676 | 0.667 | 1 | 0 | cassette | 0.667 |
| 339 | ENSG000002 | ENSG000002 | ENSG000002 | chr21 | + | ENSG000002 | ENSG000002 | TRUE | cassette^127\|alt3^84\|alt | | C1_A | chr21: | FALSE | FALSE | TRUE | 0.001 | 0.174 | 0.149 | 0.976 | 0.008 | cassette | 0.149 |
| 340 | ENSG000002 | ENSG000002 | ENSG000002 | chr21 | + | ENSG000002 | ENSG000002 | TRUE | cassette^127\|alt3^84\|alt | | C1_A | chr21: | FALSE | FALSE | TRUE | 0.004 | 0.166 | 0.144 | 0.975 | 0.009 | cassette | 0.144 |
| 341 | ENSG000002 | ENSG000002 | ENSG000002 | chr21 | + | ENSG000002 | ENSG000002 | TRUE | cassette^127\|alt3^84\|alt | | C1_A | chr21: | FALSE | FALSE | TRUE | 0.006 | 0.336 | 0.317 | 1 | 0 | cassette | 0.317 |
| 342 | ENSG000002 | ENSG000002 | ENSG000002 | chr21 | + | ENSG000002 | ENSG000002 | TRUE | cassette^127\|alt3^84\|alt | | C1_A | chr21: | FALSE | FALSE | TRUE | 0 | 0.462 | 0.464 | 1 | 0 | cassette | 0.464 |
| 343 | ENSG000002 | ENSG000002 | ENSG000002 | chr21 | + | ENSG000002 | ENSG000002 | TRUE | cassette^12 | TRUE | E1_E2_J1 | chr21:82168 | FALSE | FALSE | TRUE | 0.03 | 0.207 | 0.184 | 1 | 0 | cassette | 0.184 |
| 344 | ENSG000002 | ENSG000002 | ENSG000002 | chr21 | + | ENSG000002 | ENSG000002 | TRUE | cassette^127\|alt3^84\|alt | | C1_A | chr21: | FALSE | FALSE | TRUE | 0.001 | 0.196 | 0.173 | 0.981 | 0.005 | cassette | 0.173 |
| 345 | ENSG000002 | ENSG000002 | ENSG000002 | chr21 | + | ENSG000002 | ENSG000002 | TRUE | cassette^12 | TRUE | Distal | chr21:82058 | FALSE | FALSE | TRUE | 0.001 | 0.204 | 0.178 | 0.989 | 0.003 | AFE/ALE | 0.178 |
| 346 | ENSG000002 | ENSG000002 | ENSG000002 | chr21 | + | ENSG000002 | ENSG000002 | TRUE | cassette^12 | TRUE | C1_A1 | chr21:82072 | FALSE | FALSE | TRUE | 0.015 | 0.255 | 0.23 | 1 | 0 | cassette | 0.23 |
| 347 | ENSG000002 | ENSG000002 | ENSG000002 | chr21 | + | ENSG000002 | ENSG000002 | TRUE | cassette^12 | TRUE | Distal | chr21:82078 | FALSE | FALSE | FALSE | 0 | 0.157 | 0.13 | 0.94 | 0.024 | AFE/ALE | 0.13 |
| 348 | ENSG000002 | ENSG000002 | ENSG000002 | chr21 | + | ENSG000002 | ENSG000002 | TRUE | cassette^12 | TRUE | Proximal | chr21:82086 | FALSE | FALSE | TRUE | 0.003 | 0.324 | 0.301 | 1 | 0 | AFE/ALE | 0.301 |
| 349 | ENSG000002 | ENSG000002 | ENSG000002 | chr21 | + | ENSG000002 | ENSG000002 | TRUE | cassette^12 | TRUE | Distal | chr21:82086 | FALSE | FALSE | TRUE | 0.003 | 0.304 | 0.283 | 1 | 0 | AFE/ALE | 0.283 |
| 350 | ENSG000002 | ENSG000002 | ENSG000002 | chr21 | + | ENSG000002 | ENSG000002 | TRUE | cassette^12 | TRUE | C1_A1 | chr21:82090 | FALSE | FALSE | TRUE | 0.003 | 0.24 | 0.207 | 0.977 | 0.004 | cassette | 0.207 |
| 351 | ENSG000002 | ENSG000002 | ENSG000002 | chr21 | + | ENSG000002 | ENSG000002 | TRUE | cassette^12 | TRUE | Distal | chr21:82090 | FALSE | FALSE | TRUE | 0.003 | 0.201 | 0.173 | 0.967 | 0.008 | AFE/ALE | 0.173 |
| 352 | ENSG000002 | ENSG000002 | ENSG000002 | chr21 | + | ENSG000002 | ENSG000002 | TRUE | cassette^12 | TRUE | Proximal | chr21:82086 | FALSE | FALSE | TRUE | 0.001 | 0.15 | 0.126 | 0.956 | 0.018 | AFE/ALE | 0.126 |
| 353 | ENSG000002 | ENSG000002 | ENSG000002 | chr21 | + | ENSG000002 | ENSG000002 | TRUE | cassette^12 | TRUE | A_C2 | chr21:82005 | FALSE | FALSE | FALSE | 0.011 | 0.147 | 0.124 | 0.933 | 0.027 | cassette | 0.124 |
| 354 | ENSG000002 | ENSG000002 | ENSG000002 | chr21 | + | ENSG000002 | ENSG000002 | TRUE | cassette^12 | TRUE | Proximal | chr21:82011 | FALSE | FALSE | TRUE | 0.039 | 0.201 | 0.158 | 0.955 | 0.014 | AFE/ALE | 0.158 |
| 355 | ENSG000002 | ENSG000002 | ENSG000002 | chr21 | + | ENSG000002 | ENSG000002 | TRUE | cassette^12 | TRUE | Proximal | chr21:82072 | FALSE | FALSE | TRUE | 0.019 | 0.206 | 0.185 | 1 | 0 | AFE/ALE | 0.185 |
| 356 | ENSG000002 | ENSG000002 | ENSG000002 | chr21 | + | ENSG000002 | ENSG000002 | TRUE | cassette^12 | TRUE | Distal | chr21:82101 | FALSE | FALSE | TRUE | 0.002 | 0.432 | 0.419 | 1 | 0 | AFE/ALE | 0.419 |
| 357 | ENSG000002 | ENSG000002 | ENSG000002 | chr21 | + | ENSG000002 | ENSG000002 | TRUE | cassette^127\|alt3^84\|alt | | A_C2 | chr21: | FALSE | FALSE | FALSE | 0.021 | 0.152 | 0.121 | 0.915 | 0.031 | cassette | 0.121 |
| 358 | ENSG000002 | ENSG000002 | ENSG000002 | chr21 | + | ENSG000002 | ENSG000002 | TRUE | cassette^12 | TRUE | E2_E1_J1 | chr21:82120 | FALSE | FALSE | TRUE | 0 | 0.165 | 0.15 | 0.961 | 0.013 | cassette | 0.15 |
| 359 | ENSG000002 | ENSG000002 | ENSG000002 | chr21 | + | ENSG000002 | ENSG000002 | TRUE | cassette^12 | TRUE | Distal | chr21:82039 | FALSE | FALSE | FALSE | 0.029 | 0.169 | 0.149 | 0.908 | 0.021 | AFE/ALE | 0.149 |
| 360 | ENSG000002 | ENSG000002 | ENSG000002 | chr21 | + | ENSG000002 | ENSG000002 | TRUE | cassette^127\|alt3^84\|alt | | A_C2 | chr21: | FALSE | FALSE | TRUE | 0.01 | 0.183 | 0.148 | 0.972 | 0.008 | cassette | 0.148 |
| 361 | ENSG000002 | ENSG000002 | ENSG000002 | chr21 | + | ENSG000002 | ENSG000002 | TRUE | cassette^12 | TRUE | Distal | chr21:82069 | FALSE | FALSE | TRUE | 0 | 0.186 | 0.162 | 0.998 | 0.001 | AFE/ALE | 0.162 |
| 362 | ENSG000002 | ENSG000002 | ENSG000002 | chr21 | + | ENSG000002 | ENSG000002 | TRUE | cassette^12 | TRUE | C2_A | chr21:82126 | FALSE | FALSE | TRUE | 0.005 | 0.46 | 0.441 | 1 | 0 | cassette | 0.441 |
| 363 | ENSG000002 | ENSG000002 | ENSG000002 | chr21 | + | ENSG000002 | ENSG000002 | TRUE | cassette^12 | TRUE | C2_A2 | chr21:82108 | FALSE | FALSE | TRUE | 0 | 0.502 | 0.515 | 1 | 0 | cassette | 0.515 |
| 364 | ENSG000002 | ENSG000002 | ENSG000002 | chr21 | + | ENSG000002 | ENSG000002 | TRUE | cassette^127\|alt3^84\|alt | | A_C2 | chr21: | FALSE | FALSE | TRUE | 0.005 | 0.46 | 0.441 | 1 | 0 | cassette | 0.441 |
| 365 | ENSG000002 | ENSG000002 | ENSG000002 | chr21 | + | ENSG000002 | ENSG000002 | TRUE | cassette^12 | TRUE | Proximal | chr21:82058 | FALSE | FALSE | TRUE | 0.003 | 0.422 | 0.383 | 1 | 0 | AFE/ALE | 0.383 |
| 366 | ENSG000002 | ENSG000002 | ENSG000002 | chr21 | + | ENSG000002 | ENSG000002 | TRUE | cassette^127\|alt3^84\|alt | | A_C2 | chr21: | FALSE | FALSE | FALSE | 0.012 | 0.14 | 0.115 | 0.91 | 0.04 | cassette | 0.115 |
| 367 | ENSG000002 | ENSG000002 | ENSG000002 | chr21 | + | ENSG000002 | ENSG000002 | TRUE | cassette^12 | TRUE | C2_A1 | chr21:82154 | FALSE | FALSE | FALSE | 0.002 | 0.124 | 0.106 | 0.908 | 0.046 | cassette | 0.106 |
| 368 | ENSG000002 | ENSG000002 | ENSG000002 | chr21 | + | ENSG000002 | ENSG000002 | TRUE | cassette^127\|alt3^84\|alt | | A_C2 | chr21: | FALSE | FALSE | TRUE | 0 | 0.258 | 0.228 | 1 | 0 | cassette | 0.228 |
| 369 | ENSG000002 | ENSG000002 | ENSG000002 | chr21 | + | ENSG000002 | ENSG000002 | TRUE | cassette^127\|alt3^84\|alt | | A_C2 | chr21: | FALSE | FALSE | TRUE | 0 | 0.223 | 0.206 | 1 | 0 | cassette | 0.206 |
| 370 | ENSG000002 | ENSG000002 | ENSG000002 | chr21 | + | ENSG000002 | ENSG000002 | TRUE | cassette^12 | TRUE | C2_A | chr21:82146 | FALSE | FALSE | TRUE | 0.021 | 0.417 | 0.383 | 1 | 0 | cassette | 0.383 |
| 371 | ENSG000002 | ENSG000002 | ENSG000002 | chr21 | + | ENSG000002 | ENSG000002 | TRUE | cassette^127\|alt3^84\|alt | | A_C2 | chr21: | FALSE | FALSE | TRUE | 0 | 0.249 | 0.223 | 1 | 0 | cassette | 0.223 |
| 372 | ENSG000002 | ENSG000002 | ENSG000002 | chr21 | + | ENSG000002 | ENSG000002 | TRUE | cassette^127\|alt3^84\|alt | | A_C2 | chr21: | FALSE | FALSE | FALSE | 0.04 | 0.156 | 0.119 | 0.912 | 0.032 | cassette | 0.119 |
| 373 | ENSG000002 | ENSG000002 | ENSG000002 | chr21 | + | ENSG000002 | ENSG000002 | TRUE | cassette^12 | TRUE | E2_E1_J2 | chr21:82127 | FALSE | FALSE | TRUE | 0 | 0.22 | 0.193 | 1 | 0 | cassette | 0.193 |
| 374 | ENSG000002 | ENSG000002 | ENSG000002 | chr21 | + | ENSG000002 | ENSG000002 | TRUE | cassette^12 | TRUE | Distal | chr21:82072 | FALSE | FALSE | TRUE | 0.001 | 0.428 | 0.421 | 1 | 0 | AFE/ALE | 0.421 |
| 375 | ENSG000002 | ENSG000002 | ENSG000002 | chr21 | + | ENSG000002 | ENSG000002 | TRUE | cassette^127\|alt3^84\|alt | | A_C2 | chr21: | FALSE | FALSE | TRUE | 0.029 | 0.255 | 0.227 | 1 | 0 | cassette | 0.227 |
| 376 | ENSG000002 | ENSG000002 | ENSG000002 | chr21 | + | ENSG000002 | ENSG000002 | TRUE | cassette^127\|alt3^84\|alt | | A_C2 | chr21: | FALSE | FALSE | TRUE | 0.001 | 0.141 | 0.117 | 0.993 | 0.003 | cassette | 0.117 |
| 377 | ENSG000002 | ENSG000002 | ENSG000002 | chr21 | - | ENSG000002 | ENSG000002 | TRUE | cassette^1\|i | TRUE | A_C2 | chr21:51188 | FALSE | FALSE | TRUE | 0.002 | 0.245 | 0.221 | 1 | 0 | cassette | 0.221 |
| 378 | ENSG000002 | ENSG000002 | ENSG000002 | chr21 | + | ENSG000002 | ENSG000002 | TRUE | cassette^474\|alt3^206\|a | | C1_A | chr21: | FALSE | FALSE | TRUE | 0.001 | 0.216 | 0.191 | 0.999 | 0 | cassette | 0.191 |
| 379 | ENSG000002 | ENSG000002 | ENSG000002 | chr21 | + | ENSG000002 | ENSG000002 | TRUE | cassette^474\|alt3^206\|a | | C1_A | chr21: | FALSE | FALSE | TRUE | 0.003 | 0.22 | 0.193 | 1 | 0 | cassette | 0.193 |

| | | | | | | | | | | | | | | | | | | | | | | |
|---|---|---|---|---|---|---|---|---|---|---|---|---|---|---|---|---|---|---|---|---|---|---|
| 380 | ENSG000002 | ENSG000002 | ENSG000002 | chr21 | + | ENSG000002 | ENSG000002 | TRUE | cassette^474\|alt3^206\|a | | C1_A | chr21: | FALSE | FALSE | TRUE | 0.015 | 0.162 | 0.135 | 0.993 | 0.003 | cassette | 0.135 |
| 381 | ENSG000002 | ENSG000002 | ENSG000002 | chr21 | + | ENSG000002 | ENSG000002 | TRUE | cassette^474\|alt3^206\|a | | C1_A | chr21: | FALSE | FALSE | TRUE | 0.001 | 0.296 | 0.284 | 1 | 0 | cassette | 0.284 |
| 382 | ENSG000002 | ENSG000002 | ENSG000002 | chr21 | + | ENSG000002 | ENSG000002 | TRUE | cassette^474\|alt3^206\|a | | C1_A | chr21: | FALSE | FALSE | TRUE | 0.006 | 0.223 | 0.204 | 1 | 0 | cassette | 0.204 |
| 383 | ENSG000002 | ENSG000002 | ENSG000002 | chr21 | + | ENSG000002 | ENSG000002 | TRUE | cassette^474\|alt3^206\|a | | C1_A | chr21: | FALSE | TRUE | TRUE | 0.019 | 0.204 | 0.181 | 1 | 0 | cassette | 0.181 |
| 384 | ENSG000002 | ENSG000002 | ENSG000002 | chr21 | + | ENSG000002 | ENSG000002 | TRUE | cassette^474\|alt3^206\|a | | C1_A | chr21: | FALSE | FALSE | FALSE | 0.012 | 0.141 | 0.107 | 0.911 | 0.044 | cassette | 0.107 |
| 385 | ENSG000002 | ENSG000002 | ENSG000002 | chr21 | + | ENSG000002 | ENSG000002 | TRUE | cassette^474\|alt3^206\|a | | C1_A | chr21: | FALSE | FALSE | TRUE | 0.004 | 0.139 | 0.109 | 1 | 0 | cassette | 0.109 |
| 386 | ENSG000002 | ENSG000002 | ENSG000002 | chr21 | + | ENSG000002 | ENSG000002 | TRUE | cassette^474\|alt3^206\|a | | C1_A | chr21: | FALSE | FALSE | TRUE | 0.011 | 0.214 | 0.184 | 0.985 | 0.004 | cassette | 0.184 |
| 387 | ENSG000002 | ENSG000002 | ENSG000002 | chr21 | + | ENSG000002 | ENSG000002 | TRUE | cassette^474\|alt3^206\|a | | C1_A | chr21: | FALSE | FALSE | TRUE | 0.001 | 0.146 | 0.127 | 0.974 | 0.011 | cassette | 0.127 |
| 388 | ENSG000002 | ENSG000002 | ENSG000002 | chr21 | + | ENSG000002 | ENSG000002 | TRUE | cassette^47 | TRUE | Distal | chr21:84004 | FALSE | FALSE | TRUE | 0.003 | 0.212 | 0.183 | 1 | 0 | AFE/ALE | 0.183 |
| 389 | ENSG000002 | ENSG000002 | ENSG000002 | chr21 | + | ENSG000002 | ENSG000002 | TRUE | cassette^47 | TRUE | C1_A | chr21:84005 | FALSE | FALSE | TRUE | 0.001 | 0.249 | 0.232 | 1 | 0 | cassette | 0.232 |
| 390 | ENSG000002 | ENSG000002 | ENSG000002 | chr21 | + | ENSG000002 | ENSG000002 | TRUE | cassette^474\|alt3^206\|a | | C1_A | chr21: | FALSE | FALSE | TRUE | 0.035 | 0.271 | 0.233 | 0.992 | 0.001 | cassette | 0.233 |
| 391 | ENSG000002 | ENSG000002 | ENSG000002 | chr21 | + | ENSG000002 | ENSG000002 | TRUE | cassette^47 | TRUE | C1_A | chr21:84011 | FALSE | FALSE | TRUE | 0.003 | 0.198 | 0.173 | 1 | 0 | cassette | 0.173 |
| 392 | ENSG000002 | ENSG000002 | ENSG000002 | chr21 | + | ENSG000002 | ENSG000002 | TRUE | cassette^474\|alt3^206\|a | | C1_A | chr21: | FALSE | FALSE | TRUE | 0.005 | 0.128 | 0.104 | 0.953 | 0.024 | cassette | 0.104 |
| 393 | ENSG000002 | ENSG000002 | ENSG000002 | chr21 | + | ENSG000002 | ENSG000002 | TRUE | cassette^474\|alt3^206\|a | | C1_A | chr21: | FALSE | FALSE | TRUE | 0.022 | 0.276 | 0.246 | 1 | 0 | cassette | 0.246 |
| 394 | ENSG000002 | ENSG000002 | ENSG000002 | chr21 | + | ENSG000002 | ENSG000002 | TRUE | cassette^474\|alt3^206\|a | | C1_A | chr21: | FALSE | FALSE | FALSE | 0.036 | 0.253 | 0.194 | 0.907 | 0.009 | cassette | 0.194 |
| 395 | ENSG000002 | ENSG000002 | ENSG000002 | chr21 | + | ENSG000002 | ENSG000002 | TRUE | cassette^474\|alt3^206\|a | | C1_A | chr21: | FALSE | FALSE | FALSE | 0.016 | 0.13 | 0.104 | 0.922 | 0.04 | cassette | 0.104 |
| 396 | ENSG000002 | ENSG000002 | ENSG000002 | chr21 | + | ENSG000002 | ENSG000002 | TRUE | cassette^474\|alt3^206\|a | | C1_A | chr21: | FALSE | FALSE | TRUE | 0 | 0.156 | 0.131 | 0.993 | 0.003 | cassette | 0.131 |
| 397 | ENSG000002 | ENSG000002 | ENSG000002 | chr21 | + | ENSG000002 | ENSG000002 | TRUE | cassette^474\|alt3^206\|a | | C1_A | chr21: | FALSE | FALSE | TRUE | 0.001 | 0.139 | 0.117 | 0.986 | 0.006 | cassette | 0.117 |
| 398 | ENSG000002 | ENSG000002 | ENSG000002 | chr21 | + | ENSG000002 | ENSG000002 | TRUE | cassette^474\|alt3^206\|a | | C1_A | chr21: | FALSE | FALSE | TRUE | 0.002 | 0.206 | 0.189 | 0.988 | 0.003 | cassette | 0.189 |
| 399 | ENSG000002 | ENSG000002 | ENSG000002 | chr21 | + | ENSG000002 | ENSG000002 | TRUE | cassette^474\|alt3^206\|a | | C1_A | chr21: | FALSE | FALSE | TRUE | 0.002 | 0.222 | 0.209 | 0.994 | 0.001 | cassette | 0.209 |
| 400 | ENSG000002 | ENSG000002 | ENSG000002 | chr21 | + | ENSG000002 | ENSG000002 | TRUE | cassette^474\|alt3^206\|a | | C1_A | chr21: | FALSE | FALSE | TRUE | 0 | 0.234 | 0.215 | 1 | 0 | cassette | 0.215 |
| 401 | ENSG000002 | ENSG000002 | ENSG000002 | chr21 | + | ENSG000002 | ENSG000002 | TRUE | cassette^474\|alt3^206\|a | | C1_A | chr21: | FALSE | FALSE | TRUE | 0 | 0.295 | 0.27 | 1 | 0 | cassette | 0.27 |
| 402 | ENSG000002 | ENSG000002 | ENSG000002 | chr21 | + | ENSG000002 | ENSG000002 | TRUE | cassette^474\|alt3^206\|a | | C1_A | chr21: | FALSE | FALSE | FALSE | 0.029 | 0.156 | 0.135 | 0.939 | 0.015 | cassette | 0.135 |
| 403 | ENSG000002 | ENSG000002 | ENSG000002 | chr21 | + | ENSG000002 | ENSG000002 | TRUE | cassette^474\|alt3^206\|a | | C1_A | chr21: | FALSE | FALSE | TRUE | 0.026 | 0.212 | 0.193 | 0.982 | 0.004 | cassette | 0.193 |
| 404 | ENSG000002 | ENSG000002 | ENSG000002 | chr21 | + | ENSG000002 | ENSG000002 | TRUE | cassette^474\|alt3^206\|a | | C1_A | chr21: | FALSE | FALSE | TRUE | 0.045 | 0.236 | 0.211 | 0.992 | 0.002 | cassette | 0.211 |
| 405 | ENSG000002 | ENSG000002 | ENSG000002 | chr21 | + | ENSG000002 | ENSG000002 | TRUE | cassette^474\|alt3^206\|a | | C1_A | chr21: | FALSE | FALSE | TRUE | 0.048 | 0.224 | 0.18 | 0.982 | 0.005 | cassette | 0.18 |
| 406 | ENSG000002 | ENSG000002 | ENSG000002 | chr21 | + | ENSG000002 | ENSG000002 | TRUE | cassette^474\|alt3^206\|a | | C1_A | chr21: | FALSE | FALSE | TRUE | 0 | 0.15 | 0.133 | 0.976 | 0.01 | cassette | 0.133 |
| 407 | ENSG000002 | ENSG000002 | ENSG000002 | chr21 | + | ENSG000002 | ENSG000002 | TRUE | cassette^474\|alt3^206\|a | | C1_A | chr21: | FALSE | FALSE | TRUE | 0 | 0.145 | 0.134 | 0.952 | 0.018 | cassette | 0.134 |
| 408 | ENSG000002 | ENSG000002 | ENSG000002 | chr21 | + | ENSG000002 | ENSG000002 | TRUE | cassette^474\|alt3^206\|a | | C1_A | chr21: | FALSE | FALSE | TRUE | 0 | 0.422 | 0.401 | 1 | 0 | cassette | 0.401 |
| 409 | ENSG000002 | ENSG000002 | ENSG000002 | chr21 | + | ENSG000002 | ENSG000002 | TRUE | cassette^474\|alt3^206\|a | | C1_A | chr21: | FALSE | FALSE | TRUE | 0 | 0.395 | 0.383 | 1 | 0 | cassette | 0.383 |
| 410 | ENSG000002 | ENSG000002 | ENSG000002 | chr21 | + | ENSG000002 | ENSG000002 | TRUE | cassette^474\|alt3^206\|a | | C1_A | chr21: | FALSE | FALSE | TRUE | 0 | 0.232 | 0.211 | 1 | 0 | cassette | 0.211 |
| 411 | ENSG000002 | ENSG000002 | ENSG000002 | chr21 | + | ENSG000002 | ENSG000002 | TRUE | cassette^474\|alt3^206\|a | | C1_A | chr21: | FALSE | FALSE | TRUE | 0 | 0.261 | 0.231 | 0.997 | 0 | cassette | 0.231 |
| 412 | ENSG000002 | ENSG000002 | ENSG000002 | chr21 | + | ENSG000002 | ENSG000002 | TRUE | cassette^474\|alt3^206\|a | | C1_A | chr21: | FALSE | FALSE | TRUE | 0.001 | 0.193 | 0.168 | 0.981 | 0.005 | cassette | 0.168 |
| 413 | ENSG000002 | ENSG000002 | ENSG000002 | chr21 | + | ENSG000002 | ENSG000002 | TRUE | cassette^474\|alt3^206\|a | | C1_A | chr21: | FALSE | FALSE | TRUE | 0.011 | 0.286 | 0.264 | 1 | 0 | cassette | 0.264 |
| 414 | ENSG000002 | ENSG000002 | ENSG000002 | chr21 | + | ENSG000002 | ENSG000002 | TRUE | cassette^474\|alt3^206\|a | | C1_A | chr21: | FALSE | FALSE | FALSE | 0.024 | 0.137 | 0.111 | 0.927 | 0.03 | cassette | 0.111 |
| 415 | ENSG000002 | ENSG000002 | ENSG000002 | chr21 | + | ENSG000002 | ENSG000002 | TRUE | cassette^474\|alt3^206\|a | | C1_A | chr21: | FALSE | FALSE | TRUE | 0.007 | 0.228 | 0.191 | 0.955 | 0.006 | cassette | 0.191 |
| 416 | ENSG000002 | ENSG000002 | ENSG000002 | chr21 | + | ENSG000002 | ENSG000002 | TRUE | cassette^474\|alt3^206\|a | | C1_A | chr21: | FALSE | FALSE | TRUE | 0.007 | 0.385 | 0.33 | 0.995 | 0 | cassette | 0.33 |
| 417 | ENSG000002 | ENSG000002 | ENSG000002 | chr21 | + | ENSG000002 | ENSG000002 | TRUE | cassette^474\|alt3^206\|a | | C1_A | chr21: | FALSE | FALSE | TRUE | 0.001 | 0.588 | 0.576 | 1 | 0 | cassette | 0.576 |
| 418 | ENSG000002 | ENSG000002 | ENSG000002 | chr21 | + | ENSG000002 | ENSG000002 | TRUE | cassette^474\|alt3^206\|a | | C1_A | chr21: | FALSE | FALSE | TRUE | 0.001 | 0.175 | 0.147 | 0.991 | 0.003 | cassette | 0.147 |
| 419 | ENSG000002 | ENSG000002 | ENSG000002 | chr21 | + | ENSG000002 | ENSG000002 | TRUE | cassette^474\|alt3^206\|a | | C1_A | chr21: | FALSE | FALSE | TRUE | 0.024 | 0.274 | 0.248 | 0.977 | 0.002 | cassette | 0.248 |
| 420 | ENSG000002 | ENSG000002 | ENSG000002 | chr21 | + | ENSG000002 | ENSG000002 | TRUE | cassette^474\|alt3^206\|a | | C1_A | chr21: | FALSE | FALSE | FALSE | 0.003 | 0.208 | 0.173 | 0.945 | 0.011 | cassette | 0.173 |
| 421 | ENSG000002 | ENSG000002 | ENSG000002 | chr21 | + | ENSG000002 | ENSG000002 | TRUE | cassette^474\|alt3^206\|a | | C1_A | chr21: | FALSE | FALSE | TRUE | 0.003 | 0.293 | 0.268 | 1 | 0 | cassette | 0.268 |
| 422 | ENSG000002 | ENSG000002 | ENSG000002 | chr21 | + | ENSG000002 | ENSG000002 | TRUE | cassette^474\|alt3^206\|a | | C1_A | chr21: | FALSE | FALSE | TRUE | 0 | 0.141 | 0.116 | 0.954 | 0.021 | cassette | 0.116 |
| 423 | ENSG000002 | ENSG000002 | ENSG000002 | chr21 | + | ENSG000002 | ENSG000002 | TRUE | cassette^474\|alt3^206\|a | | C1_A | chr21: | FALSE | FALSE | TRUE | 0.006 | 0.425 | 0.394 | 1 | 0 | cassette | 0.394 |
| 424 | ENSG000002 | ENSG000002 | ENSG000002 | chr21 | + | ENSG000002 | ENSG000002 | TRUE | cassette^474\|alt3^206\|a | | C1_A | chr21: | FALSE | FALSE | TRUE | 0.007 | 0.251 | 0.224 | 0.998 | 0 | cassette | 0.224 |
| 425 | ENSG000002 | ENSG000002 | ENSG000002 | chr21 | + | ENSG000002 | ENSG000002 | TRUE | cassette^474\|alt3^206\|a | | C1_A | chr21: | FALSE | FALSE | TRUE | 0.001 | 0.338 | 0.3 | 0.996 | 0 | cassette | 0.3 |
| 426 | ENSG000002 | ENSG000002 | ENSG000002 | chr21 | + | ENSG000002 | ENSG000002 | TRUE | cassette^474\|alt3^206\|a | | C1_A | chr21: | FALSE | FALSE | TRUE | 0.004 | 0.251 | 0.233 | 0.999 | 0 | cassette | 0.233 |
| 427 | ENSG000002 | ENSG000002 | ENSG000002 | chr21 | + | ENSG000002 | ENSG000002 | TRUE | cassette^47 | TRUE | Distal | chr21:84404 | FALSE | FALSE | TRUE | 0.001 | 0.338 | 0.3 | 0.996 | 0 | AFE/ALE | 0.3 |
| 428 | ENSG000002 | ENSG000002 | ENSG000002 | chr21 | + | ENSG000002 | ENSG000002 | TRUE | cassette^474\|alt3^206\|a | | C1_A | chr21: | FALSE | FALSE | TRUE | 0.015 | 0.438 | 0.416 | 1 | 0 | cassette | 0.416 |
| 429 | ENSG000002 | ENSG000002 | ENSG000002 | chr21 | + | ENSG000002 | ENSG000002 | TRUE | cassette^474\|alt3^206\|a | | C1_A | chr21: | FALSE | FALSE | TRUE | 0.008 | 0.993 | 0.973 | 1 | 0 | cassette | 0.973 |
| 430 | ENSG000002 | ENSG000002 | ENSG000002 | chr21 | + | ENSG000002 | ENSG000002 | TRUE | cassette^474\|alt3^206\|a | | C1_A | chr21: | FALSE | FALSE | TRUE | 0.005 | 0.207 | 0.183 | 0.988 | 0.003 | cassette | 0.183 |
| 431 | ENSG000002 | ENSG000002 | ENSG000002 | chr21 | + | ENSG000002 | ENSG000002 | TRUE | cassette^47 | TRUE | Distal | chr21:84448 | FALSE | FALSE | TRUE | 0.001 | 0.176 | 0.149 | 0.958 | 0.009 | AFE/ALE | 0.149 |
| 432 | ENSG000002 | ENSG000002 | ENSG000002 | chr21 | + | ENSG000002 | ENSG000002 | TRUE | cassette^474\|alt3^206\|a | | C1_A | chr21: | FALSE | FALSE | TRUE | 0.003 | 0.176 | 0.149 | 0.966 | 0.012 | cassette | 0.149 |
| 433 | ENSG000002 | ENSG000002 | ENSG000002 | chr21 | + | ENSG000002 | ENSG000002 | TRUE | cassette^47 | TRUE | Proximal | chr21:83819 | FALSE | TRUE | TRUE | 0.001 | 0.824 | 0.815 | 1 | 0 | AFE/ALE | 0.815 |
| 434 | ENSG000002 | ENSG000002 | ENSG000002 | chr21 | + | ENSG000002 | ENSG000002 | TRUE | cassette^47 | TRUE | C1_A2 | chr21:83825 | FALSE | FALSE | TRUE | 0 | 0.313 | 0.296 | 1 | 0 | cassette | 0.296 |
| 435 | ENSG000002 | ENSG000002 | ENSG000002 | chr21 | + | ENSG000002 | ENSG000002 | TRUE | cassette^47 | TRUE | Distal | chr21:83889 | FALSE | FALSE | TRUE | 0 | 0.586 | 0.579 | 1 | 0 | AFE/ALE | 0.579 |
| 436 | ENSG000002 | ENSG000002 | ENSG000002 | chr21 | + | ENSG000002 | ENSG000002 | TRUE | cassette^47 | TRUE | Distal | chr21:83903 | FALSE | FALSE | FALSE | 0.035 | 0.214 | 0.168 | 0.937 | 0.011 | AFE/ALE | 0.168 |
| 437 | ENSG000002 | ENSG000002 | ENSG000002 | chr21 | + | ENSG000002 | ENSG000002 | TRUE | cassette^47 | TRUE | Proximal | chr21:83908 | FALSE | TRUE | TRUE | 0.001 | 0.259 | 0.234 | 1 | 0 | AFE/ALE | 0.234 |
| 438 | ENSG000002 | ENSG000002 | ENSG000002 | chr21 | + | ENSG000002 | ENSG000002 | TRUE | cassette^47 | TRUE | Proximal | chr21:83908 | FALSE | FALSE | TRUE | 0.007 | 0.214 | 0.192 | 0.991 | 0.002 | AFE/ALE | 0.192 |
| 439 | ENSG000002 | ENSG000002 | ENSG000002 | chr21 | + | ENSG000002 | ENSG000002 | TRUE | cassette^47 | TRUE | Distal | chr21:83909 | FALSE | FALSE | TRUE | 0.006 | 0.21 | 0.176 | 0.974 | 0.007 | AFE/ALE | 0.176 |
| 440 | ENSG000002 | ENSG000002 | ENSG000002 | chr21 | + | ENSG000002 | ENSG000002 | TRUE | cassette^47 | TRUE | Distal | chr21:83914 | FALSE | TRUE | TRUE | 0.002 | 0.364 | 0.339 | 1 | 0 | AFE/ALE | 0.339 |
| 441 | ENSG000002 | ENSG000002 | ENSG000002 | chr21 | + | ENSG000002 | ENSG000002 | TRUE | cassette^47 | TRUE | C1_A2 | chr21:83919 | FALSE | FALSE | TRUE | 0.006 | 0.484 | 0.468 | 1 | 0 | cassette | 0.468 |
| 442 | ENSG000002 | ENSG000002 | ENSG000002 | chr21 | + | ENSG000002 | ENSG000002 | TRUE | cassette^47 | TRUE | Proximal | chr21:84208 | FALSE | TRUE | TRUE | 0.005 | 0.72 | 0.69 | 1 | 0 | AFE/ALE | 0.69 |
| 443 | ENSG000002 | ENSG000002 | ENSG000002 | chr21 | + | ENSG000002 | ENSG000002 | TRUE | cassette^47 | TRUE | C1_C2 | chr21:84299 | FALSE | FALSE | TRUE | 0.001 | 0.242 | -0.21 | 0.991 | 0.002 | skipping | 0.21 |
| 444 | ENSG000002 | ENSG000002 | ENSG000002 | chr21 | + | ENSG000002 | ENSG000002 | TRUE | cassette^47 | TRUE | Proximal | chr21:83852 | FALSE | FALSE | TRUE | 0.007 | 0.264 | 0.239 | 1 | 0 | AFE/ALE | 0.239 |
| 445 | ENSG000002 | ENSG000002 | ENSG000002 | chr21 | + | ENSG000002 | ENSG000002 | TRUE | cassette^47 | TRUE | Proximal | chr21:83902 | FALSE | FALSE | TRUE | 0.003 | 0.186 | 0.16 | 0.995 | 0.002 | AFE/ALE | 0.16 |
| 446 | ENSG000002 | ENSG000002 | ENSG000002 | chr21 | + | ENSG000002 | ENSG000002 | TRUE | cassette^47 | TRUE | E2_E1_J1 | chr21:83810 | FALSE | FALSE | TRUE | 0.007 | 0.429 | 0.39 | 1 | 0 | cassette | 0.39 |
| 447 | ENSG000002 | ENSG000002 | ENSG000002 | chr21 | + | ENSG000002 | ENSG000002 | TRUE | cassette^474\|alt3^206\|a | | A_C2 | chr21: | FALSE | FALSE | TRUE | 0.001 | 0.242 | 0.22 | 0.989 | 0.002 | cassette | 0.22 |
| 448 | ENSG000002 | ENSG000002 | ENSG000002 | chr21 | + | ENSG000002 | ENSG000002 | TRUE | cassette^47 | TRUE | Proximal | chr21:83896 | FALSE | FALSE | TRUE | 0.026 | 0.589 | 0.546 | 1 | 0 | AFE/ALE | 0.546 |
| 449 | ENSG000002 | ENSG000002 | ENSG000002 | chr21 | + | ENSG000002 | ENSG000002 | TRUE | cassette^47 | TRUE | Proximal | chr21:83900 | FALSE | FALSE | FALSE | 0.001 | 0.176 | 0.142 | 0.926 | 0.025 | AFE/ALE | 0.142 |
| 450 | ENSG000002 | ENSG000002 | ENSG000002 | chr21 | + | ENSG000002 | ENSG000002 | TRUE | cassette^47 | TRUE | Proximal | chr21:83819 | FALSE | FALSE | TRUE | 0 | 0.158 | 0.135 | 1 | 0 | AFE/ALE | 0.135 |
| 451 | ENSG000002 | ENSG000002 | ENSG000002 | chr21 | + | ENSG000002 | ENSG000002 | TRUE | cassette^47 | TRUE | E2_E1_J1 | chr21:83843 | FALSE | FALSE | FALSE | 0.043 | 0.36 | 0.279 | 0.933 | 0.003 | cassette | 0.279 |
| 452 | ENSG000002 | ENSG000002 | ENSG000002 | chr21 | + | ENSG000002 | ENSG000002 | TRUE | cassette^474\|alt3^206\|a | | A_C2 | chr21: | FALSE | FALSE | TRUE | 0.001 | 0.204 | 0.187 | 0.987 | 0.003 | cassette | 0.187 |
| 453 | ENSG000002 | ENSG000002 | ENSG000002 | chr21 | + | ENSG000002 | ENSG000002 | TRUE | cassette^47 | TRUE | Proximal | chr21:83896 | FALSE | FALSE | TRUE | 0.031 | 0.209 | 0.176 | 0.994 | 0.002 | AFE/ALE | 0.176 |
| 454 | ENSG000002 | ENSG000002 | ENSG000002 | chr21 | + | ENSG000002 | ENSG000002 | TRUE | cassette^47 | TRUE | Proximal | chr21:83881 | FALSE | FALSE | TRUE | 0 | 0.209 | 0.187 | 0.977 | 0.005 | AFE/ALE | 0.187 |
| 455 | ENSG000002 | ENSG000002 | ENSG000002 | chr21 | + | ENSG000002 | ENSG000002 | TRUE | cassette^47 | TRUE | Proximal | chr21:83823 | FALSE | FALSE | FALSE | 0.005 | 0.156 | 0.142 | 0.902 | 0.026 | AFE/ALE | 0.142 |

| | | | | | | | | | | | | | | | | | | | | | | |
|---|---|---|---|---|---|---|---|---|---|---|---|---|---|---|---|---|---|---|---|---|---|---|
| 456 | ENSG000002 | ENSG000002 | ENSG000002 | chr21 | + | ENSG000002 | ENSG000002 | TRUE | cassette^474 | TRUE | Distal | chr21:83899 | FALSE | FALSE | TRUE | 0.001 | 0.217 | 0.197 | 0.977 | 0.005 | AFE/ALE | 0.197 |
| 457 | ENSG000002 | ENSG000002 | ENSG000002 | chr21 | + | ENSG000002 | ENSG000002 | TRUE | cassette^474\|alt3^206\|a | | A_C2 | chr21: | FALSE | FALSE | FALSE | 0 | 0.146 | 0.144 | 0.907 | 0.025 | cassette | 0.144 |
| 458 | ENSG000002 | ENSG000002 | ENSG000002 | chr21 | + | ENSG000002 | ENSG000002 | TRUE | cassette^474 | TRUE | Distal | chr21:83893 | FALSE | FALSE | TRUE | 0 | 0.446 | 0.439 | 1 | 0 | AFE/ALE | 0.439 |
| 459 | ENSG000002 | ENSG000002 | ENSG000002 | chr21 | + | ENSG000002 | ENSG000002 | TRUE | cassette^474 | TRUE | E2_E1_J2 | chr21:83958 | FALSE | FALSE | TRUE | 0.045 | 0.176 | 0.136 | 0.972 | 0.011 | cassette | 0.136 |
| 460 | ENSG000002 | ENSG000002 | ENSG000002 | chr21 | + | ENSG000002 | ENSG000002 | TRUE | cassette^474\|alt3^206\|a | | A_C2 | chr21: | FALSE | FALSE | TRUE | 0 | 0.145 | 0.119 | 0.955 | 0.021 | cassette | 0.119 |
| 461 | ENSG000002 | ENSG000002 | ENSG000002 | chr21 | + | ENSG000002 | ENSG000002 | TRUE | cassette^474\|alt3^206\|a | | A_C2 | chr21: | FALSE | FALSE | FALSE | 0.005 | 0.154 | 0.12 | 0.925 | 0.03 | cassette | 0.12 |
| 462 | ENSG000002 | ENSG000002 | ENSG000002 | chr21 | + | ENSG000002 | ENSG000002 | TRUE | cassette^474\|alt3^206\|a | | A_C2 | chr21: | FALSE | FALSE | TRUE | 0.001 | 0.292 | 0.261 | 1 | 0 | cassette | 0.261 |
| 463 | ENSG000002 | ENSG000002 | ENSG000002 | chr21 | + | ENSG000002 | ENSG000002 | TRUE | cassette^474\|alt3^206\|a | | A_C2 | chr21: | FALSE | FALSE | TRUE | 0.003 | 0.162 | 0.137 | 0.983 | 0.007 | cassette | 0.137 |
| 464 | ENSG000002 | ENSG000002 | ENSG000002 | chr21 | + | ENSG000002 | ENSG000002 | TRUE | cassette^474 | TRUE | Proximal | chr21:83925 | FALSE | FALSE | TRUE | 0.004 | 0.285 | 0.255 | 0.984 | 0.001 | AFE/ALE | 0.255 |
| 465 | ENSG000002 | ENSG000002 | ENSG000002 | chr21 | + | ENSG000002 | ENSG000002 | TRUE | cassette^474 | TRUE | A_C2 | chr21:83951 | FALSE | FALSE | FALSE | 0.001 | 0.164 | 0.147 | 0.946 | 0.016 | cassette | 0.147 |
| 466 | ENSG000002 | ENSG000002 | ENSG000002 | chr21 | + | ENSG000002 | ENSG000002 | TRUE | cassette^474\|alt3^206\|a | | A_C2 | chr21: | FALSE | FALSE | TRUE | 0.017 | 0.284 | 0.249 | 0.982 | 0.002 | cassette | 0.249 |
| 467 | ENSG000002 | ENSG000002 | ENSG000002 | chr21 | + | ENSG000002 | ENSG000002 | TRUE | cassette^474 | TRUE | Proximal | chr21:83886 | FALSE | FALSE | TRUE | 0.005 | 0.397 | 0.38 | 1 | 0 | AFE/ALE | 0.38 |
| 468 | ENSG000002 | ENSG000002 | ENSG000002 | chr21 | + | ENSG000002 | ENSG000002 | TRUE | cassette^474 | TRUE | Distal | chr21:83825 | FALSE | FALSE | TRUE | 0.001 | 0.498 | 0.503 | 1 | 0 | AFE/ALE | 0.503 |
| 469 | ENSG000002 | ENSG000002 | ENSG000002 | chr21 | + | ENSG000002 | ENSG000002 | TRUE | cassette^474\|alt3^206\|a | | A_C2 | chr21: | FALSE | FALSE | FALSE | 0 | 0.122 | 0.099 | 0.949 | 0.027 | cassette | 0.099 |
| 470 | ENSG000002 | ENSG000002 | ENSG000002 | chr21 | + | ENSG000002 | ENSG000002 | TRUE | cassette^474\|alt3^206\|a | | A_C2 | chr21: | FALSE | FALSE | TRUE | 0.046 | 0.255 | 0.208 | 1 | 0 | cassette | 0.208 |
| 471 | ENSG000002 | ENSG000002 | ENSG000002 | chr21 | + | ENSG000002 | ENSG000002 | TRUE | cassette^474\|alt3^206\|a | | A_C2 | chr21: | FALSE | FALSE | TRUE | 0.011 | 0.724 | 0.706 | 1 | 0 | cassette | 0.706 |
| 472 | ENSG000002 | ENSG000002 | ENSG000002 | chr21 | + | ENSG000002 | ENSG000002 | TRUE | cassette^474 | TRUE | C2_A | chr21:83988 | FALSE | FALSE | TRUE | 0.042 | 0.232 | 0.195 | 1 | 0 | cassette | 0.195 |
| 473 | ENSG000002 | ENSG000002 | ENSG000002 | chr21 | + | ENSG000002 | ENSG000002 | TRUE | cassette^474\|alt3^206\|a | | A_C2 | chr21: | FALSE | FALSE | TRUE | 0.034 | 0.27 | 0.229 | 0.999 | 0 | cassette | 0.229 |
| 474 | ENSG000002 | ENSG000002 | ENSG000002 | chr21 | + | ENSG000002 | ENSG000002 | TRUE | cassette^474\|alt3^206\|a | | A_C2 | chr21: | FALSE | FALSE | TRUE | 0.001 | 0.383 | 0.357 | 1 | 0 | cassette | 0.357 |
| 475 | ENSG000002 | ENSG000002 | ENSG000002 | chr21 | + | ENSG000002 | ENSG000002 | TRUE | cassette^474 | TRUE | Distal | chr21:83824 | FALSE | FALSE | TRUE | 0 | 0.128 | 0.105 | 0.954 | 0.023 | AFE/ALE | 0.105 |
| 476 | ENSG000002 | ENSG000002 | ENSG000002 | chr21 | + | ENSG000002 | ENSG000002 | TRUE | cassette^474 | TRUE | Distal | chr21:83889 | FALSE | FALSE | TRUE | 0 | 0.386 | 0.358 | 1 | 0 | AFE/ALE | 0.358 |
| 477 | ENSG000002 | ENSG000002 | ENSG000002 | chr21 | + | ENSG000002 | ENSG000002 | TRUE | cassette^474\|alt3^206\|a | | A_C2 | chr21: | FALSE | FALSE | TRUE | 0.02 | 0.241 | 0.207 | 0.999 | 0 | cassette | 0.207 |
| 478 | ENSG000002 | ENSG000002 | ENSG000002 | chr21 | + | ENSG000002 | ENSG000002 | TRUE | cassette^474 | TRUE | C2_A1 | chr21:84011 | FALSE | FALSE | TRUE | 0.011 | 0.272 | 0.25 | 1 | 0 | cassette | 0.25 |
| 479 | ENSG000002 | ENSG000002 | ENSG000002 | chr21 | + | ENSG000002 | ENSG000002 | TRUE | cassette^474 | TRUE | Distal | chr21:83939 | FALSE | FALSE | TRUE | 0.019 | 0.498 | 0.492 | 1 | 0 | AFE/ALE | 0.492 |
| 480 | ENSG000002 | ENSG000002 | ENSG000002 | chr21 | + | ENSG000002 | ENSG000002 | TRUE | cassette^474 | TRUE | Proximal | chr21:84299 | FALSE | FALSE | TRUE | 0.001 | 0.409 | 0.37 | 0.998 | 0 | AFE/ALE | 0.37 |
| 481 | ENSG000002 | ENSG000002 | ENSG000002 | chr21 | + | ENSG000002 | ENSG000002 | TRUE | cassette^474\|alt3^206\|a | | A_C2 | chr21: | FALSE | FALSE | TRUE | 0.001 | 0.336 | 0.309 | 0.999 | 0 | cassette | 0.309 |
| 482 | ENSG000002 | ENSG000002 | ENSG000002 | chr21 | + | ENSG000002 | ENSG000002 | TRUE | cassette^474 | TRUE | A_C2 | chr21:84078 | FALSE | FALSE | FALSE | 0.011 | 0.118 | 0.099 | 0.916 | 0.045 | cassette | 0.099 |
| 483 | ENSG000002 | ENSG000002 | ENSG000002 | chr21 | + | ENSG000002 | ENSG000002 | TRUE | cassette^474 | TRUE | A_C2 | chr21:84088 | FALSE | FALSE | FALSE | 0.04 | 0.165 | 0.128 | 0.92 | 0.03 | cassette | 0.128 |
| 484 | ENSG000002 | ENSG000002 | ENSG000002 | chr21 | + | ENSG000002 | ENSG000002 | TRUE | cassette^474\|alt3^206\|a | | A_C2 | chr21: | FALSE | FALSE | TRUE | 0.004 | 0.296 | 0.268 | 1 | 0 | cassette | 0.268 |
| 485 | ENSG000002 | ENSG000002 | ENSG000002 | chr21 | + | ENSG000002 | ENSG000002 | TRUE | cassette^474 | TRUE | A_C2 | chr21:84107 | FALSE | FALSE | TRUE | 0.009 | 0.466 | 0.456 | 1 | 0 | cassette | 0.456 |
| 486 | ENSG000002 | ENSG000002 | ENSG000002 | chr21 | + | ENSG000002 | ENSG000002 | TRUE | cassette^474\|alt3^206\|a | | A_C2 | chr21: | FALSE | FALSE | TRUE | 0.012 | 0.254 | 0.225 | 1 | 0 | cassette | 0.225 |
| 487 | ENSG000002 | ENSG000002 | ENSG000002 | chr21 | + | ENSG000002 | ENSG000002 | TRUE | cassette^474\|alt3^206\|a | | A_C2 | chr21: | FALSE | FALSE | FALSE | 0.026 | 0.148 | 0.126 | 0.913 | 0.022 | cassette | 0.126 |
| 488 | ENSG000002 | ENSG000002 | ENSG000002 | chr21 | + | ENSG000002 | ENSG000002 | TRUE | cassette^474\|alt3^206\|a | | A_C2 | chr21: | FALSE | FALSE | TRUE | 0.003 | 0.184 | 0.156 | 0.957 | 0.013 | cassette | 0.156 |
| 489 | ENSG000002 | ENSG000002 | ENSG000002 | chr21 | + | ENSG000002 | ENSG000002 | TRUE | cassette^474\|alt3^206\|a | | A_C2 | chr21: | FALSE | FALSE | TRUE | 0.006 | 0.3 | 0.283 | 1 | 0 | cassette | 0.283 |
| 490 | ENSG000002 | ENSG000002 | ENSG000002 | chr21 | + | ENSG000002 | ENSG000002 | TRUE | cassette^474\|alt3^206\|a | | A_C2 | chr21: | FALSE | FALSE | FALSE | 0 | 0.146 | 0.125 | 0.93 | 0.029 | cassette | 0.125 |
| 491 | ENSG000002 | ENSG000002 | ENSG000002 | chr21 | + | ENSG000002 | ENSG000002 | TRUE | cassette^474\|alt3^206\|a | | A_C2 | chr21: | FALSE | FALSE | TRUE | 0.01 | 0.126 | 0.103 | 0.985 | 0.008 | cassette | 0.103 |
| 492 | ENSG000002 | ENSG000002 | ENSG000002 | chr21 | + | ENSG000002 | ENSG000002 | TRUE | cassette^474 | TRUE | Distal | chr21:83987 | FALSE | FALSE | FALSE | 0.001 | 0.171 | 0.143 | 0.929 | 0.022 | AFE/ALE | 0.143 |
| 493 | ENSG000002 | ENSG000002 | ENSG000002 | chr21 | + | ENSG000002 | ENSG000002 | TRUE | cassette^474\|alt3^206\|a | | A_C2 | chr21: | FALSE | FALSE | FALSE | 0.001 | 0.171 | 0.143 | 0.929 | 0.022 | cassette | 0.143 |
| 494 | ENSG000002 | ENSG000002 | ENSG000002 | chr21 | + | ENSG000002 | ENSG000002 | TRUE | cassette^474\|alt3^206\|a | | A_C2 | chr21: | FALSE | FALSE | TRUE | 0 | 0.169 | 0.142 | 0.985 | 0.006 | cassette | 0.142 |
| 495 | ENSG000002 | ENSG000002 | ENSG000002 | chr21 | + | ENSG000002 | ENSG000002 | TRUE | cassette^474\|alt3^206\|a | | A_C2 | chr21: | FALSE | FALSE | FALSE | 0.034 | 0.229 | 0.185 | 0.944 | 0.008 | cassette | 0.185 |
| 496 | ENSG000002 | ENSG000002 | ENSG000002 | chr21 | + | ENSG000002 | ENSG000002 | TRUE | cassette^474\|alt3^206\|a | | A_C2 | chr21: | FALSE | FALSE | TRUE | 0 | 0.182 | 0.152 | 0.99 | 0.004 | cassette | 0.152 |
| 497 | ENSG000002 | ENSG000002 | ENSG000002 | chr21 | + | ENSG000002 | ENSG000002 | TRUE | cassette^474 | TRUE | Proximal | chr21:84386 | FALSE | FALSE | FALSE | 0.023 | 0.181 | 0.153 | 0.936 | 0.016 | AFE/ALE | 0.153 |
| 498 | ENSG000002 | ENSG000002 | ENSG000002 | chr21 | + | ENSG000002 | ENSG000002 | TRUE | cassette^474\|alt3^206\|a | | A_C2 | chr21: | FALSE | FALSE | FALSE | 0.023 | 0.181 | 0.153 | 0.936 | 0.016 | cassette | 0.153 |
| 499 | ENSG000002 | ENSG000002 | ENSG000002 | chr21 | + | ENSG000002 | ENSG000002 | TRUE | cassette^474\|alt3^206\|a | | A_C2 | chr21: | FALSE | FALSE | TRUE | 0.013 | 0.283 | 0.263 | 1 | 0 | cassette | 0.263 |
| 500 | ENSG000002 | ENSG000002 | ENSG000002 | chr21 | + | ENSG000002 | ENSG000002 | TRUE | cassette^474\|alt3^206\|a | | A_C2 | chr21: | FALSE | FALSE | TRUE | 0 | 0.244 | 0.22 | 1 | 0 | cassette | 0.22 |
| 501 | ENSG000002 | ENSG000002 | ENSG000002 | chr21 | + | ENSG000002 | ENSG000002 | TRUE | cassette^474 | TRUE | C2_A1 | chr21:84448 | FALSE | FALSE | TRUE | 0.001 | 0.207 | 0.174 | 0.971 | 0.004 | cassette | 0.174 |
| 502 | ENSG000002 | ENSG000002 | ENSG000002 | chr1 | - | ENSG000002 | ENSG000002 | TRUE | putative_afe | TRUE | Distal | chr1:167406 | FALSE | TRUE | TRUE | 0.001 | 0.227 | 0.21 | 1 | 0 | AFE/ALE | 0.21 |
| 503 | ENSG000002 | ENSG000002 | ENSG000002 | chr14 | - | ENSG000002 | ENSG000002 | TRUE | alt3^4\|alt5^ | TRUE | E1_E2_J2 | chr14:49862 | FALSE | FALSE | FALSE | 0 | 0.121 | 0.098 | 0.918 | 0.044 | cassette | 0.098 |
| 504 | ENSG000002 | ENSG000002 | ENSG000002 | chr19 | + | ENSG000002 | ENSG000002 | TRUE | alt3^1\|putat | TRUE | Proximal | chr19:35558 | FALSE | FALSE | TRUE | 0.001 | 0.165 | 0.137 | 0.959 | 0.016 | AFE/ALE | 0.137 |
| 505 | ENSG000002 | ENSG000002 | ENSG000002 | chr19 | + | ENSG000002 | ENSG000002 | TRUE | alt3^1\|putat | TRUE | Proximal | chr19:35559 | FALSE | FALSE | TRUE | 0.001 | 0.162 | 0.133 | 0.966 | 0.014 | AFE/ALE | 0.133 |
| 506 | ENSG000002 | ENSG000002 | ENSG000002 | chr19 | + | ENSG000002 | ENSG000002 | TRUE | alt3^1\|putat | TRUE | Proximal | chr19:35563 | FALSE | FALSE | TRUE | 0.001 | 0.229 | 0.198 | 1 | 0 | AFE/ALE | 0.198 |
| 507 | ENSG000002 | ENSG000002 | ENSG000002 | chr19 | + | ENSG000002 | ENSG000002 | TRUE | cassette^2\|p | TRUE | C1_A | chr19:58583 | FALSE | FALSE | TRUE | 0.011 | 0.25 | 0.223 | 1 | 0 | cassette | 0.223 |

Table S2: Identified cryptic exons in hnRNPA1 KD neurons.

| | module_id | gene_id | gene_name | seqid | strand | lsv_id | event_id | complex | module_ever | denovo | junction_nan | junctions_co | event_non_cl | event_chang | junction_cha | refPSI | KDSoma_me | deltaPSI | prob | padjust | event | abs(deltaPSI) |
|---|---|---|---|---|---|---|---|---|---|---|---|---|---|---|---|---|---|---|---|---|---|---|
| 1 | ENSG000001 | ENSG000001 | BRD8 | chr5 | - | | ENSG000001 | FALSE | putative_alt5 | FALSE | Proximal | chr5:138157 | FALSE | FALSE | TRUE | 0.017 | 0.147 | 0.119 | 0.961 | 0.016 | AFE/ALE | 0.119 |
| 2 | ENSG00000C | ENSG00000C | ZNF839 | chr14 | + | ENSG00000C | ENSG00000C | TRUE | ir^2\|afe^2 | FALSE | Distal | chr14:102317 | FALSE | FALSE | TRUE | 0.029 | 0.267 | 0.215 | 0.971 | 0.003 | AFE/ALE | 0.215 |
| 3 | ENSG00000C | ENSG00000C | ANK1 | chr8 | - | ENSG00000C | ENSG00000C | FALSE | putative_ale | TRUE | Proximal | chr8:418187 | FALSE | TRUE | TRUE | 0.005 | 0.369 | 0.346 | 1 | 0 | AFE/ALE | 0.346 |
| 4 | ENSG00000C | ENSG00000C | CROCC | chr1 | + | ENSG00000C | ENSG00000C | TRUE | ir^1\|ale^13\|afe^5\|putati | | C1_A | chr1: | FALSE | TRUE | TRUE | 0 | 0.517 | 0.505 | 1 | 0 | cassette | 0.505 |
| 5 | ENSG00000C | ENSG00000C | CROCC | chr1 | + | ENSG00000C | ENSG00000C | TRUE | ir^1\|ale^13\|afe^5\|putati | | A_C2 | chr1: | FALSE | FALSE | TRUE | 0.019 | 0.488 | 0.475 | 1 | 0 | cassette | 0.475 |
| 6 | ENSG00000C | ENSG00000C | CROCC | chr1 | + | ENSG00000C | ENSG00000C | TRUE | ir^1\|ale^13\| | TRUE | Distal | chr1:168961 | FALSE | TRUE | TRUE | 0.001 | 0.804 | 0.797 | 1 | 0 | AFE/ALE | 0.797 |
| 7 | ENSG00000C | ENSG00000C | TNRC6A | chr16 | + | ENSG00000C | ENSG00000C | TRUE | cassette^2\|t | FALSE | A_C2 | chr16:24758 | FALSE | FALSE | FALSE | 0.035 | 0.14 | 0.105 | 0.905 | 0.04 | cassette | 0.105 |
| 8 | ENSG000001 | ENSG000001 | BBC3 | chr19 | - | ENSG000001 | ENSG000001 | FALSE | afe^1 | TRUE | Distal | chr19:47226 | FALSE | FALSE | FALSE | 0.047 | 0.203 | 0.149 | 0.924 | 0.016 | AFE/ALE | 0.149 |
| 9 | ENSG000001 | ENSG000001 | CBL | chr11 | + | ENSG000001 | ENSG000001 | FALSE | cassette^1 | TRUE | C2_C1 | chr11:11920 | FALSE | FALSE | FALSE | 0.028 | 0.17 | -0.132 | 0.943 | 0.018 | skipping | 0.132 |
| 10 | ENSG000001 | ENSG000001 | ERI3 | chr1 | - | ENSG000001 | ENSG000001 | TRUE | cassette^3\|t | FALSE | C1_A | chr1:443529 | FALSE | FALSE | FALSE | 0.032 | 0.138 | 0.104 | 0.926 | 0.033 | cassette | 0.104 |
| 11 | ENSG000001 | ENSG000001 | GRIA3 | chrX | + | ENSG000001 | ENSG000001 | FALSE | ale^1 | FALSE | Proximal | chrX:123185 | FALSE | FALSE | FALSE | 0.016 | 0.163 | 0.126 | 0.928 | 0.024 | AFE/ALE | 0.126 |
| 12 | ENSG000001 | ENSG000001 | RFX1 | chr19 | - | ENSG000001 | ENSG000001 | FALSE | cassette^1 | TRUE | C1_A | chr19:13979 | FALSE | FALSE | TRUE | 0.041 | 0.261 | 0.215 | 0.996 | 0.001 | cassette | 0.215 |
| 13 | ENSG000001 | ENSG000001 | DAGLA | chr11 | + | ENSG000001 | ENSG000001 | FALSE | putative_ale | TRUE | Proximal | chr11:61726 | FALSE | FALSE | FALSE | 0.041 | 0.178 | 0.135 | 0.913 | 0.023 | AFE/ALE | 0.135 |
| 14 | ENSG000001 | ENSG000001 | RHBDL3 | chr17 | + | ENSG000001 | ENSG000001 | TRUE | cassette^1\|i | FALSE | C2_C1 | chr17:32266 | FALSE | FALSE | FALSE | 0.045 | 0.229 | -0.16 | 0.913 | 0.013 | skipping | 0.16 |
| 15 | ENSG000001 | ENSG000001 | FAM171B | chr2 | + | ENSG000001 | ENSG000001 | FALSE | putative_ale | TRUE | Proximal | chr2:186694 | FALSE | FALSE | FALSE | 0.026 | 0.121 | 0.094 | 0.901 | 0.048 | AFE/ALE | 0.094 |
| 16 | ENSG000001 | ENSG000001 | RAPGEF6 | chr5 | - | ENSG000001 | ENSG000001 | TRUE | putative_alt3 | FALSE | Proximal | chr5:131436 | FALSE | FALSE | TRUE | 0.008 | 0.28 | 0.255 | 1 | 0 | AFE/ALE | 0.255 |
| 17 | ENSG000001 | ENSG000001 | SPON2 | chr4 | - | ENSG000001 | ENSG000001 | TRUE | afe^1\|putati | TRUE | Distal | chr4:117230 | FALSE | FALSE | TRUE | 0.028 | 0.219 | 0.174 | 0.961 | 0.006 | AFE/ALE | 0.174 |
| 18 | ENSG000001 | ENSG000001 | TEX264 | chr3 | + | ENSG000001 | ENSG000001 | TRUE | cassette^3\|a | FALSE | C1_C2 | chr3:516745 | FALSE | FALSE | TRUE | 0.039 | 0.21 | -0.164 | 0.967 | 0.008 | skipping | 0.164 |
| 19 | ENSG000001 | ENSG000001 | PPP2R3B | chrX | - | ENSG000001 | ENSG000001 | TRUE | cassette^1\|a | FALSE | C2_A | chrX:361590 | FALSE | FALSE | TRUE | 0.031 | 0.179 | 0.147 | 0.974 | 0.008 | cassette | 0.147 |
| 20 | ENSG000001 | ENSG000001 | SPNS1 | chr16 | + | ENSG000001 | ENSG000001 | TRUE | cassette^2\|a | TRUE | Proximal | chr16:28978 | FALSE | TRUE | TRUE | 0.05 | 0.383 | 0.329 | 1 | 0 | AFE/ALE | 0.329 |
| 21 | ENSG000001 | ENSG000001 | ZNF417 | chr19 | - | ENSG000001 | ENSG000001 | TRUE | cassette^2\|a | FALSE | C1_A | chr19:57915 | FALSE | FALSE | TRUE | 0.012 | 0.192 | 0.156 | 0.988 | 0.003 | cassette | 0.156 |
| 22 | ENSG000001 | ENSG000001 | PITPNA | chr17 | - | ENSG000001 | ENSG000001 | FALSE | alt5^1 | TRUE | Proximal | chr17:15342 | FALSE | FALSE | TRUE | 0.035 | 0.144 | 0.119 | 0.991 | 0.004 | AFE/ALE | 0.119 |
| 23 | ENSG000001 | ENSG000001 | ZNF25 | chr10 | - | ENSG000001 | ENSG000001 | TRUE | putative_afe | TRUE | Proximal | chr10:37971 | FALSE | FALSE | FALSE | 0.042 | 0.208 | 0.15 | 0.913 | 0.017 | AFE/ALE | 0.15 |
| 24 | ENSG000001 | ENSG000001 | GUSBP1 | chr5 | + | ENSG000001 | ENSG000001 | TRUE | cassette^1\|i | FALSE | A_C2 | chr5:214915 | FALSE | FALSE | FALSE | 0.017 | 0.156 | 0.121 | 0.916 | 0.031 | cassette | 0.121 |
| 25 | ENSG000001 | ENSG000001 | WASH3P | chr15 | + | ENSG000001 | ENSG000001 | TRUE | cassette^3\|a | FALSE | Proximal | chr15:10197 | FALSE | FALSE | TRUE | 0.022 | 0.194 | 0.168 | 1 | 0 | AFE/ALE | 0.168 |
| 26 | ENSG000001 | ENSG000001 | ZNF136 | chr19 | + | ENSG000001 | ENSG000001 | FALSE | cassette^1 | FALSE | C1_C2 | chr19:12163 | FALSE | FALSE | TRUE | 0.033 | 0.19 | -0.149 | 0.967 | 0.009 | skipping | 0.149 |
| 27 | ENSG000001 | ENSG000001 | SGTB | chr5 | - | ENSG000001 | ENSG000001 | FALSE | cassette^1 | TRUE | C1_A | chr5:657047 | FALSE | FALSE | TRUE | 0.027 | 0.161 | 0.133 | 0.954 | 0.015 | cassette | 0.133 |
| 28 | ENSG000002 | ENSG000002 | PCDHA2 | chr5 | + | ENSG000002 | ENSG000002 | TRUE | ale^2\|putati | TRUE | Proximal | chr5:140795 | FALSE | FALSE | TRUE | 0.001 | 0.311 | 0.286 | 1 | 0 | AFE/ALE | 0.286 |
| 29 | ENSG000002 | ENSG000002 | PCDHA2 | chr5 | + | ENSG000002 | ENSG000002 | TRUE | ale^2\|putative_ale^8\|orp | | C1_A | chr5: | FALSE | FALSE | TRUE | 0.019 | 0.163 | 0.13 | 0.969 | 0.011 | cassette | 0.13 |
| 30 | ENSG000002 | ENSG000002 | PCDHA1 | chr5 | + | ENSG000002 | ENSG000002 | TRUE | ale^2\|putative_ale^2\|orp | | C1_A | chr5: | FALSE | FALSE | TRUE | 0.019 | 0.163 | 0.13 | 0.969 | 0.011 | cassette | 0.13 |
| 31 | ENSG000002 | ENSG000002 | ANKRD28 | chr3 | - | ENSG000002 | ENSG000002 | TRUE | cassette^2\|a | TRUE | C1_A | chr3:157125 | FALSE | FALSE | TRUE | 0 | 0.212 | 0.179 | 1 | 0 | cassette | 0.179 |
| 32 | ENSG000002 | ENSG000002 | ANKRD28 | chr3 | - | ENSG000002 | ENSG000002 | TRUE | cassette^2\|a | TRUE | C1_A | chr3:157125 | FALSE | FALSE | TRUE | 0 | 0.28 | 0.257 | 0.993 | 0.001 | cassette | 0.257 |
| 33 | ENSG000002 | ENSG000002 | ANKRD28 | chr3 | - | ENSG000002 | ENSG000002 | TRUE | cassette^2\|a | FALSE | A_C2 | chr3:156783 | FALSE | FALSE | TRUE | 0.04 | 0.339 | 0.313 | 0.981 | 0.001 | cassette | 0.313 |
| 34 | ENSG000002 | ENSG000002 | ANKRD28 | chr3 | - | ENSG000002 | ENSG000002 | TRUE | cassette^2\|alt3^38\|alt5^ | | A_C2 | chr3: | FALSE | FALSE | FALSE | 0.001 | 0.188 | 0.15 | 0.921 | 0.021 | cassette | 0.15 |
| 35 | ENSG000002 | ENSG000002 | ANKRD28 | chr3 | - | ENSG000002 | ENSG000002 | TRUE | cassette^2\|a | TRUE | E2_E1_J2 | chr3:156957 | FALSE | FALSE | TRUE | 0 | 0.186 | 0.152 | 0.962 | 0.011 | cassette | 0.152 |
| 36 | ENSG000002 | ENSG000002 | ANKRD28 | chr3 | - | ENSG000002 | ENSG000002 | TRUE | cassette^2\|a | TRUE | Distal | chr3:156962 | FALSE | FALSE | TRUE | 0 | 0.401 | 0.38 | 1 | 0 | AFE/ALE | 0.38 |
| 37 | ENSG000002 | ENSG000002 | ANKRD28 | chr3 | - | ENSG000002 | ENSG000002 | TRUE | cassette^2\|a | TRUE | E2_E1_J2 | chr3:157113 | FALSE | FALSE | TRUE | 0.013 | 0.167 | 0.135 | 0.951 | 0.015 | cassette | 0.135 |
| 38 | ENSG000002 | ENSG000002 | ANKRD28 | chr3 | - | ENSG000002 | ENSG000002 | TRUE | cassette^2\|alt3^38\|alt5^ | | A_C2 | chr3: | FALSE | FALSE | TRUE | 0 | 0.158 | 0.133 | 0.979 | 0.008 | cassette | 0.133 |
| 39 | ENSG000002 | ENSG000002 | ANKRD28 | chr3 | - | ENSG000002 | ENSG000002 | TRUE | cassette^2\|a | TRUE | E2_E1_J1 | chr3:157161 | FALSE | FALSE | TRUE | 0.007 | 0.154 | 0.128 | 0.964 | 0.016 | cassette | 0.128 |
| 40 | ENSG000002 | ENSG000002 | ANKRD28 | chr3 | - | ENSG000002 | ENSG000002 | TRUE | cassette^2\|a | TRUE | Distal | chr3:157069 | FALSE | TRUE | TRUE | 0.001 | 0.506 | 0.497 | 1 | 0 | AFE/ALE | 0.497 |
| 41 | ENSG000002 | ENSG000002 | ANKRD28 | chr3 | - | ENSG000002 | ENSG000002 | TRUE | cassette^2\|a | TRUE | Distal | chr3:157120 | FALSE | FALSE | TRUE | 0.002 | 0.441 | 0.429 | 0.998 | 0 | AFE/ALE | 0.429 |
| 42 | ENSG000002 | ENSG000002 | RPS29 | chr14 | - | ENSG000002 | ENSG000002 | TRUE | alt3^26\|alt5 | TRUE | E2_E1_J2 | chr14:49575 | FALSE | FALSE | TRUE | 0.031 | 0.127 | 0.101 | 0.978 | 0.013 | cassette | 0.101 |
| 43 | ENSG000002 | ENSG000002 | RPS29 | chr14 | - | ENSG000002 | ENSG000002 | TRUE | alt3^26\|alt5 | TRUE | E2_E1_J2 | chr14:49586 | FALSE | FALSE | FALSE | 0 | 0.13 | 0.104 | 0.911 | 0.043 | cassette | 0.104 |
| 44 | ENSG000002 | ENSG000002 | SNHG14 | chr15 | + | ENSG000002 | ENSG000002 | TRUE | cassette^13\|alt3^3\|alt5^ | | C1_A | chr15: | FALSE | FALSE | TRUE | 0.007 | 0.529 | 0.522 | 1 | 0 | cassette | 0.522 |
| 45 | ENSG000002 | ENSG000002 | SNHG14 | chr15 | + | ENSG000002 | ENSG000002 | TRUE | cassette^3\|a | TRUE | E1_E2_J2 | chr15:25193 | FALSE | FALSE | TRUE | 0.002 | 0.484 | 0.466 | 1 | 0 | cassette | 0.466 |
| 46 | ENSG000002 | ENSG000002 | SNHG14 | chr15 | + | ENSG000002 | ENSG000002 | TRUE | cassette^1\|a | FALSE | C1_A | chr15:25230 | FALSE | TRUE | TRUE | 0.021 | 0.713 | 0.693 | 1 | 0 | cassette | 0.693 |
| 47 | ENSG000002 | ENSG000002 | SNHG14 | chr15 | + | ENSG000002 | ENSG000002 | TRUE | cassette^13 | TRUE | E2_E1_J1 | chr15:25056 | FALSE | FALSE | TRUE | 0.009 | 0.224 | 0.197 | 0.998 | 0 | cassette | 0.197 |
| 48 | ENSG000002 | ENSG000002 | SNHG14 | chr15 | + | ENSG000002 | ENSG000002 | TRUE | cassette^13\|alt3^3\|alt5^ | | A_C2 | chr15: | FALSE | FALSE | TRUE | 0.006 | 0.215 | 0.201 | 0.968 | 0.004 | cassette | 0.201 |
| 49 | ENSG000002 | ENSG000002 | SNHG14 | chr15 | + | ENSG000002 | ENSG000002 | TRUE | cassette^3\|a | TRUE | Distal | chr15:25193 | FALSE | TRUE | TRUE | 0.002 | 0.527 | 0.506 | 1 | 0 | AFE/ALE | 0.506 |
| 50 | ENSG000002 | ENSG000002 | SNHG14 | chr15 | + | ENSG000002 | ENSG000002 | TRUE | cassette^1\|a | TRUE | Distal | chr15:25230 | FALSE | TRUE | TRUE | 0.021 | 0.715 | 0.705 | 1 | 0 | AFE/ALE | 0.705 |
| 51 | ENSG000002 | ENSG000002 | PARG | chr10 | - | ENSG000002 | ENSG000002 | FALSE | cassette^1 | FALSE | C2_C1 | chr10:49922 | FALSE | FALSE | FALSE | 0.033 | 0.2 | -0.145 | 0.913 | 0.017 | skipping | 0.145 |
| 52 | ENSG000002 | ENSG000002 | ENSG000002 | chr21 | + | ENSG000002 | ENSG000002 | TRUE | cassette^42\|alt3^33\|alt5 | | C1_A | chr21: | FALSE | TRUE | TRUE | 0.003 | 0.273 | 0.251 | 1 | 0 | cassette | 0.251 |
| 53 | ENSG000002 | ENSG000002 | ENSG000002 | chr21 | + | ENSG000002 | ENSG000002 | TRUE | cassette^42\|alt3^33\|alt5 | | C1_A | chr21: | FALSE | FALSE | FALSE | 0.016 | 0.154 | 0.123 | 0.925 | 0.027 | cassette | 0.123 |
| 54 | ENSG000002 | ENSG000002 | ENSG000002 | chr21 | + | ENSG000002 | ENSG000002 | TRUE | cassette^42\|alt3^33\|alt5 | | C1_A | chr21: | FALSE | FALSE | TRUE | 0.001 | 0.226 | 0.196 | 0.996 | 0.001 | cassette | 0.196 |
| 55 | ENSG000002 | ENSG000002 | ENSG000002 | chr21 | + | ENSG000002 | ENSG000002 | TRUE | cassette^42\|alt3^33\|alt5 | | C1_A | chr21: | FALSE | TRUE | TRUE | 0.03 | 0.199 | 0.171 | 1 | 0 | cassette | 0.171 |
| 56 | ENSG000002 | ENSG000002 | ENSG000002 | chr21 | + | ENSG000002 | ENSG000002 | TRUE | cassette^42\|alt3^33\|alt5 | | C1_A | chr21: | FALSE | FALSE | TRUE | 0 | 0.203 | 0.184 | 0.998 | 0 | cassette | 0.184 |
| 57 | ENSG000002 | ENSG000002 | ENSG000002 | chr21 | + | ENSG000002 | ENSG000002 | TRUE | cassette^42 | TRUE | Distal | chr21:82151 | FALSE | FALSE | TRUE | 0 | 0.397 | 0.371 | 1 | 0 | AFE/ALE | 0.371 |
| 58 | ENSG000002 | ENSG000002 | ENSG000002 | chr21 | + | ENSG000002 | ENSG000002 | TRUE | cassette^42 | TRUE | E1_E2_J1 | chr21:82171 | FALSE | FALSE | TRUE | 0.033 | 0.174 | 0.137 | 0.979 | 0.007 | cassette | 0.137 |
| 59 | ENSG000002 | ENSG000002 | ENSG000002 | chr21 | + | ENSG000002 | ENSG000002 | TRUE | cassette^42 | TRUE | C1_A | chr21:81990 | FALSE | FALSE | TRUE | 0.001 | 0.208 | 0.175 | 0.998 | 0.001 | cassette | 0.175 |
| 60 | ENSG000002 | ENSG000002 | ENSG000002 | chr21 | + | ENSG000002 | ENSG000002 | TRUE | cassette^42 | TRUE | Distal | chr21:82059 | FALSE | FALSE | TRUE | 0.001 | 0.335 | 0.294 | 1 | 0 | AFE/ALE | 0.294 |
| 61 | ENSG000002 | ENSG000002 | ENSG000002 | chr21 | + | ENSG000002 | ENSG000002 | TRUE | cassette^42 | TRUE | Distal | chr21:82005 | FALSE | FALSE | TRUE | 0.039 | 0.229 | 0.184 | 0.953 | 0.006 | AFE/ALE | 0.184 |
| 62 | ENSG000002 | ENSG000002 | ENSG000002 | chr21 | + | ENSG000002 | ENSG000002 | TRUE | cassette^42 | TRUE | C2_C1 | chr21:82059 | FALSE | FALSE | TRUE | 0.002 | 0.241 | -0.21 | 0.99 | 0.002 | skipping | 0.21 |
| 63 | ENSG000002 | ENSG000002 | ENSG000002 | chr21 | + | ENSG000002 | ENSG000002 | TRUE | cassette^42 | TRUE | Proximal | chr21:82109 | FALSE | FALSE | TRUE | 0.004 | 0.324 | 0.303 | 1 | 0 | AFE/ALE | 0.303 |
| 64 | ENSG000002 | ENSG000002 | ENSG000002 | chr21 | + | ENSG000002 | ENSG000002 | TRUE | cassette^42\|alt3^33\|alt5 | | A_C2 | chr21: | FALSE | FALSE | TRUE | 0.012 | 0.351 | 0.317 | 1 | 0 | cassette | 0.317 |
| 65 | ENSG000002 | ENSG000002 | ENSG000002 | chr21 | + | ENSG000002 | ENSG000002 | TRUE | cassette^42 | TRUE | C2_C1 | chr21:82072 | FALSE | FALSE | FALSE | 0.04 | 0.249 | -0.17 | 0.927 | 0.011 | skipping | 0.17 |
| 66 | ENSG000002 | ENSG000002 | ENSG000002 | chr21 | + | ENSG000002 | ENSG000002 | TRUE | cassette^42 | TRUE | J12 | chr21:82181 | FALSE | FALSE | TRUE | 0.027 | 0.303 | 0.276 | 0.999 | 0 | cassette | 0.276 |
| 67 | ENSG000002 | ENSG000002 | ENSG000002 | chr21 | + | ENSG000002 | ENSG000002 | TRUE | cassette^42 | TRUE | C2_A | chr21:82120 | FALSE | FALSE | TRUE | 0.002 | 0.188 | 0.17 | 0.971 | 0.007 | cassette | 0.17 |
| 68 | ENSG000002 | ENSG000002 | ENSG000002 | chr21 | + | ENSG000002 | ENSG000002 | TRUE | cassette^42 | TRUE | Proximal | chr21:82151 | FALSE | TRUE | TRUE | 0.002 | 0.685 | 0.683 | 1 | 0 | AFE/ALE | 0.683 |
| 69 | ENSG000002 | ENSG000002 | ENSG000002 | chr21 | + | ENSG000002 | ENSG000002 | TRUE | cassette^42 | TRUE | Distal | chr21:82063 | FALSE | FALSE | TRUE | 0.007 | 0.923 | 0.913 | 1 | 0 | AFE/ALE | 0.913 |
| 70 | ENSG000002 | ENSG000002 | ENSG000002 | chr21 | + | ENSG000002 | ENSG000002 | TRUE | cassette^126\|alt3^121\|a | | C1_A | chr21: | FALSE | FALSE | TRUE | 0.009 | 0.294 | 0.265 | 1 | 0 | cassette | 0.265 |
| 71 | ENSG000002 | ENSG000002 | ENSG000002 | chr21 | + | ENSG000002 | ENSG000002 | TRUE | cassette^126\|alt3^121\|a | | C1_A | chr21: | FALSE | FALSE | TRUE | 0 | 0.175 | 0.151 | 0.989 | 0.003 | cassette | 0.151 |

| | | | | | | | | | | | | | | | | | | | | | | |
|---|---|---|---|---|---|---|---|---|---|---|---|---|---|---|---|---|---|---|---|---|---|---|
| 72 | ENSG000002 | ENSG000002 | ENSG000002 | chr21 | + | ENSG000002 | ENSG000002 | TRUE | cassette^126\|alt3^121\|a | | C1_A | chr21: | FALSE | FALSE | TRUE | 0.004 | 0.341 | 0.307 | 0.996 | 0 | cassette | 0.307 |
| 73 | ENSG000002 | ENSG000002 | ENSG000002 | chr21 | + | ENSG000002 | ENSG000002 | TRUE | cassette^126 | TRUE | C1_A | chr21:83933 | FALSE | FALSE | TRUE | 0.006 | 0.268 | 0.252 | 1 | 0 | cassette | 0.252 |
| 74 | ENSG000002 | ENSG000002 | ENSG000002 | chr21 | + | ENSG000002 | ENSG000002 | TRUE | cassette^126\|alt3^121\|a | | C1_A | chr21: | FALSE | FALSE | TRUE | 0.019 | 0.146 | 0.124 | 0.95 | 0.018 | cassette | 0.124 |
| 75 | ENSG000002 | ENSG000002 | ENSG000002 | chr21 | + | ENSG000002 | ENSG000002 | TRUE | cassette^126\|alt3^121\|a | | C1_A | chr21: | FALSE | TRUE | TRUE | 0.032 | 0.253 | 0.214 | 1 | 0 | cassette | 0.214 |
| 76 | ENSG000002 | ENSG000002 | ENSG000002 | chr21 | + | ENSG000002 | ENSG000002 | TRUE | cassette^126\|alt3^121\|a | | C1_A | chr21: | FALSE | FALSE | TRUE | 0.02 | 0.173 | 0.147 | 0.974 | 0.008 | cassette | 0.147 |
| 77 | ENSG000002 | ENSG000002 | ENSG000002 | chr21 | + | ENSG000002 | ENSG000002 | TRUE | cassette^126\|alt3^121\|a | | C1_A | chr21: | FALSE | FALSE | TRUE | 0.028 | 0.255 | 0.219 | 1 | 0 | cassette | 0.219 |
| 78 | ENSG000002 | ENSG000002 | ENSG000002 | chr21 | + | ENSG000002 | ENSG000002 | TRUE | cassette^126 | TRUE | E1_E2_J1 | chr21:83986 | FALSE | FALSE | TRUE | 0 | 0.186 | 0.16 | 1 | 0 | cassette | 0.16 |
| 79 | ENSG000002 | ENSG000002 | ENSG000002 | chr21 | + | ENSG000002 | ENSG000002 | TRUE | cassette^126\|alt3^121\|a | | C1_A | chr21: | FALSE | FALSE | TRUE | 0.013 | 0.157 | 0.132 | 0.955 | 0.016 | cassette | 0.132 |
| 80 | ENSG000002 | ENSG000002 | ENSG000002 | chr21 | + | ENSG000002 | ENSG000002 | TRUE | cassette^126\|alt3^121\|a | | C1_A | chr21: | FALSE | FALSE | TRUE | 0 | 0.147 | 0.123 | 0.951 | 0.02 | cassette | 0.123 |
| 81 | ENSG000002 | ENSG000002 | ENSG000002 | chr21 | + | ENSG000002 | ENSG000002 | TRUE | cassette^126\|alt3^121\|a | | C1_A | chr21: | FALSE | FALSE | TRUE | 0 | 0.265 | 0.244 | 1 | 0 | cassette | 0.244 |
| 82 | ENSG000002 | ENSG000002 | ENSG000002 | chr21 | + | ENSG000002 | ENSG000002 | TRUE | cassette^126\|alt3^121\|a | | C1_A | chr21: | FALSE | FALSE | FALSE | 0 | 0.119 | 0.096 | 0.913 | 0.046 | cassette | 0.096 |
| 83 | ENSG000002 | ENSG000002 | ENSG000002 | chr21 | + | ENSG000002 | ENSG000002 | TRUE | cassette^126 | TRUE | E1_E2_J2 | chr21:83998 | FALSE | FALSE | FALSE | 0.009 | 0.148 | 0.121 | 0.932 | 0.024 | cassette | 0.121 |
| 84 | ENSG000002 | ENSG000002 | ENSG000002 | chr21 | + | ENSG000002 | ENSG000002 | TRUE | cassette^126\|alt3^121\|a | | C1_A | chr21: | FALSE | FALSE | TRUE | 0.025 | 0.325 | 0.304 | 1 | 0 | cassette | 0.304 |
| 85 | ENSG000002 | ENSG000002 | ENSG000002 | chr21 | + | ENSG000002 | ENSG000002 | TRUE | cassette^126\|alt3^121\|a | | C1_A | chr21: | FALSE | FALSE | FALSE | 0.016 | 0.227 | 0.192 | 0.932 | 0.01 | cassette | 0.192 |
| 86 | ENSG000002 | ENSG000002 | ENSG000002 | chr21 | + | ENSG000002 | ENSG000002 | TRUE | cassette^126\|alt3^121\|a | | C1_A | chr21: | FALSE | FALSE | FALSE | 0.015 | 0.131 | 0.114 | 0.927 | 0.032 | cassette | 0.114 |
| 87 | ENSG000002 | ENSG000002 | ENSG000002 | chr21 | + | ENSG000002 | ENSG000002 | TRUE | cassette^126\|alt3^121\|a | | C1_A | chr21: | FALSE | FALSE | TRUE | 0.034 | 0.19 | 0.163 | 0.982 | 0.004 | cassette | 0.163 |
| 88 | ENSG000002 | ENSG000002 | ENSG000002 | chr21 | + | ENSG000002 | ENSG000002 | TRUE | cassette^126\|alt3^121\|a | | C1_A | chr21: | FALSE | FALSE | TRUE | 0.004 | 0.412 | 0.39 | 1 | 0 | cassette | 0.39 |
| 89 | ENSG000002 | ENSG000002 | ENSG000002 | chr21 | + | ENSG000002 | ENSG000002 | TRUE | cassette^126\|alt3^121\|a | | C1_A | chr21: | FALSE | FALSE | TRUE | 0 | 0.212 | 0.186 | 0.997 | 0.001 | cassette | 0.186 |
| 90 | ENSG000002 | ENSG000002 | ENSG000002 | chr21 | + | ENSG000002 | ENSG000002 | TRUE | cassette^126\|alt3^121\|a | | C1_A | chr21: | FALSE | FALSE | TRUE | 0.004 | 0.379 | 0.346 | 1 | 0 | cassette | 0.346 |
| 91 | ENSG000002 | ENSG000002 | ENSG000002 | chr21 | + | ENSG000002 | ENSG000002 | TRUE | cassette^126\|alt3^121\|a | | C1_A | chr21: | FALSE | FALSE | TRUE | 0.001 | 0.182 | 0.173 | 0.964 | 0.008 | cassette | 0.173 |
| 92 | ENSG000002 | ENSG000002 | ENSG000002 | chr21 | + | ENSG000002 | ENSG000002 | TRUE | cassette^126\|alt3^121\|a | | C1_A | chr21: | FALSE | FALSE | TRUE | 0.002 | 0.248 | 0.216 | 1 | 0 | cassette | 0.216 |
| 93 | ENSG000002 | ENSG000002 | ENSG000002 | chr21 | + | ENSG000002 | ENSG000002 | TRUE | cassette^126\|alt3^121\|a | | C1_A | chr21: | FALSE | FALSE | TRUE | 0 | 0.167 | 0.147 | 0.974 | 0.008 | cassette | 0.147 |
| 94 | ENSG000002 | ENSG000002 | ENSG000002 | chr21 | + | ENSG000002 | ENSG000002 | TRUE | cassette^126\|alt3^121\|a | | C1_A | chr21: | FALSE | FALSE | TRUE | 0 | 0.199 | 0.171 | 1 | 0 | cassette | 0.171 |
| 95 | ENSG000002 | ENSG000002 | ENSG000002 | chr21 | + | ENSG000002 | ENSG000002 | TRUE | cassette^126\|alt3^121\|a | | C1_A | chr21: | FALSE | TRUE | TRUE | 0 | 0.422 | 0.399 | 1 | 0 | cassette | 0.399 |
| 96 | ENSG000002 | ENSG000002 | ENSG000002 | chr21 | + | ENSG000002 | ENSG000002 | TRUE | cassette^126\|alt3^121\|a | | C1_A | chr21: | FALSE | FALSE | TRUE | 0.001 | 0.21 | 0.172 | 0.981 | 0.005 | cassette | 0.172 |
| 97 | ENSG000002 | ENSG000002 | ENSG000002 | chr21 | + | ENSG000002 | ENSG000002 | TRUE | cassette^126\|alt3^121\|a | | C1_A | chr21: | FALSE | FALSE | TRUE | 0.001 | 0.196 | 0.164 | 0.978 | 0.005 | cassette | 0.164 |
| 98 | ENSG000002 | ENSG000002 | ENSG000002 | chr21 | + | ENSG000002 | ENSG000002 | TRUE | cassette^126\|alt3^121\|a | | C1_A | chr21: | FALSE | FALSE | TRUE | 0 | 0.214 | 0.185 | 0.981 | 0.004 | cassette | 0.185 |
| 99 | ENSG000002 | ENSG000002 | ENSG000002 | chr21 | + | ENSG000002 | ENSG000002 | TRUE | cassette^126\|alt3^121\|a | | C1_A | chr21: | FALSE | FALSE | TRUE | 0.001 | 0.22 | 0.192 | 0.999 | 0 | cassette | 0.192 |
| 100 | ENSG000002 | ENSG000002 | ENSG000002 | chr21 | + | ENSG000002 | ENSG000002 | TRUE | cassette^126\|alt3^121\|a | | C1_A | chr21: | FALSE | FALSE | TRUE | 0.017 | 0.23 | 0.198 | 0.992 | 0.001 | cassette | 0.198 |
| 101 | ENSG000002 | ENSG000002 | ENSG000002 | chr21 | + | ENSG000002 | ENSG000002 | TRUE | cassette^126\|alt3^121\|a | | C1_A | chr21: | FALSE | FALSE | FALSE | 0.001 | 0.137 | 0.116 | 0.913 | 0.037 | cassette | 0.116 |
| 102 | ENSG000002 | ENSG000002 | ENSG000002 | chr21 | + | ENSG000002 | ENSG000002 | TRUE | cassette^126\|alt3^121\|a | | C1_A | chr21: | FALSE | FALSE | TRUE | 0 | 0.539 | 0.517 | 1 | 0 | cassette | 0.517 |
| 103 | ENSG000002 | ENSG000002 | ENSG000002 | chr21 | + | ENSG000002 | ENSG000002 | TRUE | cassette^126 | TRUE | Distal | chr21:84387 | FALSE | FALSE | TRUE | 0.024 | 0.172 | 0.139 | 0.955 | 0.014 | AFE/ALE | 0.139 |
| 104 | ENSG000002 | ENSG000002 | ENSG000002 | chr21 | + | ENSG000002 | ENSG000002 | TRUE | cassette^126\|alt3^121\|a | | C1_A | chr21: | FALSE | FALSE | TRUE | 0.001 | 0.221 | 0.196 | 1 | 0 | cassette | 0.196 |
| 105 | ENSG000002 | ENSG000002 | ENSG000002 | chr21 | + | ENSG000002 | ENSG000002 | TRUE | cassette^126\|alt3^121\|a | | C1_A | chr21: | FALSE | FALSE | TRUE | 0.026 | 0.18 | 0.147 | 0.974 | 0.007 | cassette | 0.147 |
| 106 | ENSG000002 | ENSG000002 | ENSG000002 | chr21 | + | ENSG000002 | ENSG000002 | TRUE | cassette^126\|alt3^121\|a | | C1_A | chr21: | FALSE | FALSE | TRUE | 0.005 | 0.677 | 0.664 | 1 | 0 | cassette | 0.664 |
| 107 | ENSG000002 | ENSG000002 | ENSG000002 | chr21 | + | ENSG000002 | ENSG000002 | TRUE | cassette^126 | TRUE | Proximal | chr21:83900 | FALSE | FALSE | FALSE | 0.016 | 0.181 | 0.148 | 0.935 | 0.013 | AFE/ALE | 0.148 |
| 108 | ENSG000002 | ENSG000002 | ENSG000002 | chr21 | + | ENSG000002 | ENSG000002 | TRUE | cassette^126 | TRUE | Distal | chr21:84088 | FALSE | FALSE | TRUE | 0 | 0.146 | 0.121 | 0.978 | 0.009 | AFE/ALE | 0.121 |
| 109 | ENSG000002 | ENSG000002 | ENSG000002 | chr21 | + | ENSG000002 | ENSG000002 | TRUE | cassette^126 | TRUE | Distal | chr21:84244 | FALSE | FALSE | TRUE | 0.005 | 0.228 | 0.211 | 0.994 | 0.001 | AFE/ALE | 0.211 |
| 110 | ENSG000002 | ENSG000002 | ENSG000002 | chr21 | + | ENSG000002 | ENSG000002 | TRUE | cassette^126 | TRUE | Distal | chr21:83818 | FALSE | FALSE | TRUE | 0.001 | 0.173 | 0.153 | 0.966 | 0.01 | AFE/ALE | 0.153 |
| 111 | ENSG000002 | ENSG000002 | ENSG000002 | chr21 | + | ENSG000002 | ENSG000002 | TRUE | cassette^126\|alt3^121\|a | | A_C2 | chr21: | FALSE | FALSE | TRUE | 0.001 | 0.299 | 0.284 | 0.951 | 0.005 | cassette | 0.284 |
| 112 | ENSG000002 | ENSG000002 | ENSG000002 | chr21 | + | ENSG000002 | ENSG000002 | TRUE | cassette^126 | TRUE | E2_E1_J1 | chr21:83807 | FALSE | FALSE | TRUE | 0 | 0.24 | 0.208 | 0.969 | 0.005 | cassette | 0.208 |
| 113 | ENSG000002 | ENSG000002 | ENSG000002 | chr21 | + | ENSG000002 | ENSG000002 | TRUE | cassette^126 | TRUE | Proximal | chr21:83876 | FALSE | FALSE | FALSE | 0.034 | 0.208 | 0.168 | 0.939 | 0.009 | AFE/ALE | 0.168 |
| 114 | ENSG000002 | ENSG000002 | ENSG000002 | chr21 | + | ENSG000002 | ENSG000002 | TRUE | cassette^126 | TRUE | C2_C1 | chr21:83876 | FALSE | FALSE | FALSE | 0 | 0.126 | -0.111 | 0.921 | 0.036 | skipping | 0.111 |
| 115 | ENSG000002 | ENSG000002 | ENSG000002 | chr21 | + | ENSG000002 | ENSG000002 | TRUE | cassette^126\|alt3^121\|a | | A_C2 | chr21: | FALSE | FALSE | TRUE | 0.021 | 0.21 | 0.171 | 0.975 | 0.005 | cassette | 0.171 |
| 116 | ENSG000002 | ENSG000002 | ENSG000002 | chr21 | + | ENSG000002 | ENSG000002 | TRUE | cassette^126\|alt3^121\|a | | A_C2 | chr21: | FALSE | FALSE | TRUE | 0.001 | 0.273 | 0.256 | 1 | 0 | cassette | 0.256 |
| 117 | ENSG000002 | ENSG000002 | ENSG000002 | chr21 | + | ENSG000002 | ENSG000002 | TRUE | cassette^126\|alt3^121\|a | | A_C2 | chr21: | FALSE | FALSE | TRUE | 0.001 | 0.203 | 0.171 | 0.995 | 0.001 | cassette | 0.171 |
| 118 | ENSG000002 | ENSG000002 | ENSG000002 | chr21 | + | ENSG000002 | ENSG000002 | TRUE | cassette^126 | TRUE | Distal | chr21:83929 | FALSE | FALSE | FALSE | 0 | 0.134 | 0.114 | 0.944 | 0.025 | AFE/ALE | 0.114 |
| 119 | ENSG000002 | ENSG000002 | ENSG000002 | chr21 | + | ENSG000002 | ENSG000002 | TRUE | cassette^126 | TRUE | Distal | chr21:83876 | FALSE | FALSE | FALSE | 0.001 | 0.148 | 0.125 | 0.939 | 0.022 | AFE/ALE | 0.125 |
| 120 | ENSG000002 | ENSG000002 | ENSG000002 | chr21 | + | ENSG000002 | ENSG000002 | TRUE | cassette^126\|alt3^121\|a | | A_C2 | chr21: | FALSE | FALSE | TRUE | 0 | 0.203 | 0.187 | 1 | 0 | cassette | 0.187 |
| 121 | ENSG000002 | ENSG000002 | ENSG000002 | chr21 | + | ENSG000002 | ENSG000002 | TRUE | cassette^126\|alt3^121\|a | | A_C2 | chr21: | FALSE | FALSE | TRUE | 0 | 0.212 | 0.206 | 0.984 | 0.003 | cassette | 0.206 |
| 122 | ENSG000002 | ENSG000002 | ENSG000002 | chr21 | + | ENSG000002 | ENSG000002 | TRUE | cassette^126 | TRUE | Distal | chr21:83899 | FALSE | FALSE | TRUE | 0.019 | 0.195 | 0.167 | 0.987 | 0.003 | AFE/ALE | 0.167 |
| 123 | ENSG000002 | ENSG000002 | ENSG000002 | chr21 | + | ENSG000002 | ENSG000002 | TRUE | cassette^126\|alt3^121\|a | | A_C2 | chr21: | FALSE | FALSE | TRUE | 0.036 | 0.171 | 0.151 | 0.999 | 0 | cassette | 0.151 |
| 124 | ENSG000002 | ENSG000002 | ENSG000002 | chr21 | + | ENSG000002 | ENSG000002 | TRUE | cassette^126\|alt3^121\|a | | A_C2 | chr21: | FALSE | FALSE | TRUE | 0.001 | 0.287 | 0.26 | 1 | 0 | cassette | 0.26 |
| 125 | ENSG000002 | ENSG000002 | ENSG000002 | chr21 | + | ENSG000002 | ENSG000002 | TRUE | cassette^126\|alt3^121\|a | | A_C2 | chr21: | FALSE | FALSE | TRUE | 0.013 | 0.237 | 0.196 | 0.966 | 0.004 | cassette | 0.196 |
| 126 | ENSG000002 | ENSG000002 | ENSG000002 | chr21 | + | ENSG000002 | ENSG000002 | TRUE | cassette^126\|alt3^121\|a | | A_C2 | chr21: | FALSE | FALSE | FALSE | 0.003 | 0.157 | 0.128 | 0.932 | 0.025 | cassette | 0.128 |
| 127 | ENSG000002 | ENSG000002 | ENSG000002 | chr21 | + | ENSG000002 | ENSG000002 | TRUE | cassette^126\|alt3^121\|a | | A_C2 | chr21: | FALSE | FALSE | TRUE | 0.001 | 0.185 | 0.159 | 0.993 | 0.002 | cassette | 0.159 |
| 128 | ENSG000002 | ENSG000002 | ENSG000002 | chr21 | + | ENSG000002 | ENSG000002 | TRUE | cassette^126\|alt3^121\|a | | A_C2 | chr21: | FALSE | FALSE | TRUE | 0.024 | 0.279 | 0.259 | 1 | 0 | cassette | 0.259 |
| 129 | ENSG000002 | ENSG000002 | ENSG000002 | chr21 | + | ENSG000002 | ENSG000002 | TRUE | cassette^126\|alt3^121\|a | | A_C2 | chr21: | FALSE | FALSE | TRUE | 0.002 | 0.252 | 0.224 | 1 | 0 | cassette | 0.224 |
| 130 | ENSG000002 | ENSG000002 | ENSG000002 | chr21 | + | ENSG000002 | ENSG000002 | TRUE | cassette^126\|alt3^121\|a | | A_C2 | chr21: | FALSE | FALSE | FALSE | 0 | 0.137 | 0.123 | 0.933 | 0.018 | cassette | 0.123 |
| 131 | ENSG000002 | ENSG000002 | ENSG000002 | chr21 | + | ENSG000002 | ENSG000002 | TRUE | cassette^126\|alt3^121\|a | | A_C2 | chr21: | FALSE | FALSE | FALSE | 0.027 | 0.178 | 0.135 | 0.913 | 0.021 | cassette | 0.135 |
| 132 | ENSG000002 | ENSG000002 | ENSG000002 | chr21 | + | ENSG000002 | ENSG000002 | TRUE | cassette^126\|alt3^121\|a | | A_C2 | chr21: | FALSE | FALSE | TRUE | 0.034 | 0.665 | 0.634 | 1 | 0 | cassette | 0.634 |
| 133 | ENSG000002 | ENSG000002 | ENSG000002 | chr21 | + | ENSG000002 | ENSG000002 | TRUE | cassette^126\|alt3^121\|a | | A_C2 | chr21: | FALSE | FALSE | TRUE | 0.003 | 0.184 | 0.157 | 0.994 | 0.002 | cassette | 0.157 |
| 134 | ENSG000002 | ENSG000002 | ENSG000002 | chr21 | + | ENSG000002 | ENSG000002 | TRUE | cassette^126\|alt3^121\|a | | A_C2 | chr21: | FALSE | FALSE | TRUE | 0.038 | 0.278 | 0.253 | 0.994 | 0.001 | cassette | 0.253 |
| 135 | ENSG000002 | ENSG000002 | ENSG000002 | chr21 | + | ENSG000002 | ENSG000002 | TRUE | cassette^126\|alt3^121\|a | | A_C2 | chr21: | FALSE | FALSE | TRUE | 0.013 | 0.148 | 0.123 | 1 | 0 | cassette | 0.123 |
| 136 | ENSG000002 | ENSG000002 | ENSG000002 | chr21 | + | ENSG000002 | ENSG000002 | TRUE | cassette^126\|alt3^121\|a | | A_C2 | chr21: | FALSE | FALSE | TRUE | 0.041 | 0.305 | 0.262 | 1 | 0 | cassette | 0.262 |
| 137 | ENSG000002 | ENSG000002 | ENSG000002 | chr21 | + | ENSG000002 | ENSG000002 | TRUE | cassette^126\|alt3^121\|a | | A_C2 | chr21: | FALSE | FALSE | TRUE | 0.048 | 0.519 | 0.395 | 0.991 | 0 | cassette | 0.395 |
| 138 | ENSG000002 | ENSG000002 | ENSG000002 | chr21 | + | ENSG000002 | ENSG000002 | TRUE | cassette^126 | TRUE | C2_A | chr21:84421 | FALSE | FALSE | TRUE | 0.033 | 0.185 | 0.146 | 0.97 | 0.009 | cassette | 0.146 |
| 139 | ENSG000002 | ENSG000002 | ENSG000002 | chr21 | + | ENSG000002 | ENSG000002 | TRUE | cassette^126\|alt3^121\|a | | A_C2 | chr21: | FALSE | FALSE | TRUE | 0.029 | 0.127 | 0.102 | 0.972 | 0.014 | cassette | 0.102 |
| 140 | ENSG000002 | ENSG000002 | ENSG000002 | chr21 | + | ENSG000002 | ENSG000002 | TRUE | cassette^126\|alt3^121\|a | | A_C2 | chr21: | FALSE | FALSE | TRUE | 0 | 0.169 | 0.147 | 1 | 0 | cassette | 0.147 |
| 141 | ENSG000002 | ENSG000002 | ENSG000002 | chr21 | + | ENSG000002 | ENSG000002 | TRUE | cassette^126 | TRUE | Proximal | chr21:84331 | FALSE | FALSE | TRUE | 0 | 0.393 | 0.366 | 1 | 0 | AFE/ALE | 0.366 |
| 142 | ENSG000002 | ENSG000002 | ENSG000002 | chr21 | + | ENSG000002 | ENSG000002 | TRUE | cassette^126\|alt3^121\|a | | A_C2 | chr21: | FALSE | FALSE | FALSE | 0.002 | 0.155 | 0.124 | 0.936 | 0.025 | cassette | 0.124 |
| 143 | ENSG000002 | ENSG000002 | ENSG000002 | chr21 | + | ENSG000002 | ENSG000002 | TRUE | cassette^126\|alt3^121\|a | | A_C2 | chr21: | FALSE | FALSE | TRUE | 0 | 0.213 | 0.187 | 0.971 | 0.005 | cassette | 0.187 |

| | | | | | | | | | | | | | | | | | | | | | | | | |
|---|---|---|---|---|---|---|---|---|---|---|---|---|---|---|---|---|---|---|---|---|---|---|---|---|
| 144 | ENSG000002 | ENSG000002 | ENSG000002 | chr21 | + | | ENSG000002 | ENSG000002 | TRUE | cassette^126\|alt3^121\|a | | A_C2 | chr21: | FALSE | FALSE | TRUE | 0.008 | 0.31 | 0.292 | 0.997 | 0 | cassette | | 0.292 |
| 145 | ENSG000002 | ENSG000002 | ENSG000002 | chr21 | + | | ENSG000002 | ENSG000002 | TRUE | cassette^126 | TRUE | C2_A1 | chr21:84027 | FALSE | FALSE | TRUE | 0.001 | 0.165 | 0.142 | 1 | 0 | cassette | | 0.142 |
| 146 | ENSG000002 | ENSG000002 | ENSG000002 | chr14 | - | | ENSG000002 | ENSG000002 | TRUE | alt3^5\|alt5^ | TRUE | E1_E2_J2 | chr14:49862 | FALSE | FALSE | TRUE | 0 | 0.164 | 0.138 | 0.981 | 0.007 | cassette | | 0.138 |
| 147 | ENSG000002 | ENSG000002 | ENSG000002 | chr3 | + | | ENSG000002 | ENSG000002 | TRUE | orphan_junc | TRUE | Orphan | chr3:134783 | FALSE | FALSE | FALSE | 0.046 | 0.186 | 0.139 | 0.944 | 0.016 | cassette | | 0.139 |
| 148 | ENSG000002 | ENSG000002 | ENSG000002 | chr3 | + | | ENSG000002 | ENSG000002 | TRUE | orphan_junc | TRUE | Orphan | chr3:134783 | FALSE | FALSE | TRUE | 0.014 | 0.528 | 0.514 | 1 | 0 | cassette | | 0.514 |
